# A druggable addiction to *de novo* pyrimidine biosynthesis in diffuse midline glioma

**DOI:** 10.1101/2021.11.30.470644

**Authors:** Sharmistha Pal, Jakub P. Kaplan, Huy Nguyen, Sylwia A. Stopka, Milan R. Savani, Michael S. Regan, Quang-De Nguyen, Kristen L. Jones, Lisa A. Moreau, Jingyu Peng, Marina G. Dipiazza, Andrew J. Perciaccante, Xiaoting Zhu, Bradley R. Hunsel, Kevin X. Liu, Rachid Drissi, Mariella G. Filbin, Samuel K. McBrayer, Nathalie Y.R. Agar, Dipanjan Chowdhury, Daphne Haas-Kogan

## Abstract

Diffuse midline glioma (DMG) is a uniformly fatal pediatric cancer driven by oncohistones that do not readily lend themselves to drug development. To identify druggable targets for DMG, we conducted a genome-wide CRISPR screen that reveals a DMG selective dependency on the *de novo* pathway for pyrimidine biosynthesis. This metabolic vulnerability reflects an elevated rate of uridine/uracil degradation that depletes DMG cells of substrates for the alternate salvage pathway for pyrimidine biosynthesis. A clinical stage inhibitor of DHODH (rate limiting enzyme in the *de novo* pathway) diminishes UMP pools, generates DNA damage, and induces apoptosis through suppression of replication forks--an “on target” effect, as shown by uridine rescue. MALDI mass spectroscopy imaging demonstrates that this DHODH inhibitor (BAY2402234) accumulates in brain at therapeutically relevant concentrations, suppresses *de novo* pyrimidine biosynthesis *in vivo*, and prolongs survival of mice bearing intracranial DMG xenografts, highlighting BAY2402234 as a promising therapy against DMGs.

## INTRODUCTION

Diffuse midline glioma (DMG), hitherto known as diffuse intrinsic pontine glioma (DIPG), is a high-grade glioma that develops in the midline structures of the brain. DMGs occur mostly in children (median age of 6-7 years), and less frequently in young adults. These malignant and universally fatal gliomas are not amenable to surgical resection due to the tumor site and their diffuse and infiltrative growth. Moreover, no chemotherapeutic, biological, or any other systemic agent has proven effective against DMGs. Radiotherapy prolongs survival of most patients, but tumor progression and death are inevitable. With median survival of less than a year, DMGs are the leading cause of brain cancer deaths in children (Braunstein et al., 2017; Cooney et al., 2017; Jones et al., 2017; Ostrom et al., 2017; Sturm et al., 2017).

Unlike most other pediatric cancers for which immense progress has been made in identifying efficacious novel treatments and improving patient outcomes, survival for DMGs has not changed in over 50 years. Given the failure of conventional therapeutic modalities, considerable effort has been devoted to development of targeted therapeutics. Seminal work over the last decade has identified the prevalent oncogenic drivers of DMG as a set of recurrent amino acid substitution mutations in histone H3.3 (*H3F3A*) and H3.1 (*HIST1H3B* and *HIST1H3C*) (Buczkowicz et al., 2014; Khuong-Quang et al., 2012; Wu et al., 2012). The H3K27M amino acid substitution is the most common (80%) leading to the designation, “H3K27M mutant diffuse midline glioma (DMG)” by the World Health Organization. The outcome of this single amino acid substitution is a genome-wide reconfiguration of chromatin architecture via loss of repressive H3K27 trimethylation (Bender et al., 2013; Harutyunyan et al., 2019; Wu et al., 2012). A subset of DMGs contain mutations in alternate effectors of chromatin architecture. In particular, EZHIP overexpression has been described as a second route towards loss of H3K27 trimethylation in a subset of those DMGs lacking H3K27M mutations (Castel et al., 2020).

Unfortunately, while H3K27M is a key driver in DMG, the mutated histone H3 has no enzymatic function and thus does not lend itself to development of small molecule antagonists (Bender et al., 2013; Krug et al., 2019; Larson et al., 2019; Nagaraja et al., 2019; Silveira et al., 2019). Various workarounds, including histone deacetylase (HDAC) inhibitors, are being developed but have yet to show any benefit for DMG patients. For example, the HDAC inhibitor panobinostat, identified in a drug screen, counteracts epigenetic dysregulation and restores H3K27 trimethylation but poses a challenge for clinical use due to its inability to cross the blood brain barrier (BBB) (Nagaraja et al., 2017). Besides epigenetic alterations, other common alterations in DMGs include inactivation of the TP53 pathway, activation of PI3K, PDGF or ACVR1 signaling, and dysregulation of G1-S cell cycle checkpoint (Mackay et al., 2017). Attempts to target these signaling pathways have yet to show any clinical benefit.

Against this backdrop, an emerging body of data shows that tumor cells adapt to the stresses created by oncogenic driver mutations (Luo et al., 2009; Pagliarini et al., 2015). These adaptations can create vulnerabilities to otherwise innocuous inhibitors of common metabolic pathways. Studies summarized herein explore this route towards a targeted therapeutic for DMG.

## RESULTS

### A genome-wide CRISPR loss-of-function screen for DMG vulnerabilities

To uncover intrinsic vulnerabilities of DMGs, we performed a genome-wide CRISPR-based dependency screen in three distinct DMG cell lines to identify genes whose loss would lead to DMG cell death (Figure S1A). As expected, many of the identified genes were known to be pan-cancer essential (Table S1) and these were excluded from further analysis. A Venn diagram depicting overlap among the remaining genes (henceforth referred to as DMG dependency genes, Table S2) in the three DMG lines revealed 213 genes shared between at least two cell lines but only 13 dependency genes common to all three DMG lines (Figure 1A). We performed Ingenuity pathway analysis (IPA) on the 213 genes shared by at least two of the cell lines and found that the top five biological pathways defined by these dependency genes (Figure 1B) are associated with metabolism and DNA damage repair. We chose to focus on uridine-5’-phosphate (UMP) biosynthesis because every component of the *de novo* arm of this pathway was among the DMG dependency genes (Figure 1C). Specifically, the three genes that execute *de novo* pyrimidine biosynthesis [carbamoyl-phosphate synthetase 2, aspartate transcarbamylase, dihydroorotase, (*CAD*); dihydroorotate dehydrogenase (*DHODH*); uridine monophosphate synthetase (*UMPS*)] to produce UMP, the precursor for all pyrimidine nucleotides, were identified in our screen as DMG dependencies (Figures 1C and D). We validated this DMG dependency on *de novo* pyrimidine synthesis using shRNA to knockdown expression of *CAD* and *DHODH.* As indicated, knockdown of either *CAD* or *DHODH* inhibited proliferation in all three DMG cells tested (Figures 1E, S1B and S1C). We further asked whether shRNA knockdown of *CAD* and *DHODH* was cytostatic or cytotoxic. As shown in Figure S1D, knockdown of either *CAD* or *DHODH* induced apoptosis of DMG cells as measured by Annexin V staining. Consistent with the above finding of DMG dependency on *de novo* pyrimidine biosynthesis, we observed upregulation of the pathway genes in two published RNA sequencing datasets of DMG patient tumor relative to the profiled matched control tissue (Berlow et al., 2018; Zhu et al., 2021). Significant upregulation of *CAD*, which catalyzes the committed step in *de novo* pyrimidine biosynthesis, was observed in both datasets (Figure S1E), highlighting the importance of the *de novo* pyrimidine biosynthesis pathway in DMGs.

**Figure 1.**
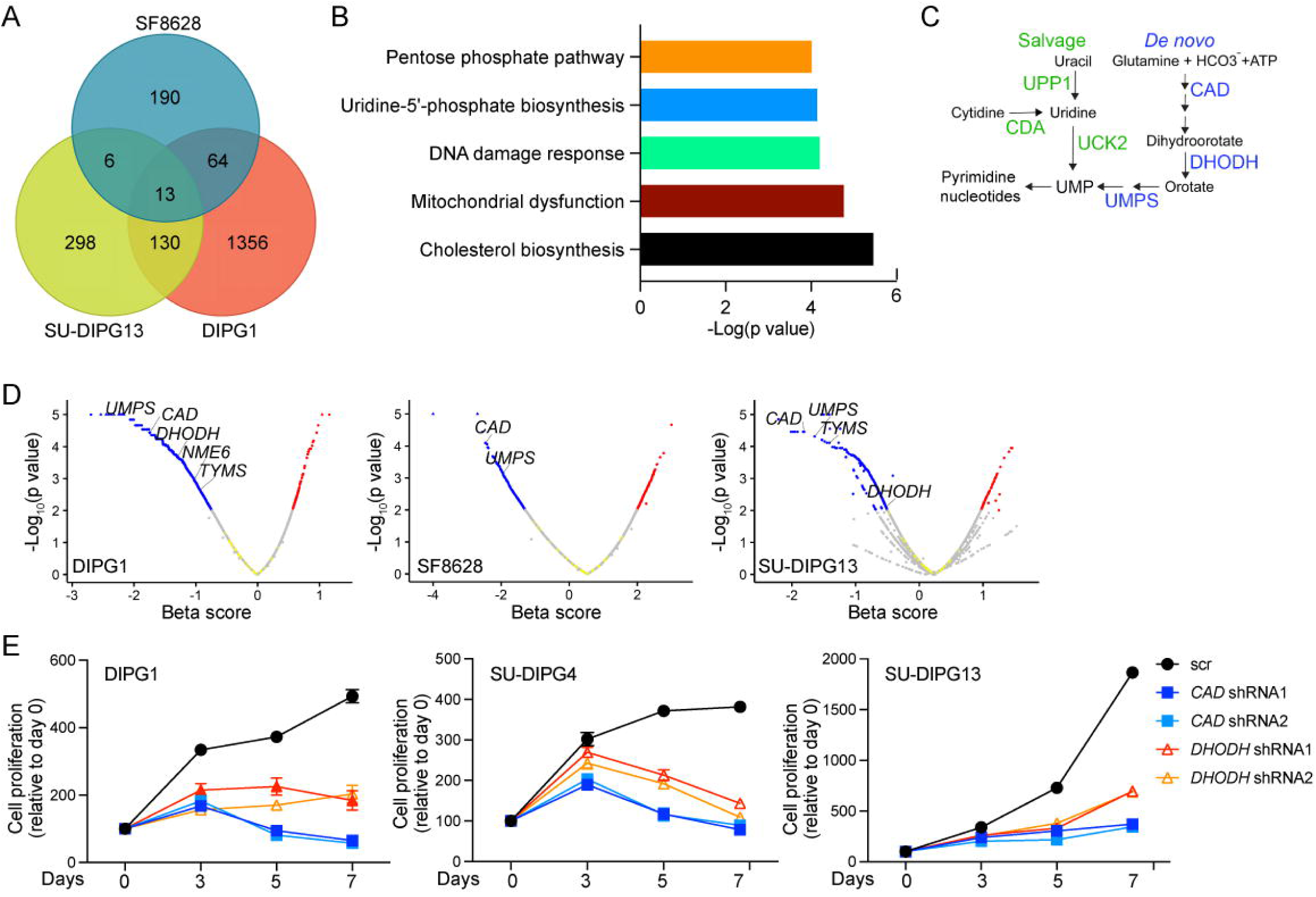
Genome-wide CRISPR screen identifies pathways critical for DMG cell survival. (A) Venn diagram shows overlap of genes identified as DMG dependencies in three tested DMG cell lines. (B) Ingenuity pathway analysis (IPA) defines DMG dependency pathways based on genes scored as dependency genes in at least two DMG cell lines. (C) Schematic of the *de novo* and salvage pathways for pyrimidine biosynthesis. (D) Volcano plots showing genes belonging to the Gene Ontology Resource (GO) pathway of pyrimidine biosynthesis (including uridine-5’-phosphate biosynthesis) whose depletion is significantly associated with DMG cell death (blue dots). (E) DMG cell proliferation following knockdown of *de novo* pyrimidine synthesis genes, *CAD* and *DHODH,* using two distinct shRNAs for each gene, relative to control (scr) shRNA. Data are represented as mean ± SEM (n=4). See also Figure S1 and Tables S1 and S2.

### *De novo* pyrimidine biosynthesis as a druggable DMG vulnerability

BAY2402234 is a clinical stage, small molecule inhibitor of DHODH. We evaluated the impact of BAY2402234 on DMG growth and viability using a panel of patient-derived cell lines derived from both H3 wild-type and H3 mutant DMG cell lines (Table S3). In addition, we asked whether dependency on *de novo* pyrimidine biosynthesis was DMG-specific or more broadly applicable to high-grade gliomas by testing the response of isocitrate dehydrogenase (IDH) wild-type adult glioblastomas (aGBM) and normal immortalized human astrocytes to BAY2402234. As indicated, all tested DMG cell lines were exquisitely sensitive to growth inhibition by BAY2402234 relative to aGBMs and astrocytes (IC50 range 0.11-0.63 nM for DMG versus 0.87-6.2 nM for aGBM, p=0.05; Figure 2A). Furthermore, we observed no correlation between proliferation rate and sensitivity to BAY2402234 indicating that DMG-specific sensitivity to DHODH inhibition was not proliferation-related (Figure S2A).

**Figure 2.**
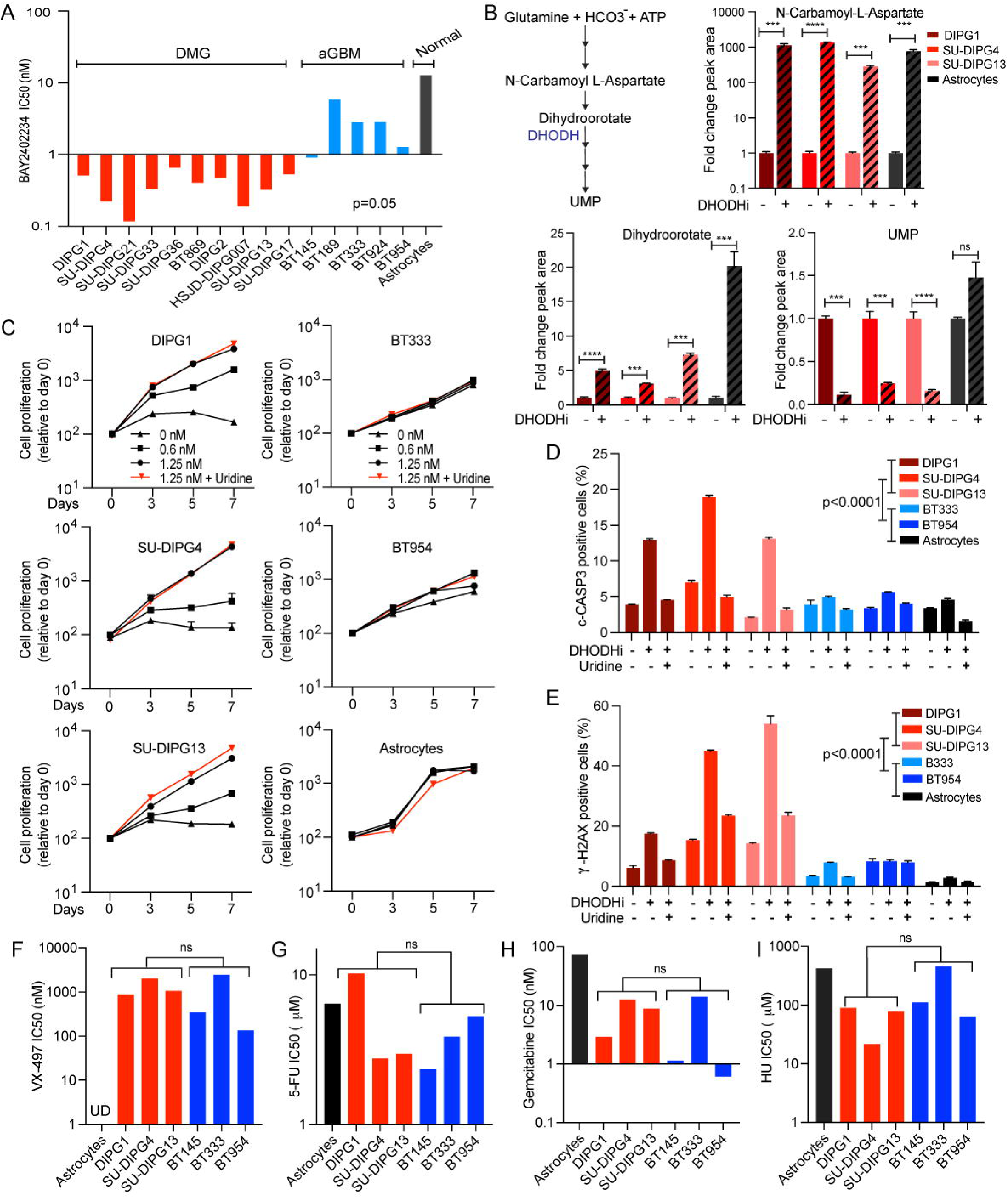
The DHODH inhibitor, BAY2402234, inhibits *de novo* pyrimidine synthesis and induces DNA damage and apoptosis specifically in DMGs compared to aGBMs and astrocytes; a differential sensitivity not observed towards *de novo* GMP synthesis inhibitor and chemotherapy agents. (A) DMG, adult GBM (aGBM), and immortalized human astrocytes (normal) were treated with increasing doses of BAY2402234 in quadruplicate for 5 days and IC50 values were determined using PRISM software. p=0.05, unpaired t-test of IC50 values for DMG vs aGBM. (B) Schematic representation of metabolites and DHODH in *de novo* pyrimidine biosynthesis (left). Liquid chromatography-tandem mass spectrometry (LC-MS/MS) analysis of control and BAY2402234 (DHODHi;1.25 nM, 24 hours) treated DMG cells. Bar graph show fold-change relative to control sample (mean ± SEM, n=3; * p<0.05, ** p<0.01, *** p<0.001, **** p<0.0001, unpaired t-test). (C) Proliferation of DMG, aGBM, and immortalized human astrocytes treated with increasing doses of BAY2402234 (DHODHi) as well as combination of 1.25 nM BAY2402234 plus exogenous uridine (100 µM). Data are plotted as mean ± SEM (n=4). (D and E) Apoptosis and DNA damage, as measured by flow cytometry for cleaved CASPASE-3 (c-CASP3)- and γ-H2AX-positive cells, respectively, in BAY2402234-treated (DHODHi; 1.25 nM, 48 hours) DMG, aGBM, and immortalized astrocytes cells with or without uridine supplementation (100 mM). Data show mean ± SEM (n=3), unpaired t-test. (F-I) DMG and aGBM cells were treated with increasing doses of (F) *de novo* GMP synthesis inhibitor (VX-497), (G) 5-fluorouracil (5-FU), (H) gemcitabine, and (I) hydroxyurea (HU) for 5 days in quadruplicate and IC50 values determined by PRISM software. ns represents not significant p value in unpaired t-test of IC50 for DMG versus aGBM groups. See also Figure S2 and Table S3.

To validate *de novo* pyrimidine synthesis as the BAY2402234 target, we quantified pathway metabolites in cells treated with BAY2402234 using liquid chromatography-tandem mass spectrometry (LC-MS/MS). Twenty-four hours after treatment with BAY2402234, metabolites that lie upstream of DHODH (N-carbamoyl-L-aspartate and dihydroorotate), accumulated in both DMGs and astrocytes (Figure 2B). In contrast, the pathway end-product, UMP, downstream of DHODH was depleted in BAY2402234-treated DMG cells, indicating on-target inhibition of *de novo* pyrimidine synthesis by the drug (Figure 2B). Of note, in astrocytes, UMP levels are not reduced by BAY2402234, consistent with lack of dependence of astrocytes on *de novo* pyrimidine synthesis for maintenance of UMP pools.

To confirm that growth inhibition of DMGs is an on-target response to suppression of *de novo* pyrimidine biosynthesis by BAY2402234, we asked whether supraphysiological uridine supplementation would rescue BAY2402234-induced growth inhibition. Validation by uridine rescue is possible because pyrimidine nucleotides can by synthesized by either the *de novo* or salvage pathway, but uridine supplementation specifically rescues inhibition of the former pathway by replenishing UMP pools. As indicated, exogenous uridine completely rescued the DMG specific dose-dependent effects of BAY2402234 on cell growth, (Figure 2C) and drug-induced apoptosis (Figures 2D and S2B. The essential roles of pyrimidines in DNA replication and repair led us to evaluate drug-induced DNA damage levels by quantifying γ-H2AX, a marker for DNA damage. Following BAY2402234 treatment, we documented 2-4-fold increases in γ-H2AX-positive DMG cells, and this induction of DNA damage was completely rescued by uridine supplementation (Figures 2E and S2C). Of note, BAY2402234 induced little to no apoptosis or DNA damage in aGBMs or astrocytes (Figures 2D and 2E).

We show above that DMGs are preferentially dependent on *de novo* pyrimidine biosynthesis. To ask whether this dependency was specific to pyrimidines, we used VX-497, a small molecule inhibitor of IMPDH, which specifically blocks *de novo* purine guanosine-5’-monophosphate (GMP) synthesis (Figure 2F). Unlike BAY2402234, VX-497 did not result in differential sensitivity of DMGs compared to aGBMs. Similarly, no pronounced sensitivity was observed in DMGs compared to aGBMs when treated with chemotherapy agents, 5-fluorouracil (5-FU), gemcitabine, and hydroxyurea, which all interfere with nucleotide synthesis and/or DNA replication (Figures 2G-I).

### Mechanism of DMG dependence on d*e novo* pyrimidine biosynthesis

We hypothesized that the marked sensitivity of DMGs to BAY2402234 (compared to aGBMs and normal astrocytes) is due to the extreme dependence of DMGs on *de novo* pyrimidine biosynthesis to maintain UMP pools. Consistent with this hypothesis, we observed more pronounced BAY2402234-induced depletion of UMP pools in DMGs (Figure 3A). Important here is the documentation that BAY2402234 successfully inhibited DHODH in all DMGs and aGBMs as N-carbamoyl-L-aspartate levels increased in all lines following drug treatment (Figure 3B).

**Figure 3.**
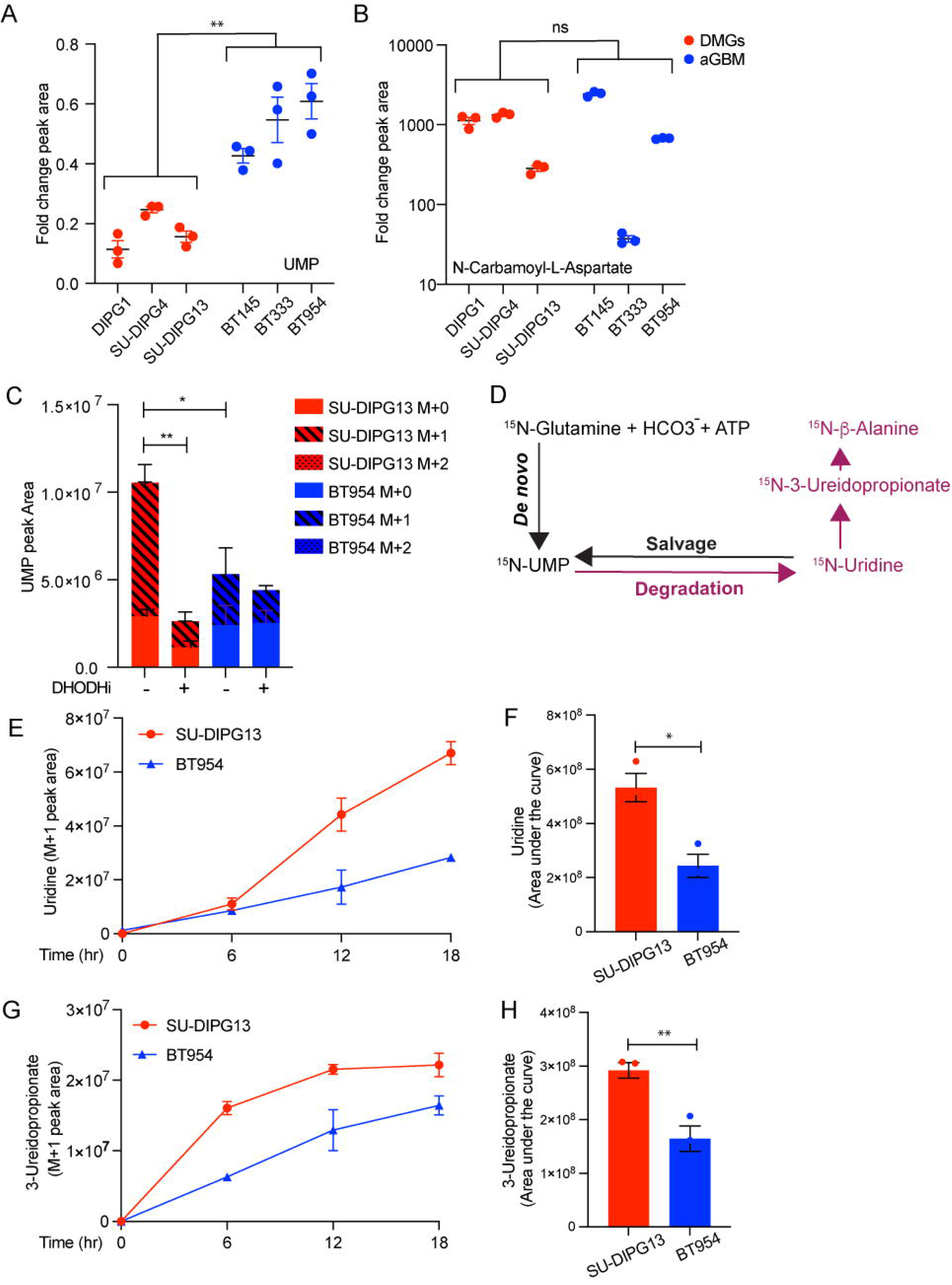
DMG exhibit increased flux through *de novo* pyrimidine biosynthesis and pyrimidine degradation, leading to profound depletion of UMP following *de novo* biosynthesis inhibition by BAY2402234. (A and B) Fold change in UMP and N-carbamoyl-L-aspartate in DMG and aGBM lines treated with BAY2402234 (DHODHi; 1.25 nM, 24 hours) (n=3). DMG and aGBM groups compared using unpaired t-test; ** p<0.01, ns-not significant. (C) Peak area of unlabeled (M+0) and labelled ^15^N-UMP (M+1) in SU-DIPG13 (DMG) and BT954 (aGBM) following 12 hours of exposure to media containing ^15^N-glutamine with or without BAY2402234 (DHODHi). For DHODHi samples, cells were pretreated with BAY2402234 for 16 hours prior to switching to media containing ^15^N-glutamine. * p<0.05 and ** p<0.01 for the comparisons of indicated M+1 peak areas, ANOVA with Tukey’s multiple comparison test. (D) Schematic of ^15^N-glutamine flux through *de novo* and salvage pyrimidine biosynthesis and pyrimidine degradation pathways. (E-H) Flux through pyrimidine degradation represented as peak areas of labeled ^15^N-uridine (M+1) and ^15^N-3-ureidopropionate (M+1) in SU-DIPG13 and BT954 exposed to media containing ^15^N-glutamine for specified times, as indicated. Panels F and H show area under the curve (unpaired t-test; *p<0.05 and **p<0.01). See also Figure S3.

To unequivocally demonstrate the marked reliance of DMGs on *de novo* pyrimidine biosynthesis we traced the incorporation of glutamine-nitrogen into UMP using amide-^15^N-labeled glutamine in DMG and aGBM. The nitrogen atom at position 3 of the uracil base is derived from the amide group of glutamine, while position 1 nitrogen is contributed by aspartate. Thus, in our tracing experiments, we expect only 1 nitrogen to be labeled through *de novo* synthesized UMP (M+1), while M+2 labeling, which represents UMP labeled at both positions, should be absent. As expected, we observe either unlabeled UMP (M+0) or M+1 UMP species (Figure 3C). Our proposed mechanism of unique DMG sensitivity makes a couple of predictions, both documented to be true in Figure 3C: (1) A larger fraction of the UMP pool in DMGs will be derived from glutamine through *de novo* biosynthesis, reflected in a larger M+1 peak (70% in DMG versus 50% in aGBM). (2) Depletion of labeled UMP by BAY2402234 will be more pronounced in DMG than aGBM (80% versus 35%, respectively). Internal controls validating the isotope tracing experiments are as follows: (1) labeled glutamine fractions are comparable in DMG versus aGBM (Figure S3A). (2) BAY2402234 effectively inhibits DHODH in both lines resulting in BAY2402234-induced increases in ^15^N-labeled dihydroorotate (Figure S3B).

Having confirmed, using stable isotope tracing experiments, *de novo* pyrimidine biosynthesis as the key source of UMP in DMG cells relative to aGMB, we sought to explore the underlying mechanism for this differential pathway dependency. Two possible mechanisms exist: (1) DMGs have reduced capacity to utilize the salvage pathway to compensate for inhibition of the *de novo* pathway; (2) DMGs display increased UMP and uridine degradation, resulting in less substrate (uridine) available for UMP synthesis through the salvage pathway. Mechanism 1 is negated by our data showing that exogenous uridine rescues all manifestations of BAY2402234 treatment (Figures 2C-E). In contrast, mechanism 2 is supported by studies showing greater conversion of ^15^N-labeled UMP (originating from ^15^N-labeled glutamine) into labeled uridine and 3-ureidopropionate in DMG relative to aGBM (Figures 3D-H, S3C). Thus, increased flux through the pyrimidine degradation pathway in DMGs results in limited availability of the substrate (uridine) required for synthesis of UMP through the pyrimidine salvage pathway, driving marked reliance on the *de novo* pathway for pyrimidine availability, and enhanced sensitivity to DHODH inhibition.

Having identified increased pyrimidine degradation as the mechanism underlying exquisite DMG sensitivity to DHODH inhibition, we asked whether this sensitivity was due to differential expression of genes responsible for pyrimidine homeostasis, including *de novo* or salvage pyrimidine synthesis, and pyrimidine degradation. Consistent with our stable isotope tracing results, we observed higher expression of dihydropyrimidine dehydrogenase (*DPYD)*, the first committed step of pyrimidine degradation, in DMGs relative to normal astrocytes (Figure S4A). Enhanced use of *de novo* pyrimidine biosynthesis in DMGs relative to aGBMs evident in our isotope tracing experiments cannot be explained by differences in *de novo* pathway gene expression (Figure S4A). In contrast, modestly higher levels of salvage pathway gene expression in aGBM compared with DMGs are consistent with greater reliance of DMGs on *de novo* pyrimidine biosynthesis (Figure S4A). We expanded these analyses to pre-treatment patient tumor samples by examining publicly available gene expression datasets (Cerami et al., 2012; Gao et al., 2013). As was true for the DMG cell lines, *DPYD* expression was higher in DMG primary tumors compared to pediatric non-DMG high-grade gliomas (Figure S4B). Of note, expression levels of *de novo* and salvage pathway genes were similar between DMGs and pediatric non-DMG high-grade tumors, again highlighting pyrimidine degradation as the key determinant of BAY2402234 sensitivity of DMGs (Figure S4B).

To further define a causal role for pyrimidine degradation in determining sensitivity to DHODH inhibition, we used genetic approaches to examine isogenic DMGs and aGBMs differing only in their DPYD expression. Using doxycycline-inducible shRNAs we diminished DPYD expression in DMGs and assessed sensitivity to BAY2402234. Reduced DPYD expression in DMGs curtailed tumor cell sensitivity to BAY2402234 in a dose-dependent fashion (Figures 4A-B, S4C). A pharmacologic approach confirmed our genetic approach as gimeracil, a DPYD inhibitor, also diminished DMG sensitivity to BAY2402234 (Figure S4D). Conversely, we over-expressed DPYD in aGBMs, and found enhanced sensitivity of aGBMs to BAY2402234 (Figures 4C-D).

**Figure 4.**
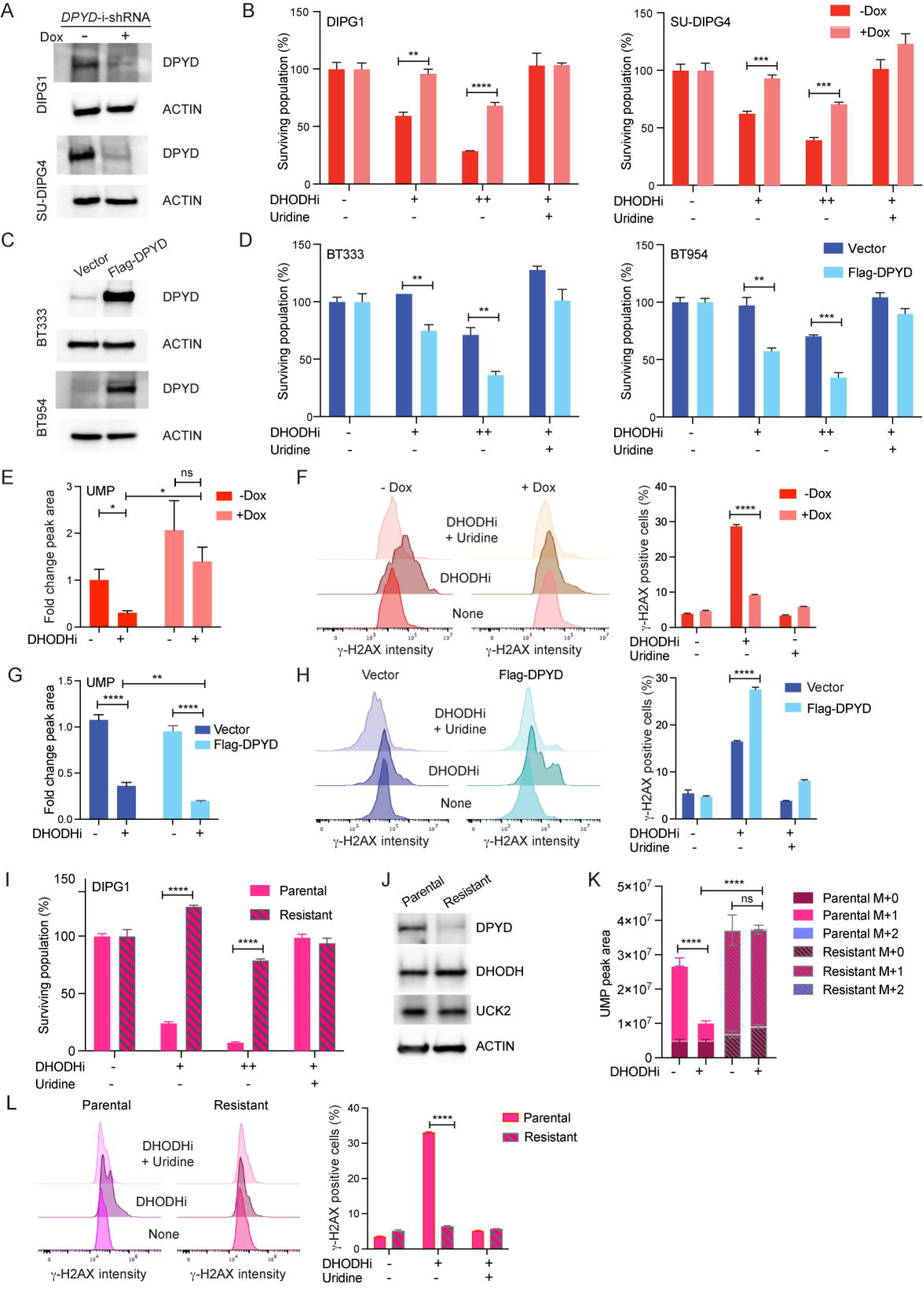
DPYD expression modulates sensitivity to BAY2402234. (A) Western blot analyses of extracts from DIPG1 and SU-DIPG4 cells expressing doxycycline-inducible shRNA against *DPYD* with or without doxycycline (Dox; 500 ng/ml) treatment for 72 hours. (B) DMG cells as in Panel 4A were treated with BAY2402234 (DHODHi; +1 nM, ++2 nM for DIPG1; +0.5 nM, ++1 nM for SU-DIPG4), with or without uridine supplementation (100 µM), for 72 hours and surviving populations quantified. Unpaired t-test, ** p<0.01, *** p<0.001, **** p<0.0001. (C) Western blot analyses of extracts from aGBM lines BT333 and BT954 expressing Flag-DPYD or empty vector as control. (D) aGBM cells as in Panel 4C were treated with BAY2402234 (DHODHi; +1.25 nM, ++2.5 nM for BT333; +0.5 nM, ++1 nM for BT954), with or without uridine supplementation (100 µM), for 6 days and surviving populations quantified. Unpaired t-test, ** p<0.01, *** p<0.001. (E and F) UMP peak area (E) and γ-H2AX as measured by flow cytometry (F) in DIPG1 cells as in Panel 4A, treated with BAY2402234 (DHODHi; 1 nM) for 24 and 48 hours, respectively. Unpaired t-test, * p<0.05, **** p<0.0001, ns-not significant. (G and H) UMP peak area (G) and γ-H2AX as measured by flow cytometry (H) in BT333 cells as in Panel 4C, treated with BAY2402234 (DHODHi; 2.5 nM) for 24 and 72 hours, respectively. Unpaired t-test, ** p<0.01, **** p<0.0001. (I-L) BAY2402234-resistant derivative of DIPG1 assessed for (I) sensitivity to BAY2402234 (DHODHi; +0.625 nM, ++1.25 nM) for 5 days. Unpaired t-test, * p<0.05, ** p<0.01, *** p<0.001, **** p<0.0001.; (J) Western blot analyses for indicated proteins; and (K) UMP peak area of unlabeled (M+0) and labeled ^15^N-UMP (M+1) in parental and resistant DIPG1 after exposure to media containing ^15^N-glutamine (for 12 hours) with or without BAY2402234. For DHODHi samples, cells were pretreated with BAY2402234 (DHODHi; 1.25 nM) for 14 hours prior to switching to media containing ^15^N-glutamine. **** p<0.0001, ns-not significant for the comparisons of indicated M+1 peak areas, ANOVA with Tukey’s multiple comparison test. (L) γ-H2AX (flow cytometry) in BAY2402234-treated (DHODHi: 1 nM, 48 hours) DIPG1 parental and resistant lines with or without uridine supplementation (100 µM). See also Figure S4.

We utilized our isogenic lines differing in DPYD expression to interrogate not only cell viability after BAY2402234 treatment but also BAY2402234-induced depletion of UMP pools and DNA damage. The marked BAY2402234-induced depletion of UMP pools seen in DMGs was reversed by decreased DPYD expression (using doxycycline-inducible shRNA) (Figure 4E). Similarly, induction of DNA damage by BAY2402234 (reflected in γ-H2AX staining) was markedly blunted by reduced DPYD expression (Figure 4F). Again, consistent with the causal role of DPYD in determining BAY2402234 sensitivity, over-expression of DPYD in aGBM increases BAY2402234-induced UMP pool depletion (Figure 4G) and DNA damage (Figure 4H).

To further corroborate the causal role of DPYD expression in DMG sensitivity to BAY2402234, we generated a drug-resistant DMG line derived from the parental DIPG1 cell line. The resistant line indeed shows dramatically reduced sensitivity to BAY2402234 treatment relative to the parental line (Figure 4I). Acquired resistance to BAY2402234 was associated with potent downregulation of DPYD expression, without changes in expression levels of DHODH and UCK2 (Figure 4J). These data reinforce our findings from DPYD silencing and overexpression studies (Figures 4A-H) and independently corroborate our conclusion that pyrimidine degradation is a key determinant of BAY2402234 sensitivity. BAY2402234 is unable to diminish UMP pools in the resistant line as evident in stable isotope tracing experiments of amide-^15^N-labeled glutamine (Figures 4K and S4E), despite efficient DHODH inhibition reflected in increased dihydroorotate levels after treatment in both parental and resistant lines (Figure S4F). Finally, in resistant DIPG1 cells, BAY2402234 also failed to induce DNA damage (Figure 4L). Taken together, our data indicate that DMG cells are reliant on *de novo* pyrimidine biosynthesis because of increased pyrimidine degradation leading to limited substrate for pyrimidine salvage.

### DHODH inhibition arrests cell cycle progression and leads to replication stress in DMG cells

Multiple cell functions that rely on pyrimidine nucleotides might be affected by BAY2402234 treatment that results in UMP depletion. Accordingly, we investigated BAY2402234’s effects on cell cycle progression and replication (Figure 5). BAY2402234 induced accumulation of DMG cells in S-phase, as indicated by BrdU uptake (Figure 5A). Given the documented accumulation in S-phase we asked whether BAY2402234 caused replication stress in DMG cells. To address this question, we measured DNA-bound Replication Protein A (RPA), a known indicator of replication stress. As a positive control we use treatment with hydroxyurea (HU), which depletes cellular dNTP pools and is known to induce replication stress. As shown (Figure 5B), we observe increased RPA foci after treatment with HU. We also documented increased RPA foci in BAY2402234-treated DMG cells, and this was completely rescued by exogenous uridine (Figure 5B).

**Figure 5.**
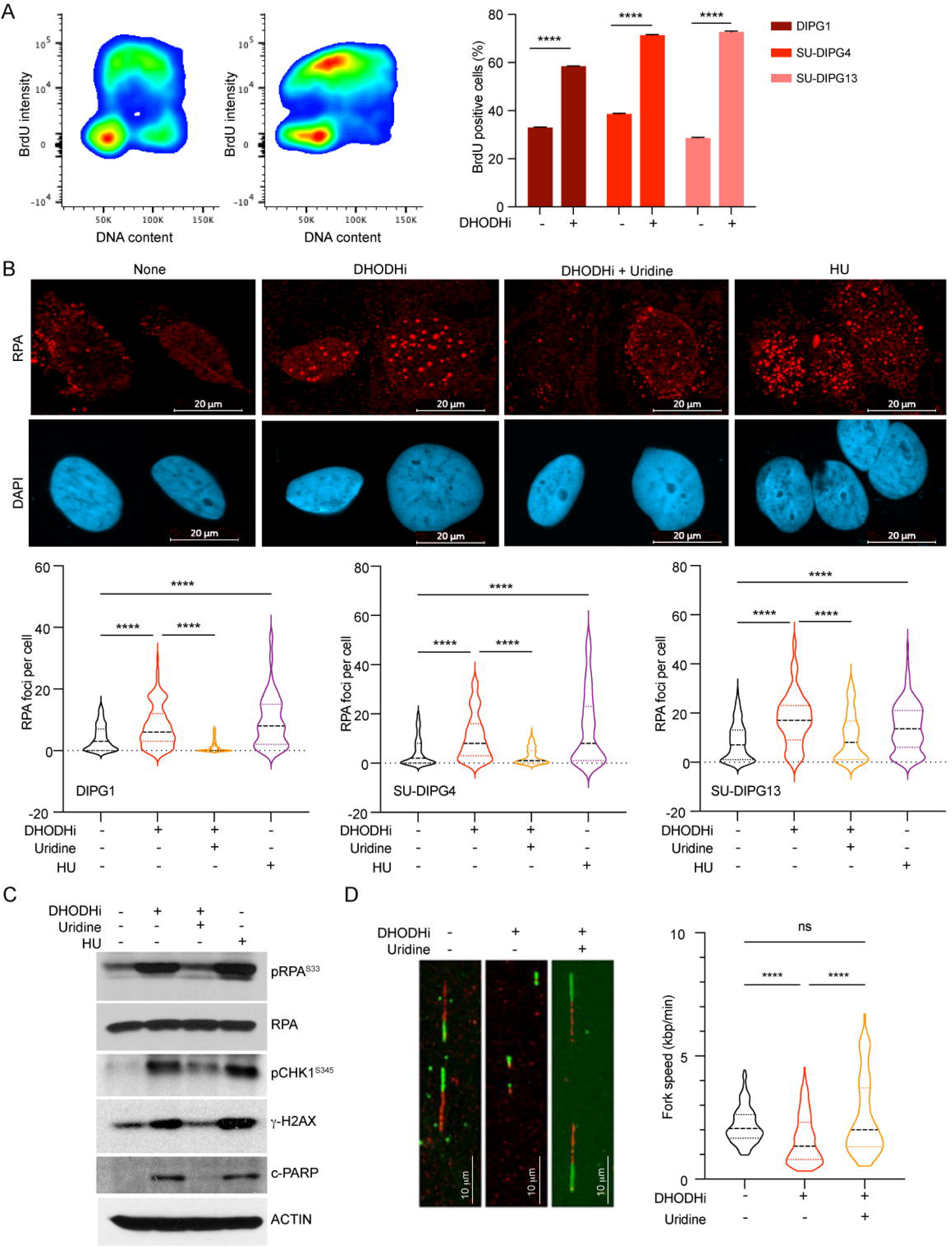
DHODH inhibition arrests DMG cells in S phase and increases replication stress. (A) S phase population, detected as BrdU positive cells by flow cytometry, in control and BAY2402234 (DHODHi; 1.25 nM, 24 hours) treated DMG cells. Distribution of SU-DIPG4 cells (left panels) and quantification of S phase populations (right panel; mean ± SEM, n=3). Unpaired t-test, **** p<0.0001. (B) BAY2402234-treated DMG cells (DHODHi; 1.25 nM, 24 hours) with or without uridine supplementation (100 µM), were analyzed for the presence of chromatin-bound RPAs by immunofluorescence staining of RPA2 foci. Hydroxyurea (HU), a known inducer of replication stress, is included as a positive control. Top panel shows representative images of RPA (red) and DAPI (blue). Violin plot shows the frequency distributions of foci/cell in at least 100 cells (bottom panel). Kruskal-Wallis test, **** p<0.0001. (C) Western blot analyses of whole cell extracts of SU-DIPG4 cells treated as in Panel 5B, evaluated for indicated proteins. pRPA2^S33^: phosphorylated RPA2 at serine 33. pCHK1^S345^: phosphorylated CHK1 at serine 345. c-PARP: cleaved PARP. (D) SU-SIPG4 cells treated with BAY2402234 (DHODHi; 1.25 nM, 48 hours; with or without 100 µM uridine) were sequentially labeled with CldU and IdU to document replicating DNA using DNA combing assay. Representative image is shown in the left panel. Fork speed was determined for at least 100 replicating fibers for each condition and represented as violin plots (the frequency distributions of the fork speed). Kruskal-Wallis test, **** p<0.0001, ns-not significant. See also Figures S5.

The presence of DNA-bound RPA activates ATR, which phosphorylates both RPA2 at serine 33 (S33), and the downstream effector CHK1 at serine 345 (S345). To confirm ATR activation after DHODH inhibition, we examined phosphorylation of RPA2 and CHK1 by Western blot analysis in DMG and aGBM (Figures 5C, S5). We observed induction of phosphorylation at RPA2^S33^ and CHK1^S345^ after BAY2402234 or HU treatment in DMG but only after HU treatment in BT333. Consistent with our previous findings, induction of replication stress and ATR activation were associated with increased levels of γ-H2AX and cleaved PARP, indicating increased DNA damage. Uridine replenishment rescued the BAY2402234-induced effects on RPA/CHK1 and DNA damage (Figure 5C), indicating the specificity of the drug and confirming that *de novo* pyrimidine biosynthesis mediates the phenotype. To further confirm actual stalling of replication forks we directly evaluated replication fork progression in BAY2402234-treated DMG using the DNA fiber combing assay. We observed that DHODH inhibition significantly decreased replication fork speed as indicated by smaller tracks of incorporated CldU and IdU (dUTP analog) after BAY2402234 treatment. Again, these findings were entirely reversed by uridine supplementation, as reflected in normal replication fork progression in DMGs treated with BAY2402234 in the presence of exogenous uridine (Figure 5D). Taken together, our data indicate that DHODH inhibition is efficacious against DMGs by impeding *de novo* pyrimidine biosynthesis, thus causing replication stress, and DNA damage.

### Inhibition of *de novo* pyrimidine biosynthesis by BAY2402234 prolongs survival of mice harboring DMG orthotopic tumors

Having demonstrated promising anti-tumor activity *in vitro*, we asked whether BAY2402234 can cross the blood brain barrier (BBB) and accumulate in DMG xenografts within murine brainstems. For this purpose, we implanted a luciferized DMG cell line (DIPG1) into the brainstem (pons) of mice to establish intracranial DMG xenografts and then administered BAY2402234 by oral gavage. After four days of treatment, brain tissues were collected for analysis using matrix-assisted laser desorption/ionization mass spectroscopy imaging (MALDI-MSI). As shown in Figures 6A and S6A-B, MALDI-MSI shows accumulation of BAY2402234 in intracranial pontine DMGs. To document the drug’s target engagement and *in vivo* biochemical activity, we quantified intra-tumoral levels of *de novo* pyrimidine biosynthesis metabolites upstream and downstream of DHODH. Following treatment with BAY2402234, N-carbamoyl-L-aspartate and dihydroorotate, which lie upstream of DHODH, accumulated, whereas UMP, which lies downstream of DHODH, diminished throughout the pontine tumors, compared to vehicle-treated mice (Figures 6B, S6C-D). As additional pharmacodynamic endpoints we measured DNA damage by γ-H2AX immunostaining and observed induction of DNA damage in BAY2402234-treated pontine DMGs (Figure 6C). All the *in vivo* pharmacodynamic results were consistent with our *in vitro* findings.

**Figure 6.**
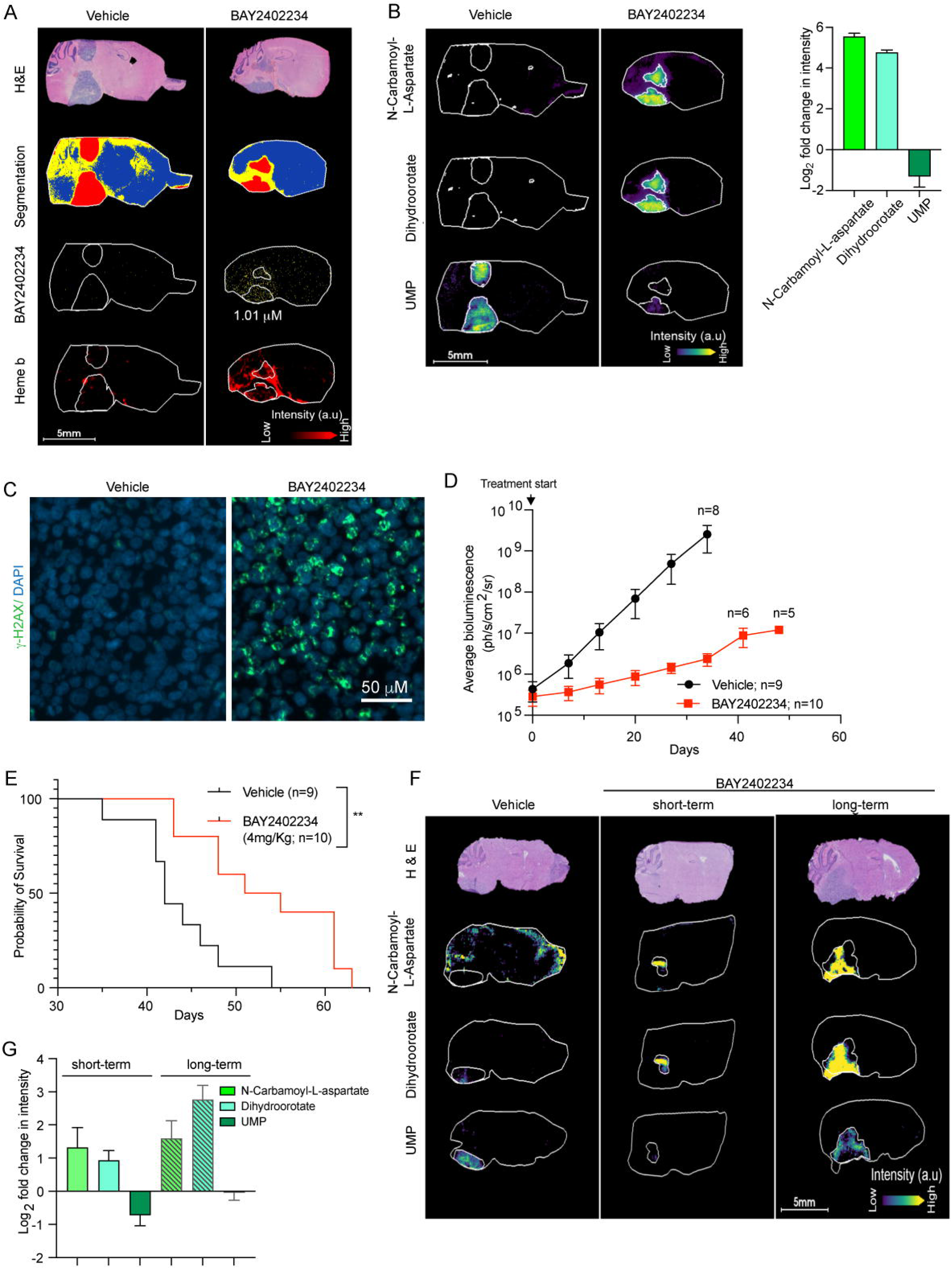
BAY2402234 is brain penetrant and inhibits *de novo* pyrimidine synthesis *in vivo* to prolong survival of mice bearing orthotopic DMG tumors. (A-C) MALDI MSI and optical microscopy imaging of brain tissue serial sections from mice harboring DIPG1 orthotopic tumor and treated with vehicle or BAY2402234 (4 mg/kg daily for 4 days). (A) Serial sagittal sections were stained with H&E and neighboring sections analyzed by MALDI MSI to quantitate levels of BAY2402234. Heme b was used as a marker of vasculature. (B) Metabolites of *de novo* pyrimidine synthesis analyzed by MALDI MSI (as in Panel 6A), shown in representative serial sections (left panel) and fold change quantified relative to vehicle treated section (right panel) from mice treated with BAY2402234 or vehicle (mean ± SEM, n=3 per treatment). (C) γ-H2AX staining of tumor regions in brain sections (same as Panel 6A) from vehicle- or BAY2402234-treated mice. (D) Tumor growth monitored by weekly bioluminescence imaging (BLI) of mice with SU-DIPG13P* orthotopic tumor; data represent average BLI signal values ± SEM. Number of mice is indicated by n. (E) Kaplan-Meier curve delineating survival of mice bearing SU-DIPG13-P* orthotopic DMG tumors treated daily (until euthanasia) with vehicle or BAY2402234. Log-rank (Mantel-Cox) test, ** p<0.01 (F and G) MALDI MSI and optical microscopy imaging of brain tissue serial sections from mice harboring SU-DIPG13-P* orthotopic tumors, treated with vehicle or BAY2402234 (4 mg/kg daily) using short-term (4 days) or long-term (35-37 days) drug regimens. (F) Representative images and (G) quantification of metabolites fold change relative to vehicle (mean ± SEM, n=3 per treatment). See also Figures S6.

We next asked whether BAY2402234 would prolong survival of mice bearing DMG intracranial xenografts. Towards this end, we used two DMG xenograft models; one was a highly aggressive cell line (DIPG1) with median survival of untreated mice of ∼7 days and the second was the SU-DIPG13-P* line which is moderately aggressive (median survival of untreated mice ∼43 days). BAY2402234 is well tolerated in mice and treatment significantly decreased tumor burden as measured by bioluminescence signals (Figures S6E, 6D, S6F). Moreover, BAY2402234 administration prolonged survival of mice in both highly and moderately aggressive DMG models (Figures 6E, S6F). For genetic validation of the BAY2402234 *in vivo* effects, we implanted luciferized DIPG1 cells transduced with shRNA targeting either *CAD* or *DHODH* or control shRNA (as in Figure 1E). While DIPG1 cells transduced with control shRNA formed pontine tumors *in vivo*, cells with *CAD* or *DHODH* knockdown failed to grow and establish tumors *in vivo* (Figure S6G).

As indicated in Figures 6E and S6F, we noted that although BAY2402234 markedly inhibited tumor growth and prolonged overall survival *in vivo*, DMG-bearing animals ultimately did succumb to their tumors. We asked whether this was due to an acquired inability of BAY2402234 to inhibit DHODH or the development of a compensatory mechanism to evade drug efficacy. To address this question, we performed MALDI MSI on tumors exposed to prolonged BAY2402234 treatment (35-37 days). After prolonged treatment, BAY2402234 continued to inhibit the pathway, as evident by accumulation of N-carbamoyl-L-aspartate and dihydroorotate; however, UMP levels were restored to untreated levels (Figures 6F and 6G). This finding mirrored results in DIPG1 cells with acquired resistance to BAY2402234 (Figure 4K) and in DMG cells in which DPYD was silenced (Figure 4E). Although *in vitro* we showed DPYD expression as the mechanism underlying modulation of sensitivity to BAY2402234, lack of robust anti-DPYD antibodies for immunohistochemistry precluded examination of DPYD expression *in vivo* in these resistant tumors. Nonetheless, these results nominate UMP pool preservation as a key feature of BAY2402234 resistance (Figures 6F and 6G), which is compromised by high pyrimidine degradation rates in *in vitro* models.

### Synergistic anti-cancer effects of DHODH- and ATR-inhibition in DMGs

We showed above that DHODH inhibition results in activation of ATR, a key kinase that is critical for mediating the DNA damage response induced by replication stress (Saldivar et al., 2017). We therefore hypothesized that simultaneously inhibiting ATR and DHODH would result in synergistic cytotoxicity in DMG cells. To test this hypothesis, we treated DMG cells with BAY2402234, elimusertib (ATR inhibitor), or a combination thereof. In DMG lines, combined DHODH and ATR inhibition dramatically augmented RPA foci formation, replication stress, and induction of DNA damage (Figures 7A-B, S7A-C) compared with either monotherapy. Figure 7B shows that elimusertib effectively inhibits ATR signaling, reflected in loss of RPA2^S33^ and CHK1^S345^ phosphorylation. Moreover, we observe enhanced DNA damage (γ-H2AX) and apoptosis (PARP and CASPASE-3 cleavage) resulting from combination therapy (Figure 7B) relative to either treatment alone. To confirm that elimusertib specifically inhibits ATR-mediated phosphorylation events, we examined an RPA site, T21, that is phosphorylated in response to replication stress in an ATR-independent fashion. We show that RPA^T21^ is phosphorylated after treatment with BAY2402234 or HU, a phosphorylation that persists even in the presence of elimusertib (Figure S7D).

**Figure 7.**
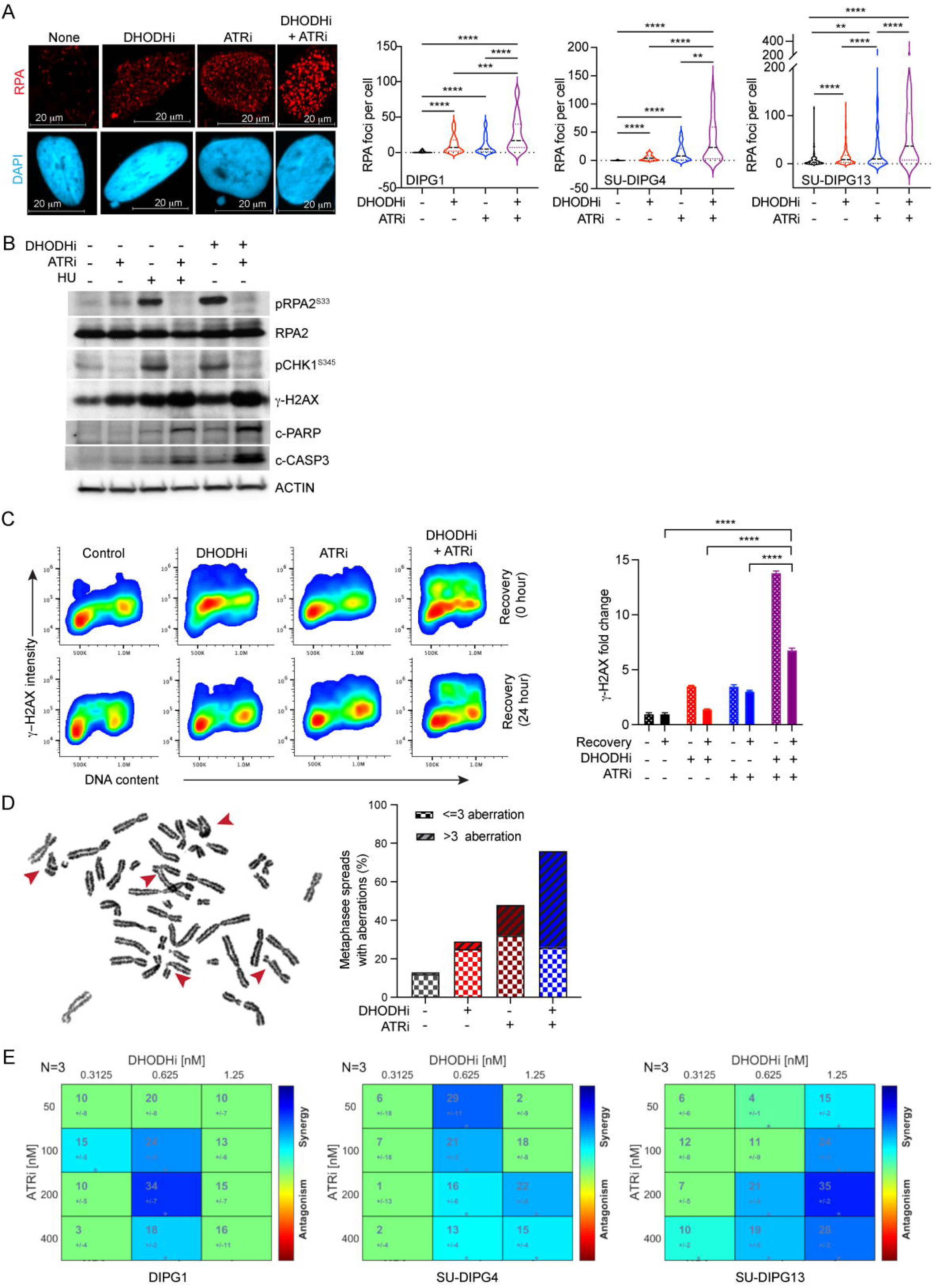
ATR inhibition augments DHODH inhibitor-induced replication stress and DNA damage. (A) Quantification of RPA foci by immunostaining in DMG cells treated (24 hours) with BAY2402234 (DHODHi; 1.25 nM), elimusertib (ATRi; 200 nM), or combination thereof. Representative images (left panel) and distributions of RPA foci/cell from at least 100 cells are represented as violin plots (right panel). Kruskal-Wallis test; ** p<0.01, *** p<0.001, **** p<0.0001. (B) Western blot analyses of indicated proteins, in SU-DIPG4 cells (treated as in Panel 7A). pRPA2^S33^: phosphorylated RPA2 at serine 33. pCHK1^S345^: phosphorylated CHK1 at serine 345. c-PARP: cleaved PARP. c-CASP3: cleaved CASPASE-3. (C) Flow cytometry quantitation of γ-H2AX-positive cells, in BAY2402234- (DHODHi; 1.25 nM; 24 hours), elimusertib- (ATRi; 200 nM; 24 hours), or combination-treated SU-DIPG4 cultures, before (top panel) or 24 hours after (bottom panel) drug washout. Representative flow cytometry density plots (left panel) and quantification of fold change (mean ± SEM, n=3; right panel). ANOVA, **** p<0.0001. (D) Metaphase spreads of SU-DIPG4 cells treated with BAY2402234 (DHODHi; 1.25 nM; 24 hours), elimusertib (ATRi; 100 nM; 24 hours), or combination therapy and scored for mitotic abnormalities (i.e., chromatid/chromosomal breaks, radials, etc.). Representative metaphase spread with red arrows indicating events scored as damage (left panel) and quantification of at least 25 metaphase spreads scored per treatment (right panel). (E) Synergistic cytotoxicity induced by combination therapy of DHODH- and ATR-inhibitors. DMG cells were treated with specified doses of BAY2402234 and elimusertib (72 hours; triplicates). Cell viability was determined by Cell-Titer Glo and analyzed using Combenefit software to assess synergy distributions and synergy-antagonism relationships. n=3 and * p<0.05. See also Figures S7.

Next, we asked whether replication stress induced by DHODH inhibition is reversible. DMG cultures were treated with BAY2402234 for 24 hours. The drug was then removed, and cells were cultured for an additional 24 hours. We monitored DNA damage by γ-H2AX expression immediately before and after the 24-hour drug recovery period (Figure 7C). We observed that the limited DNA damage induced by BAY2402234 or elimusertib monotherapy was partially reversible. In contrast, the considerable levels of DNA damage induced by 24 hours of combination treatment proved irreversible in approximately 50% of cells, suggesting replication fork collapse.

We sought to quantitate the degree of chromosome/chromatid aberrations that resulted from either monotherapy or combination treatment. Cells were allowed to progress into mitosis and then captured at metaphase using colcemid. Analysis of metaphase spreads disclosed chromosome/chromatid aberrations in 29% after DHODH inhibition, 48% after ATR inhibition, and 76% after combined DHODH and ATR inhibition (Figure 7D).

The marked induction of replication stress, DNA damage, and chromosome/chromatid aberrations suggested that combined inhibition of DHODH and ATR would also lead to synergistic cell death. We tested this prediction using the Combenefit software (Di Veroli et al., 2016) to analyze monotherapy and combination treatments of DMG cultures. We found synergistic DMG cell killing by combined BAY2402234 and elimusertib therapy (Figure 7E). The marked synergy seen with BAY2402234 and elimusertib combination treatment confirms replication stress as a mechanism of cell death induced by DHODH inhibition.

## DISCUSSION

DMGs are thought to arise from developmentally stalled progenitor cells that are locked into a replication competent state (Filbin et al., 2018), driven by amino acid substitution mutants of histone H3. These mutant histone oncoproteins function to reconfigure chromatin and are essentially undruggable (Larson et al., 2019; Nagaraja et al., 2019; Silveira et al., 2019). Against this backdrop, we undertook an unbiased genome-wide CRISPR screen to identify DMG vulnerabilities. The dependencies identified in our screen included previously documented reliance on cholesterol biosynthesis and TCA cycle dysregulation (Chung et al., 2020; Phillips et al., 2019). However, one robust dependency identified and validated herein is the *de novo* pyrimidine biosynthesis pathway. A companion study (Shi et al.) describes similar findings for IDH-mutant glioma--another glioma common in young people.

Purines and pyrimidines are the *sine qua non* of replicative DNA synthesis and the metabolic pathways involved in nucleotide biosynthesis have historically been attractive targets for cancer drug development, like methotrexate (Robinson et al., 2020). Accordingly, it is not surprising that multiple cancer types are vulnerable to inhibitors of *de novo* pyrimidine biosynthesis (Brown et al., 2017; Mathur et al., 2017; White et al., 2011). The list includes adult glioblastoma stem-like cells that are more sensitive to inhibition of *de novo* pyrimidine synthesis than their non-stem counterparts. However, unlike DMGs studied here that are selectively dependent on *de novo* pyrimidine biosynthesis, adult glioblastoma stem cells show a general reliance on nucleotide metabolism for cell growth (Wang et al., 2019b; Wang et al., 2017).

DMGs are selectively and exquisitely sensitive to inhibition of *de novo* pyrimidine synthesis. We interrogated the mechanism underlying this marked sensitivity and found that DMGs upregulate pyrimidine degradation, which consumes pyrimidine salvage pathway substrates (uridine and uracil) and augments UMP depletion when the *de novo* pathway is blocked. We further show that expression of *DPYD* is higher in DMGs and that DPYD expression levels dictate BAY2402234 sensitivity in isogenic glioma systems. These findings identify a possible mechanism of sensitivity to DHODH inhibition in DMGs and also suggest DPYD expression as a predictive biomarker of response to DHODH inhibitor therapy in this disease and perhaps other tumor contexts. A fundamental question is how DPYD, and enhanced pyrimidine degradation might contribute to the malignant phenotype. This question is pertinent beyond DMGs as DPYD and its enzymatic product, dihydrouracil, have been shown to drive tumor aggressiveness in various cancers (Shaul et al., 2014; Siddiqui and Ceppi, 2020).

Drug resistance is an important concern and consideration in cancer medicine as emergence of resistance to monotherapy is the rule rather than the exception. Accordingly, we also interrogated mechanisms of acquired resistance to DHODH inhibition in DMGs. Our findings indicate that resistance emerges in DMGs after prolonged BAY2402234 exposure, both *in vitro* and *in vivo*. Furthermore, DPYD downregulation is observed in DMG cells with acquired resistance to BAY2402234, highlighting the promise of DPYD as a predictive biomarker of drug resistance. Here, the synergistic response of DMGs to the combination of BAY2402234 and the ATR antagonist elimusertib suggests a strategy for combating resistance to BAY2402234 monotherapy. The serine/threonine protein kinase ATR regulates the intra-S-phase checkpoint and maintains genome stability in the presence of stalled replication forks that are induced by DHODH antagonists such as BAY2402234. Since ATR plays no role in pyrimidine biosynthesis, resistance to combination DHODH and ATR inhibitor therapy in DMGs would require the unlikely emergence of concurrent mutations in two independent biological pathways.

It is noteworthy that many oncogenic drivers and tumor suppressors that were first identified in childhood cancers, have gone on to prove widely relevant to adult cancers, including ETV6/RUNX1 (most common fusion in childhood acute lymphoblastic leukemia), EWSR1/FLI1 (gene fusion that defines Ewing sarcoma), and Wilms’ tumor gene (WT1) (Jiang et al., 2021; Rahal et al., 2018; Rosenfeld et al., 2003). Of relevance to this study the histone H3 amino acid substitutions first observed in pediatric DMGs have now been reported in adult solid tumors such as acute myeloid leukemia and melanoma (Lowe et al., 2019). Likewise, mutations in IDH1 or IDH2, first reported in gliomas of young adults, have since been noted as common driver mutations of acute myeloid leukemias that occur most commonly in older adults. For the road ahead, it will be of interest to see if DHODH antagonists are efficacious in genetically informed clinical trials of adult cancers, moving away from histology and toward genetic underpinnings to guide therapy.

## ACKNOWLEDGEMENTS

We thank Dr. Charles D. Stiles, Dr. William G. Kaelin, Jr. and Dr. Antje M. Wengner for helpful suggestions and discussions, Dr. Myles Brown for providing the human CRISPR knockout libraries H1 and H3, Dr. Naiara Santana-Codina and Dr. Shrabasti Roychowdhury for guidance in cell-based metabolite analysis and DNA fiber assays, respectively. This research was supported in part by the William M. Wood Foundation (D.H-K), Seed Grant in Epigenetics and Gene Dynamics, Harvard Medical School (D.H-K), Defeat DIPG ChadTough Research Grant (D.H-K), and Innovations Research Fund of Dana-Farber Cancer Institute (D.H-K). M.R.S is supported by NCI (F30CA271634), N.Y.R.A is supported by NIH grants U54-CA210180 (N.Y.R.A), P41-EB028741 (N.Y.R.A), T32EB025823 (S.A.S) and by the Pediatric Low-Grade Astrocytoma Program at PBTF (N.Y.R.A). S.K.M is supported by awards from the NCI (R01CA258586), the Cancer Prevention and Research Institute of Texas (RR190034), and the V Foundation for Cancer Research (V2020-006).

## AUTHOR CONTRIBUTIONS

Conceptualization, S.P, D.C, S.K.M, N.Y.R.A and D.H-K ; Methodology, S.P, S.A.S, M.R.S, Q-D.N, K.L.J, R.D, X.Z, N.Y.R.A, S.K.M and D.H-K; Investigation, S.P, J.P.K, S.A.S, M.R.S, Q-D.N, K.L.J, L.A.M, A.J.P, M.G.D, B.R.H, R.D, X.Z and S.K.M; Validation, S.P, J.P.K, S.A.A , M.R.S, H.N, N.Y.R.A , S.K.M and D.H-K; Formal analysis, S.P, J.P.K, H.N, S.A.S, M.R.S, K.L.J, L.A.M, R.D, X.Z, N.Y.R.A and S.K.M ; Visualization, S.P, H.N, S.A.S, M.R.S, N.Y.R.A, S.K.M and D.H-K; Writing – Original Draft, S.P, J.P.K, K.X.L, D.H-K; Writing – Review & Editing, All authors; Funding Acquisition, D.H-K, S.K.M, and N.Y.R.A; Resources, J.P, N.Y.R.A, M.G.F, S.K.M and D.H-K ; Project administration, D.H-K; Supervision, D.H-K., D.C, S.K.M, and N.Y.R.A.

## DECLARATION OF INTERESTS

N.Y.R.A is key opinion leader for Bruker Daltonics, scientific advisor to Invicro, and receives support from Thermo Finnegan and EMD Serono. S.K.M. has served as a paid advisor to Agios Pharmaceuticals. The other authors declare no competing interests.

## STAR Methods

### RESOURCE AVAILABILITY

#### Lead Contact

Further information and requests for resources and reagents should be directed to and will be fulfilled by the lead Contact, Daphne Haas-Kogan

#### Materials Availability

This study did not generate new unique reagents.

#### Data and Code Availability

- CRISPR screen data reported in this article is deposited on Mendelel at (Mendeley Data, V1, doi: 10.17632/yy2w57vkns.1) and will be publicly available as of the date of publication.
- This paper reports data derived from existing published and analyzed datasets. The dataset sources are listed in the key resource table. All other data reported in this paper will be shared by the lead contact upon request.
- This paper does not report original code.
- Any additional information required to reanalyze the data reported in this paper is available from the lead contact upon request.

## EXPERIMENTAL MODEL AND SUBJECT DETAILS

### Cell lines and culture conditions

SF8628 (female) was obtained from the Brain Tumor Research Center (BTRC) Tissue Bank at the University of California, San Francisco (UCSF, San Francisco, CA) and authenticated by the UCSF Genomics Core using short tandem repeat (STR) profiling. Cell lines BT145 (male), BT189 (male), BT333 (female), BT424 (male), BT954 (male), BT924 (male), BT189 (male), BT869 (female), SU-DIPG13 (female), and SU-DIPG17 were obtained from The Center for Patient Derived Models (CPDM) at Dana-Farber Cancer Institute (DFCI), and authenticated by STR profiling, except SU-DIPG17, at CPDM; CCHMC-DIPG1 (male) and CCHMC-DIPG2 (male), referred as DIPG1 and DIPG2 were obtained from Dr. Rachid Drissi at Cincinnati Children’s Hospital Medical Center. The SU-DIPG13P* (female), SU-DIPG4 (female), SU-DIPG21, SU-DIPG33, and SU-DIPG36 lines were provided by Dr. Michelle Monje at Stanford University, and HSJD-DIPG007 was obtained from Dr. Angel Montero Carcaboso at Hospital Sant Joan de Deu, Barcelona, Spain. Immortalized astrocytes were purchased from ATCC (SVG p12). SF8628, and immortalized astrocytes were cultured as adherent cells in DMEM supplemented with 10% and 5% FBS, respectively and penicillin-streptomycin (1X). All other cell lines were cultured in 1:1 neurobasal media and DMEM F-12 with MEM sodium pyruvate solution (1 mM), MEM non-essential amino acids (1X), GlutaMAX-I supplement (1X), HEPES buffer (10 mM), penicillin-streptomycin (1X), B27 supplement (1X), human-EGF (20 ng/ml), human-FGF2 (20 ng/ml), PDGF-AA (10 ng/ml), PDGF-BB (10 ng/ml), and 0.0002% heparin. All lines except SF8628, immortalized astrocytes, SU-DIPG4, SU-DIPG21, SU-DIPG33, and SU-DIPG36 were maintained as neurospheres, with media renewed every 4 days and dissociated every week by repeated pipetting or accutase when single cell population was required. All cells were cultured aseptically at 37°C in humidified incubators with 5% CO_2_. Cell line authentication, if performed, is noted above. Gender of the cell lines, if known, is noted above.

### Animals and housing conditions

Animal experiments were performed after approval by the Dana-Farber Institutional Care and Use Committee (IACUC) and were conducted as per NIH guidelines for animal welfare. Animals were housed and cared according to standard guidelines with free access to water and food. All experiments were performed on 5 weeks-old female NSG mice (NOD.Cg-Prkdcscid Il2rgtm1Wjl/SzJ). Mice were randomly distributed among treatment groups following orthotopic DMG cell implantation and a maximum of 5 mice were maintained per cage per treatment group. Mice were euthanized as they developed the neurological symptoms.

## METHOD DETAILS

### Drugs and chemicals

BAY2402234 (DHODH inhibitor), and elimusertib (ATR inhibitor), were obtained from Bayer AG under a material transfer agreement. Hydroxyurea (HU) was purchased from Sigma-Aldrich, gimeracil (DPYD inhibitor) from Santa Cruz Biotechnology, and VX-497, 5-fluorouracil (5-FU), and gemcitabine from Selleck Chemicals.

### Genome wide CRISPR screen for gene deletion

SF8628, DIPG1 and SU-DIPG13 (2×10^8^ cells) were transduced with the human CRISPR sgRNA library by spinfection. For SF8628 and DIPG1 cells the H1 library (Addgene #1000000132) (Xu et al., 2015) was used while the screen in SU-DIPG13 was performed using the H3 library (Addgene #133914). Cells were resuspended at 1×10^6^/ml in 50 ml tubes, mixed with pre-determined lentiviral supernatant volume that corresponds to a MOI (multiplicity of infection) of ∼0.3 and 8 μg/ml polybrene (Santa Cruz Biotechnology) and centrifuged at 200 RPM for 45 minutes at room temperature. Cells were then resuspended and plated in 75 cm^2^ flasks for 48 hours. After 48 hours, cells were collected and re-plated in fresh media with puromycin (SF8628 and SU-DIPG13: 2 µg/ml; DIPG1: 80 µg/ml). After 72 hours, half of the cells were collected by centrifugation, washed, and stored at -80°C as the P0 sample. The remaining cells were collected, re-plated in fresh media and cultured for another 10 passages before they were harvested as P10 samples. At least 2-3×10^7^ cells were plated at each passage. Two biological replicates of the screen were performed for each cell line.

To isolate genomic DNA, cell pellets were resuspended in DMS cell lysis buffer (10 mM Tris HCl pH 8.0, 1 mM EDTA, 0.2% SDS, 300 mM NaCl) at 1×10^7^ cells/ml with RNAase A at 100 mg/ml and incubated at 65°C for 1 hour. Proteinase K was added at a final concentration of 100 mg/ml and incubated at 55°C with rotation overnight. Next day, genomic DNA was purified by phenol:chloroform:isoamyl alcohol (25:24:1) extraction followed by isopropanol precipitation and 70% ethanol wash. The DNA was resuspended in water and re-extracted as above and finally dissolved in water and stored at 4°C overnight before proceeding with library preparation. For library preparation, 200 mg of DNA per condition were PCR amplified using Q5 high fidelity DNA polymerase (NEB Inc.) for 16 cycles using primers (Forward: 5’AATGGACTATCATATGCTTACCGTAACTTGAAAGTATTTCG3’; Reverse: 5’TCTACTATTCTTTCCCCTGCACTGTACCTGTGGGCGATGTGCGCTCTG3’). A second round of PCR was performed on the above amplified products for 8 cycles to attach Illumina sequencing adapters and to barcode samples. The PCR products were run on 2% agarose gel and purified using the gel purification kit before they were sent out for sequencing on the Illumina HiSeq (PE150) platform (Novagene Inc.).

MAGeCK was used to process and analyze the CRISPR screen data as follows (Li et al., 2014). First, for each sample, the count function of MAGeCK was used to map the paired-end sequencing data against the library to generate the sgRNA-level counts data for analysis. The mapping was performed separately for each of the two read sets against each of the two orientations of the library. The sgRNA-level counts dataset from the single read-library orientation combination with the highest mapping percentage was selected for further analysis. Quality control checks were performed based on the statistics generated from the MAGeCK’s count function (Wang et al., 2019a). Most replicate samples had mapping success rates of at least 65%. There were two samples with mapping percentages lower than 65%, but above 60%. All day-0 samples had low Gini indices (<0.15), that are measures of evenness of sgRNA counts, and low percentages of zero-count sgRNAs (<1%). Next, the MAGeCK MLE was used to analyze the sgRNA-level counts data to determine whether the abundance of sgRNAs for each gene were different between P10 and P0 samples for each cell line. The analysis was performed on data of the two replicates together in paired mode and using the negative control sgRNAs that targeted the AAVS1 loci for normalization (Chen et al., 2018). We then determined the significant negatively selected genes from the MAGeCK MLE results of each cell line. A gene was considered as a significant negatively selected gene if it had a negative beta score and *p*-value of less than 0.01. To confirm the biological significance of these genes, we checked the representation of pan-cancer essential genes (Table S4), and they represented a large fraction (>45%) of the negatively selected genes. Finally we filtered out pan-cancer essential genes (Table S4) from the significant negatively selected genes to define the “DMG specific dependency genes” and performed pathway analysis on the gene set that was identified as DMG specific dependency in at least two of the DMG lines using the Core Analysis of IPA (QIAGEN Inc., https://www.qiagenbioinformatics.com/products/ingenuitypathway-analysis; release 2019-06-15).

### Lentiviral production and DMG cell infection

For lentiviral production, shRNA clones targeting *CAD* (TRCN0000045910, TRCN0000045908) and *DHODH* (TRCN0000025868, TRCN0000025839) were purchased from Sigma-Aldrich, doxycycline inducible shRNA targeting *DPYD* (V2THS_84048, V2THS_84045) were purchased from Horizon Discovery while the lentiviral vectors expressing scrambled shRNA (Sarbassov et al., 2005) and flag-tagged *DPYD* (Shaul et al., 2014) were bought from Addgene Inc. Briefly, 90% confluent HEK29T3 cells in 100 mm plates were transfected with pMD2.G (1.5 μg; Addgene plasmid # 12259), psPAX2 (3.0 μg; Addgene plasmid # 12260), and shRNA plasmid (6.6 μg) using Lipofectamine 2000 as per manufacturer’s instructions. Viral supernatant was harvested at 48- and 72-hours post-transfection, combined, centrifuged at 3,000 rpm for 10 minutes to remove debris, and filtered through 0.45 μm filter. Viral supernatant was either immediately used or stored at −80°C. Five ml of the lentivirus containing supernatant was used to transduce 1×10^6^ DMG cells in the presence of 8 μg/mL polybrene and the cells were spun for 30 minutes at 200 rpm before they were plated. After 24 hours, cells were replated in fresh media to remove lentiviral particles and used to set up downstream experiments or establish stable cell lines.

### Proliferation assays

To follow cell proliferation after shRNA transduction, cells were harvested after 24 hours of infection, 1000 cells were plated per well in fresh media in 96 well plates in quadruplicates (day 0), and the remaining cells were cultured for additional 48 hours before total RNA was extracted to measure gene knockdown using RT-qPCR. Cell proliferation was followed using CellTiter-Glo as per manufacturer’s instruction in quadruplicates for each condition on Day 0, 3, 5 and 7 post-plating in 96 well plates and represented relative to day 0. To follow cell proliferation after BAY2402234 treatment, 500 cells per well (except for SU-DIPG4 and immortalized human astrocytes; 250 cells per well) were plated in quadruplicate in 96 well plates overnight before BAY2402234 was added and CellTiter-Glo signal was measured as above. To assess specificity of BAY2402234, cells were plated and maintained in either low or high uridine media which included 10 nM or 100 µM supplementation of uridine, respectively.

### Reverse transcription-quantitative polymerase chain reaction (RT-qPCR) assay

Total RNA was isolated for RT-PCR analysis using Trizol (Invitrogen). Reverse transcription was performed on 2 μg of total RNA using Superscript IV (Invitrogen Inc.) and real-time PCR was performed on Roche Light Cycler 96 using the specific primers with SYBR Green chemistry (see Table S5 and key resource table). Expression of mRNA was calculated using the ΔΔ*C*_t_ method relative to scrambled shRNA-treated sample or to astrocytes and normalized to 18S rRNA.

### Cell treatment, and drug sensitivity

For all cell-based assays, cells were plated in media supplemented with 10 nM uridine and for uridine rescue experiments, media was supplemented with 100 µM uridine. Unless specified otherwise in figure legends, BAY2402234, elimusertib, and HU were used at a final concentration of 1.25 nM, 200 nM and 500 nM, respectively. To measure DMG drug sensitivities, 500 cells were plated in quadruplicates or triplicates in 96-well plates for all lines (except for SU-DIPG4 and immortalized human astrocytes; 250 cells per well) and treated as indicated in figure legends. Cell viability was assessed by CellTiter-Glo assay (timepoints noted in figure legends) to determine percentage of surviving cells and calculate IC50. For analysis of synergy and antagonism relationship, Combenefit software was used (Di Veroli et al., 2016).

### Steady-state metabolite analysis

2×10^6^ cells were plated overnight before addition of BAY2402234 (1.25 nM, unless noted otherwise in figure legends) for 24 hours. Next day, cells were supplemented with 2 ml of fresh media with or without drug and after 2 hours cells were harvested and processed in the cold room. Cells were collected by centrifugation and washed with cold saline solution. Pellets were resuspended in 300 µL of cold 80% methanol (LC-MS grade Honeywell catalog number 34966 diluted with LC-MS grade water from Pierce catalog number 51140) and incubated for 20 minutes at 4°C in a shaker. Subsequently, samples were centrifuged to remove debris, 200 µL of the extract was transferred to a new tube, dried in a SpeedVac and stored at -80°C until LC-MS analysis. For metabolite analysis samples were re-suspended in 20 µL HPLC grade water and 5-7 µl were injected and analyzed using a hybrid 6500 QTRAP triple quadrupole mass spectrometer (AB/SCIEX) coupled to a Prominence UFLC HPLC system (Shimadzu) via selected reaction monitoring (SRM) of a total of 289 endogenous water-soluble metabolites for steady-state analyses of samples as previously described (Yuan et al., 2012). Experiments were performed with three biological replicates and data were normalized using MinMax normalization across all measured metabolites in each sample before abundance of a metabolite was calculated relative to the DMSO treated sample.

### ^15^N-Amide glutamine isotope tracing through pyrimidine metabolism pathway

To determine the contribution of *de novo* pathway in UMP biosynthesis, ^15^N-glutamine tracing experiments were performed in DMG (SU-DIPG13), aGBM (BT954) and DIPG1 parental and resistant derivative. 2×10^6^ (SU-DIPG13 and BT954) or 1×10^6^ (DIPG1 parental and resistant derivatives) were plated overnight before addition of BAY2402234. Cells were treated with DMSO or BAY2402234 (SU-DIPG13, BT954: 0.5 nM, 16 hours and DIPG1 parental and resistant derivative: 1.25 nM, 14 hours). Next, samples were washed (2 times with glutamine free media) and re-plated for 12 hours in glutamine free media that was supplemented with either 2 mM ^15^N-glutamine (Cambridge isotope laboratories) or 2 mM GlutaMAX (Gibco Inc.; represents 0 hours of ^15^N-glutamine tracing) with or without BAY2402234 as above. Finally, cells were collected and processed to dried pellets as described above in “Steady-state metabolite analysis on cultured cells**”.** To study flux through *de novo* pyrimidine biosynthesis and pyrimidine degradation, SU-DIPG13 and BT954 cells plated overnight were collected, washed (2 times with glutamine free media), and re-plated in glutamine free media that was supplemented with either 2 mM ^15^N-glutamine (Cambridge isotope laboratories) or 2 mM GlutaMAX (represents the 0 hours timepoint of ^15^N-glutamine tracing). Samples were collected and processed after 6, 12, or 18 hours of exposure to ^15^N-glutamine-containing media. Cells growing in media without ^15^N-glutamine (0 hour ^15^N-glutamine tracing timepoint) were collected alongside the samples collected for the 18 hour timepoint.

Dried pellets of ^15^N-glutamine tracing samples were shipped over dry ice to Dr. Samuel K. McBrayer’s laboratory at University of Texas Southwestern Medical Center. For LC-MS analyses, samples were analyzed as previously described (Faubert et al., 2021; Tasdogan et al., 2020). Dried metabolites were resuspended in 30-60 µl 80% acetonitrile. Resuspended samples were vortexed for 20 minutes at 4°C, then centrifuged at 21,100 × g for 10 minutes at 4°C. 10 µl of the supernatant was injected and analyzed with a Q-Exactive HF-X or Orbitrap Exploris hybrid quadrupole-orbitrap mass spectrometer (ThermoFisher) coupled to a Vanquish Flex UHPLC system (ThermoFisher). Chromatographic resolution of metabolites was achieved on a Millipore ZIC-pHILIC column using a linear gradient of 10 mM ammonium formate (pH 9.8) and acetonitrile. Spectra were acquired with a resolving power of either 120,000 or 240,000 full width at half maximum, a scan range set to either 80-1,200 m/z or 70-1,050 m/z, and polarity switching. Data-dependent MS/MS data was acquired on unlabeled samples to confirm metabolite identity when necessary. Peaks were integrated using El-Maven 0.12.0 software (Elucidata). Total ion counts were quantified using TraceFinder 5.1 SP2 software (ThermoFisher). Peaks were normalized to total ion counts using the R statistical programming language. For stable isotope tracing studies, correction for natural abundance of metabolite labeling was performed using the AccuCor package (version 0.2.3) in the R statistical programming language (Su et al., 2017).

### Cell cycle, apoptosis, and γ-H2AX detection assays

To precisely quantify S-phase cell population, 1×10^5^ cells were plated in 2 ml media overnight and treated with 1.25 nM BAY2402234 for 24 hours before cells were incubated in the presence of BrdU (10 µM) for 45 minutes and harvested for analysis of BrdU positive cells using the BrdU flow kit according to manufacturer’s protocol (BD Biosciences). Nuclei were stained with 7-AAD. Data were collected on CytoflexS and analyzed by FlowJo software.

To detect and measure apoptotic cell populations, cleaved CASPASE3 (c-CASP3) was measured by flow cytometry and analyzed by FlowJo software. 1 × 10^5^ cells were treated with 1.25 nM BAY2402234 for 48 hours before cells were harvested and fixed in 4% paraformaldehyde for 15 minutes. Cells were then permeabilized and re-fixed in cold 90% methanol overnight. Cells were washed on the following day, blocked in 1X PBS containing 1 mg/ml BSA and 2.5 µl of human FcX-TruStain (BioLegend Inc.) for 15 minutes and incubated with PE-c-CASPASE3 antibody (Cell Signaling #9978) for 1 hour before cells were washed, nuclei stained with Hoechst 33342 and data collected on CytoflexS.

To detect apoptosis following shRNA mediated *CAD* or *DHODH* knockdown, Annexin V staining was performed as per manufacturer’s instructions (Biolegend Inc.). Briefly, 1×10^6^ cells were infected with *CAD*, *DHODH,* or control scrambled (scr) shRNA lentiviral particles in the presence of 8 µg/ml polybrene for 14-16 hours before cells were washed and replated in media. After 48 hours, cells were harvested, washed with cold 2 ml staining buffer (BioLegend Inc, catalog # 420201) and resuspended in cold Annexin V binding buffer at 1×10^6^ cells per ml. Approximately 0.1×10^6^ cells were incubated with 5 µl FITC-Annexin V for 15 minutes on ice in the dark. Nuclei were stained with propidium iodide (1 µg/ml) for 5 minutes on ice in the dark, before 400 µl of Annexin V Binding Buffer was added to each tube and data were immediately collected by flow cytometry on CytoflexS.

To detect DNA damage, γ-H2AX-positive cells were determined after 48 hours of exposure to 1.25 nM BAY2402234 using the fixation, permeabilization, blocking, and staining protocols as described for c-CASP3 staining except FITC-γ-H2AX antibody (BioLegend Inc.) was used to detect DNA damage and data were collected by flow cytometry as previously described (Pal et al., 2018). Nuclei was stained with Hoechst 33342.

For all flow cytometry-based assays, data analysis was performed using FlowJo. Cells were gated on SSC-A/FSC-A and singlet discrimination (DNA stain-A/DNA stain-H) was performed before assessment of antigen staining (BrdU, c-CASP3, Annexin V or γ-H2AX).

### Establishment of modified cell lines

To derive a BAY2402234-resistant cell line, DIPG1 cells were grown in BAY2402234, with doses doubling every 2 weeks (starting dose 0.3 nM, final dose 2.5 nM), then maintained in 0.5 nM BAY2402234, but cultured for 1-4 days without BAY2402234 before experiments executed.

To establish, luciferized DIPG1; DIPG1 and SU-DIPG4 cells (expressing doxycycline inducible shRNA against *DPYD)*; and BT954 and BT333 cells (overexpressing flag-tagged DPYD or empty vector), cells were transduced with respective lentivirus for 24 hours, before cells were washed and re-plated in media without lentiviral particles (Morgenstern and Land, 1990; Ni et al., 2016; Shaul et al., 2014). Puromycin selection was begun 48 hours later: DIPG1 (80 µg/ml), SU-DIPG4 (2 µg/ml), BT954 (1 µg/ml), and BT333 (25 µg/ml); Blasticidin selection was also started 48 hours later (2 µg/ml) for luciferized DIPG1. Cells were grown in the presence of the antibiotics for at least 2 weeks before being used for experiments as needed.

### Western blotting and immunofluorescence

10×10^6^ cells were treated with DMSO, BAY2402234, elimusertib, or HU for 24 hours before whole cell lysates were prepared in UTB buffer (50 mM Tris-HCl pH 7.5, 8M Urea, 150 mM β-mercaptoethanol), for the analysis of DNA/ chromatin bound proteins in Figures 5, 7 and S5, supplemented with complete mini protease inhibitor cocktail (Roche) and phosphatase inhibitor cocktail (Roche) and sonicated. For detection of cytoplasmic proteins as in Figures 4 and S4, whole cell lysates were prepared in IP buffer (20 mM Tris-HCl [pH8.0], 300 mM NaCl, 1 mM EDTA, 0.5% NP40) supplemented with the mini protease inhibitor cocktail and phosphatase inhibitor cocktail. Whole cell lysate (25-60 mg) was separated by SDS-PAGE, transferred to PDVF membrane, blocked (5% non-fat milk in Tris buffered saline except for detection of phosphorylated proteins wherein 5% BSA was used), and incubated with primary antibody at recommended dilution overnight before secondary antibody incubation and detection with ECL reagent. Primary antibodies are listed in the Key resource table.

For detection of chromatin-bound RPA by immunofluorescence staining, cells were plated overnight, then treated with BAY2402234, elimusertib, HU, or DMSO for 24 hours. Cells were harvested and pre-extracted in 1X PBS containing 0.5% Triton-X100 on ice for 10 minutes before being washed and fixed in 4% paraformaldehyde for 15 minutes. Cells were blocked in 1X PBS containing 3% goat serum, 0.1% Triton X-100, 1 mmol/L EDTA, and 1 mg/mL BSA for 1 hour and cytospun onto super-frosted slides. Slides were dried for 10 minutes and then incubated with primary antibody (RPA2 at 1:200, ab2175, Abcam Inc.; γ-H2AX at 1:1000, 05-636, Millipore) in blocking buffer overnight at 4°C. After overnight incubations, slides were washed prior to incubation with fluorophore conjugated secondary antibody solution for 1 hour. Slides were washed, nuclei counterstained with DAPI and mounted. Images were taken at 63X magnification using the Zeiss microscope and number of RPA foci and γ-H2AX staining intensity per nucleus were determined using CellProfiler software (Carpenter et al., 2006) for at least 100 cells per condition. The Dapi stain was first used to identify nuclei and then the number of foci within the nuclei were counted or the signal intensity for γ-H2AX within the nuclei was determined.

### DNA fiber assay

To study impacts on replication fork progression, replicating DNA was labeled and analyzed by DNA fiber assays. Briefly, after 24 hours of DMSO or BAY2402234 treatment, cells were pulsed with 50 mM of CldU (Sigma-Aldrich, C-6891) for 30 minutes, washed and again pulsed with 250 mM of IdU (Sigma-Aldrich, I-7125) for 30 minutes followed by washing and accutase treatment to get single cell suspensions. Next, 2×10^4^ cells were embedded in agarose plugs and processed using the FiberPrepKit (Genomic Vision, EXT-001) according to manufacturer’s instructions. The plugs were treated with protease overnight at 50°C, washed extensively, agarose was melted and digested overnight with β-agarose at 42°C before DNA solutions were combed on coated coverslips (Genomic Vision) using the FiberComb Molecular Coming System; coverslips were dehydrated for 2 hours at 60°C and stored at -20°C until immunodetection. Next, coverslips were subjected to chemical denaturation in fresh solution containing 0.5M NaOH and 1M NaCl for 8 minutes at room temperature, washed and dehydrated sequentially in 70%, 90%, and 100% ethanol bath for 3 minutes each.

Immunodetection of the replication tracks was performed using primary antibodies diluted in 1% BSA [Rat anti-CldU (Abcam, ab6326); mouse anti-IdU (BD Biosciences, 347580)] for 110 minutes at 37°C followed by washing and secondary antibodies goat anti-rat Cy5 (Abcam, ab6565) and goat anti-mouse Cy3.5 (Abcam, ab6946) for 45 minutes at 37°C. To visualize single stranded DNA (ssDNA), coverslips were also stained with mouse anti-ssDNA (DSHB University of Iowa) for 75 minutes at 37°C and detected with goat anti-mouse BV480 (BD Biosciences, 564877) for 45 minutes at 37°C. During immunodetection all washes (repeated three times) were performed with 1X PBS containing 0.05% Tween-20 for 3 minutes. Finally, coverslips were dehydrated in 70%, 90%, and 100% ethanol bath for 3 minutes each, dried, mounted on slides (Genomic Vision) and scanned using the Fiber Vision automated scanner (Genomic Vision). The total length of IdU and the CldU labeled tracks were measured for at least 95 DNA fibers using ImageJ software and to determine the fork speed (kbp/min), total length was divided by IdU and CldU incorporation time (60 minutes) and multiplied by a factor of 2.59 (Jackson and Pombo, 1998).

### Metaphase spreads

To prepare metaphase spreads, cell cultures treated with DHODHi and/or ATRi for 24 hours were harvested and washed twice in 20-fold excess media to remove drugs before re-plating in drug-free fresh media. After 3-4 hours, 30 ng/mL of Colcemid (Roche) was added and cultures were incubated for an additional 17 hours to arrest cells at metaphase. Cells were harvested and treated with accutase to obtain single cell suspension. Next, cells were collected by centrifugation and treated with warm hypotonic solution (0.075M KCl) for 20 minutes at 37°C.

Following hypotonic treatment, fresh fixative (3-parts methanol:1-part acetic acid) was added to the cells for 10 minutes at room temperature. The fixing step was repeated three times. To make slides, cells were resuspended in a fresh fixative solution of methanol and acetic acid, dropped on a pre-cleaned slide, and stained with Giemsa stain. Analysis was performed on a brightfield microscope at 100X magnification. Mitotic abnormalities were visually scored and at least 25 metaphase spreads were scored for each condition.

### Establishment of orthotopic tumor in mice

All animal studies were performed according to Dana-Farber Cancer Institute Institutional Animal Care and Use Committee (IACUC)-approved protocols. Animals were injected intraperitoneally with the analgesic buprenorphine 0.05 mg/kg and then anesthetized with isoflurane 2–3% mixed with medical air and placed on a stereotactic frame. Next, a small incision and a small burr hole was made with a 25-gauge needle and SU-DIPG13-P* or DIPG1 cells expressing luciferase gene (1×10^5^ cells for SU-DIPG13P* or 5×10^4^ cells for DIPG1; in 3 μl PBS) were injected stereo-tactically into the right pons (stereotactic coordinates zeroed on bregma: -1.5 mm X(ML), -5.5 mm Y(AP) and -5.0 mm Z(DV)) of 5 weeks-old female NSG mice (NOD.Cg-Prkdcscid Il2rgtm1Wjl/SzJ, The Jackson Laboratory, Bar Harbor, ME) at rate of 1 µl/min with use of an infusion pump before the incision was closed. Mice were then checked daily for signs of distress, including seizures, weight loss, or tremors, and euthanized as they developed neurological symptoms, including head tilt, seizures, sudden weight loss, loss of balance, and/or ataxia. To monitor the ability of *CAD* or *DHODH* knockdown DMG cells to establish orthotopic tumors, luciferized DIPG1 cells were transduced with lentivirus expressing the gene specific shRNA or control scrambled shRNA for 16 hours, washed and grown in virus free media for 24 hours before they were implanted in mouse brainstem as described above.

### Bioluminescence imaging and *in vivo* treatment

Tumor growth was monitored weekly using the IVIS Spectrum In Vivo Imaging System (PerkinElmer), starting at day 8 post-cell injections. Briefly, mice were injected subcutaneously with 75 mg/kg D-luciferin potassium salt (Promega E1605) in sterile PBS and anesthetized with 2% isoflurane in medical air. Serial bioluminescence images were acquired using automated exposure set-up. Peak bioluminescence signal intensity within selected regions of interest was quantified using the Living Image Software (PerkinElmer) and expressed as photon flux (ph/sec/cm^2^/sr). Mice were treated orally with BAY2402234 (4 mg/Kg) or vehicle once daily until euthanasia for efficacy studies and for 4 consecutive days for the pharmacodynamic study before mice were euthanized and brain tissues were harvested. BAY2402234 was formulated in 90% PEG400 and 10% ethanol and administered by oral gavage.

### Tumor and tissue analysis by MALDI mass spectroscopy and microscopy

Tissue preparation for MALDI MSI and microscopy: Mouse brains were dissected, snap-frozen in liquid nitrogen, and stored at -80^0^C. Sagittal cryosections of 10 µm thickness were collected from each mouse brain and thaw-mounted onto indium tin oxide (ITO) slides. Whole healthy mouse control brain tissues were homogenized and spiked with BAY2402234 concentrations ranging from 0.2-50 µM. The nine spiked homogenates were then dispensed into a nine-well gelatin (40%) tissue microarray array (TMA) mold with 1.5 mm core diameter channels to create a quantitative tissue mimetics array. The tissue mimetics were frozen and cryo-sectioned at the same thickness as the mouse brain tissues and underwent the same sample preparation. Serial sections were obtained for MALDI MSI, hematoxylin and eosin (H&E), γ-H2AX immunostaining and DAPI staining. Optical and fluorescence microscopy images were acquired using 10X and 40X objectives (Zeiss Observer Z.1, Oberkochen, Germany) and a FITC filter.

MALDI Matrix preparation and application: For BAY2402234 tissue quantification, a 2,5-dihydroxybenzoic acid (160 mg/mL) matrix solution dissolved in 70:30 methanol: 0.1% TFA with 1% DMSO was used. The matrix was applied onto tissue sections using a TM sprayer (HTX Technologies, Chapel Hill, NC) with a two-pass cycle at a flow rate (0.18 mL/min), spray nozzle velocity (1200 mm/min), nitrogen gas pressure (10 psi), spray nozzle temperature (75°C), and track spacing (2 mm). A recrystallization step was performed with 5% acetic acid solution at 85°C for 6 minutes. Dihydroorotate, N-carbamoyl-L-aspartate, and uridine monophosphate (UMP) were imaged using a 1,5-diaminonaphthalene hydrochloride (4.3 mg/mL) matrix solution in 4.5/5/0.5 HPLC grade water/ethanol/1 M HCl (v/v/v). The TM-sprayer parameters included a four-pass cycle with a flow rate (0.09 mL/min), spray nozzle velocity (1200 mm/min), spray nozzle temperature (75°C), nitrogen gas pressure (10 psi), track spacing (2 mm).

MALDI MRM mass spectrometry imaging: Tissue and tissue mimetic sections were imaged using a timsTOF fleX mass spectrometer (Bruker Daltonics, Billerica, MA) operating in positive ion mode by multiple reaction monitoring (MRM) scanning between *m/z* 100-650. A BAY2402234 solution was infused through the ESI source to optimize the MRM settings, ion transfer funnels, quadrupole, collision cell, and focus pre-TOF parameters. The optimal collision energy for the BAY2402234 precursor was 35 eV with a 3 *m/z* isolation width for the precursor to product ion transition 521.101➔376.091 corresponding to [C_21_H_18_ClF_5_N_4_O_4_+H]^+^ and [C_15_H_12_F_4_N_3_O_4_+H]^+^, respectively (SI Fig 1. a). The ESI method was then transferred to a MALDI source method and tuned using an Agilent tune mix solution (Agilent Technologies, Santa Clara, CA). For the MALDI MSI conditions, the laser repetition rate was set to 10,000 Hz, with 2,000 laser shots per 50 µm pixel size. Data visualization using SCiLS Lab software (version 2021a premium, Bruker Daltonics, Billerica, MA) was used without data normalization. A linear regression correlating the ion intensity and BAY2402234 concentration range between 0.2-2 µM from the tissue mimetic was established with a correlation coefficient of 0.994 (SI Fig 1. bc). Limit of detection (LOD) of 0.11 µM (S/N ratio of >3) and limit of quantification (LOQ) of 0.35 µM (S/N ratio of >10) were calculated.

Metabolite MALDI MSI: Dihydroorotate, N-carbamoyl-L-aspartate, and UMP were imaged from the BAY2402234 dosed and vehicle tissue sections with the Q-TOF instrument operated in negative ion mode in full scan mode for *m/z* 50-400. Standard solutions were infused using the ESI source to optimize the instrument parameters. Dihydroorotate and N-carbamoyl-L-aspartate fragmented in full MS scan mode, so their product ions were directly monitored in MS mode by setting the collision energy to 10 eV. The product ion monitored for dihydroorotate was *m/z* 113.037 [C_4_H_6_N_2_O_2_-H]^-^ and for N-carbamoyl-L-aspartate *m/z* 132.031 [C_4_H_7_NO_4_-H]^-^ (SI Fig 2. b). Using a serial section, a full scan approach was used to image UMP by monitoring its precursor ion at *m/z* 323.029 [C_9_H_13_N_2_O_9_P-H]^-^. All metabolites were confirmed using direct on-tissue MSMS and compared to standards under the same analytical conditions. The metabolite fold changes are calculated by dividing the average MSI intensity of all pixels per tumor ROI (region of interest) for treatment over vehicle followed by a log2 transformation.

## QUANTIFICATION AND STATISTICAL ANALYSIS

Data analysis and visualization was performed with PRISM 9 (GraphPad software). Unless indicated, all data are represented as mean ± SEM and plotted using PRISM software. The statistical test applied is indicated in the figure legends. IC50 for drug responses were calculated using the PRISM software. For two group comparisons, the t-test was used. Whenever multiple groups were compared, one-way ANOVA was used, and Tukey test was performed to correct for multiple comparisons in PRISM. For datasets where multiple t-test were performed across two groups e.g for every drug dose as in figure 4 B and D, unpaired t-test with multiple comparisons based on false discovery rate using two-stage step-up (Benjamini, Krieger, and Yekutieli) was used in PRISM. For datasets that failed normality test, we performed nonparametric Kruskal-Wallis testing with Dunn’s test to correct for multiple comparisons in PRISM. Asterisks indicate statistically significant (*, p<0.05; **, p<0.01; ***, p<0.001; ****, p<0.0001) values.

## SUPPLEMENTAL INFORMATION

Table S2. DMG dependency genes identified in each DMG line using CRISPR screen. Related to Figure 1.

Table S4. Gene list defined as pan-cancer essential genes. Related to STAR Methods.

