## Supplemental Figures and Tables for "A druggable addiction to *de novo* pyrimidine biosynthesis in diffuse midline glioma"

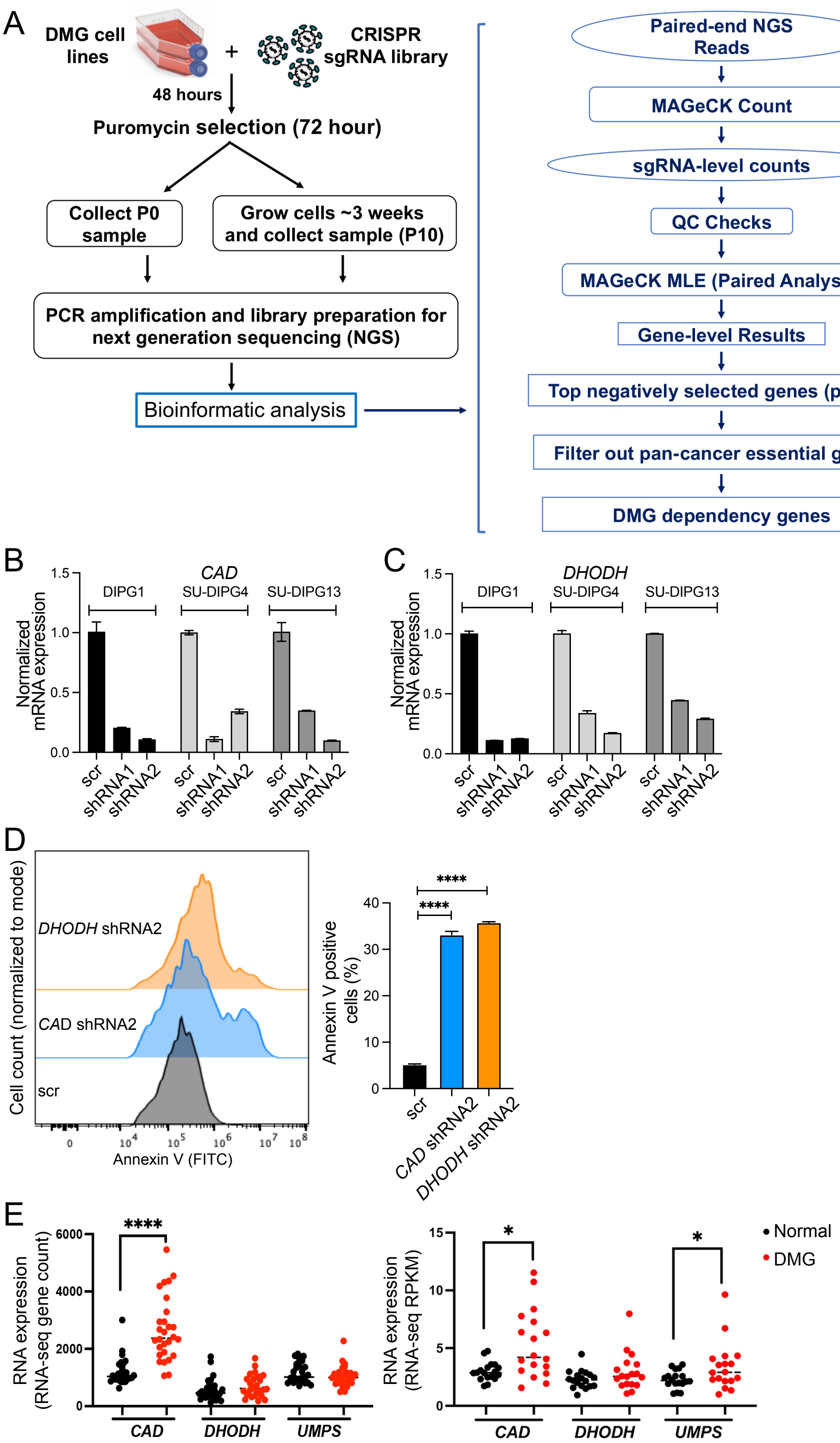

A
DMG cell lines
CRISPR sgRNA library
48 hours
Puromycin selection (72 hour)
Collect P0 sample
Grow cells ~3 weeks and collect sample (P10)
PCR amplification and library preparation for next generation sequencing (NGS)
Bioinformatic analysis
Paired-end NGS Reads
MAGeCK Count
sgRNA-level counts
QC Checks
MAGeCK MLE (Paired Analysis)
Gene-level Results
Top negatively selected genes (p<0.01)
Filter out pan-cancer essential genes
DMG dependency genes
B
CAD
DIPG1
SU-DIPG4
SU-DIPG13
Normalized mRNA expression
scr
shRNA1
shRNA2
C
DHODH
D
DHODH shRNA2
CAD shRNA2
scr
Cell count (normalized to mode)
Annexin V (FITC)
Annexin V positive cells (%)
****
E
RNA expression (RNA-seq gene count)
RNA expression (RNA-seq RPKM)
CAD
DHODH
UMPS
Normal
DMG

**Figure S1. CRISPR screen identifies *de novo* pyrimidine synthesis as a dependency in DMG. Related to Figure 1.**

(A) Flow chart of CRISPR screen workflow in DMG cell lines. Two hundred million cells from each of three DMG cell lines (SF8628, DIPG1, and SU-DIPG13) were infected with a human lentiviral library of sgRNA; 48 hours after infection, cells expressing sgRNA were selected with puromycin (for 72 hours) with subsequent steps, as indicated. Two independent CRISPR screen experiments were performed in each line and analyzed using MAGeCK software.

(B and C) shRNA knockdown of *CAD* and *DHODH* expression in DMG cells. mRNA expression was determined by reverse transcription-quantitative PCR assays (RT-qPCR) at 72 hours after infection of DIPG1, SU-DIPG4, or SU-DIPG13 with lentivirus expressing shRNA targeting either (B) *CAD or* (C) *DHODH.* Two distinct shRNAs were used for each targeted gene. 18S was used for normalization and data are represented relative to scrambled (scr) control shRNA.

(D) SU-DIPG13 cells were infected with lentivirus expressing shRNA targeting either *CAD* or *DHODH* and induction of apoptosis was measured by Annexin V staining followed by flow cytometry analysis. Scr shRNA served as control. Representative overlay histograms of Annexin V staining (left panel) and quantification of Annexin V positive cells (n=3; right panel). Unpaired t test, **** $p<0.0001$.

(E) Expression of *de novo* pyrimidine synthesis genes in DMG patient tumors (red) and matched normal tissues (black) from two published datasets.  Zhu et al. left panel (Zhu et al., 2021) and Berlow et al. right panel (Berlow et al., 2018). Paired t-test, * $p<0.05$, **** $p<0.0001$.

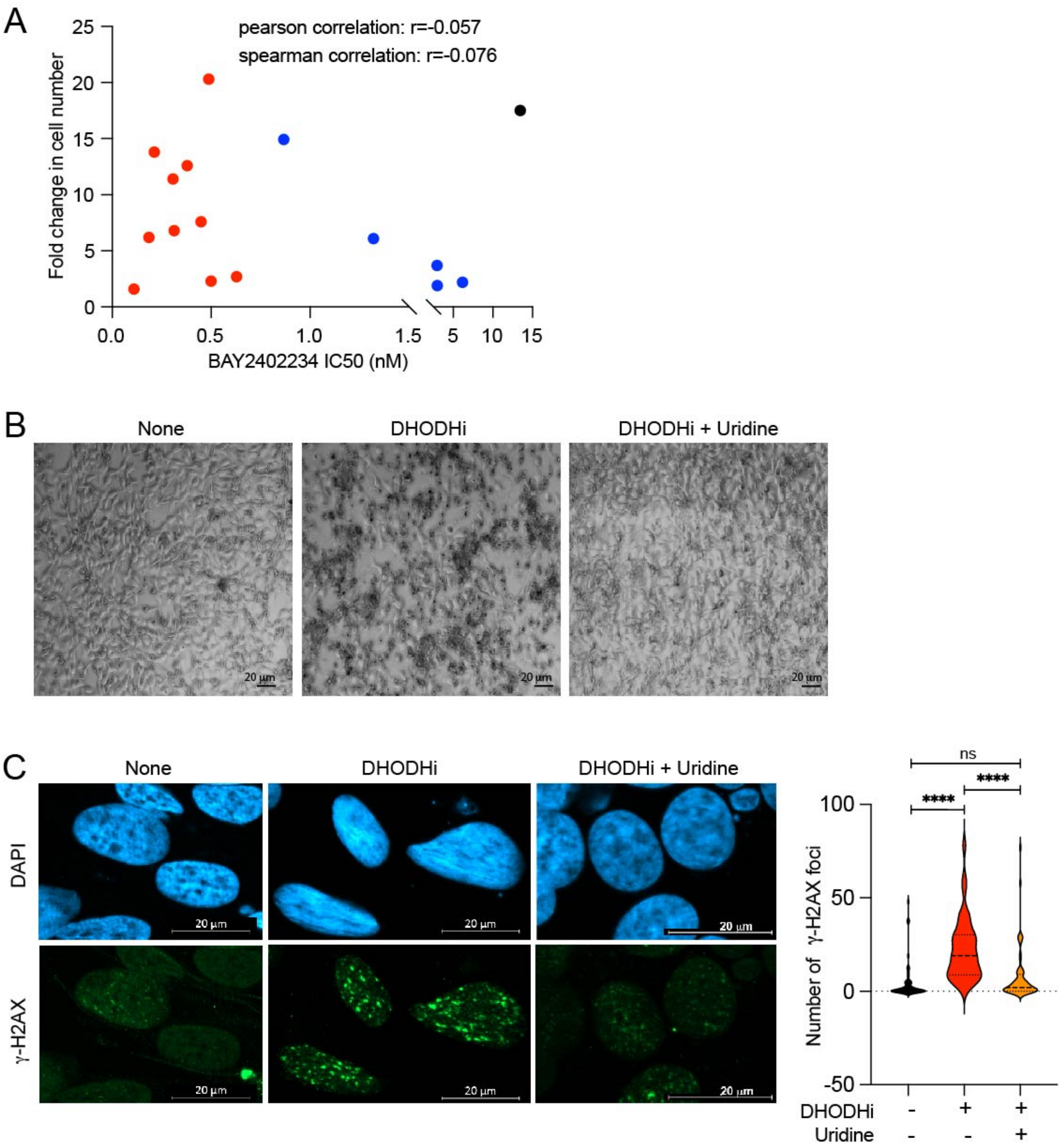
A
pearson correlation: r=-0.057
spearman correlation: r=-0.076
Fold change in cell number
BAY2402234 IC50 (nM)
B
None
DHODHi
DHODHi + Uridine
20 µm
C
None
DHODHi
DHODHi + Uridine
DAPI
γ-H2AX
20 µm
ns
****
****
Number of γ-H2AX foci
DHODHi
Uridine

**Figure S2. BAY2402234 inhibits DMG cell proliferation and induces DNA damage, a phenomenon not correlated with proliferation rate. Related to Figure 2.**

(A) Lack of correlation between cell proliferation and BAY2402234 sensitivity. Scatter plot of cell proliferation (fold change in cell number after 5 days in culture) versus IC50 of BAY2402234-treated cells (5 days of treatment) was analyzed by both Pearson and Spearman correlations, with no correlation observed.

(B) Representative images of SU-DIPG4 with or without BAY2402234 (DHODHi; 1.25 nM, 72 hours), with or without uridine supplementation (100 μM). Scale bar is 20 μm.

(C) Immunofluorescence staining of γ-H2AX in SU-DIPG4 with or without BAY2402234 (DHODHi; 1.25 nM, 48 hours), with or without uridine supplementation (100 μM). Representative images in upper panel (scale bar is 20 μm); quantification of γ-H2AX foci shown as violin plots in lower panel. ANOVA, **** $p<0.0001$, ns-not significant.

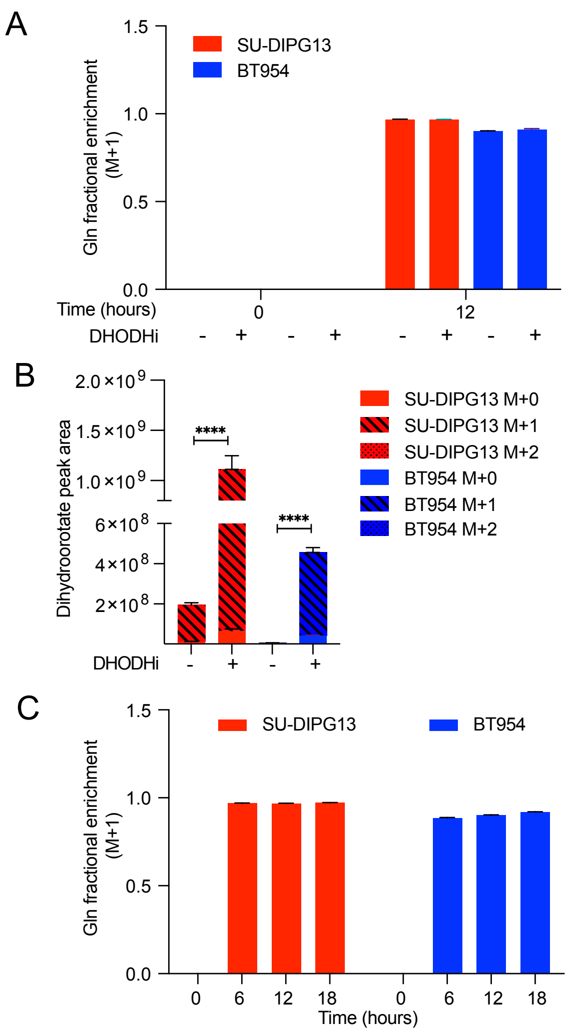

A
Gln fractional enrichment (M+1)
1.5
1.0
0.5
0.0
SU-DIPG13
BT954
Time (hours)
0
12
DHODHi
- + - + - + - +
B
Dihydroorotate peak area
2.0×10^9
1.5×10^9
1.0×10^9
6×10^8
4×10^8
2×10^8
****
****
SU-DIPG13 M+0
SU-DIPG13 M+1
SU-DIPG13 M+2
BT954 M+0
BT954 M+1
BT954 M+2
DHODHi
- + - +
C
Gln fractional enrichment (M+1)
1.5
1.0
0.5
0.0
SU-DIPG13
BT954
0 6 12 18 0 6 12 18
Time (hours)

**Figure S3. Experimental controls documenting efficient $^{15}$N-glutamine uptake and inhibition of *de novo* pyrimidine biosynthesis by BAY2402234. Related to Figure 3.**

(A) Fractional enrichment of $^{15}$N-glutamine (M+1) in SU-DIPG13 (DMG) and BT954 (aGBM) following 0 or 12 hours of exposure to media containing $^{15}$N-glutamine with or without BAY2402234 (DHODHi; 0.5 nM, 28 hours).

(B) Peak area of unlabeled (M+0) and labelled $^{15}$N-dihydroorotate (M+1) in SU-DIPG13 (DMG) and BT954 (aGBM) following 12 hours of exposure to media containing $^{15}$N-glutamine with or without BAY2402234 (DHODHi). For DHODHi samples, cells were pretreated with BAY2402234 for 16 hours prior to switching to media containing $^{15}$N-glutamine. **** $p<0.0001$ for the comparisons of indicated M+1 peak areas, ANOVA with Tukey's multiple comparison test.

(C) Time course of fractional enrichment of $^{15}$N-glutamine (M+1) in SU-DIPG13 (DMG) and BT954 (aGBM) exposed to media containing $^{15}$N-glutamine for indicated times.

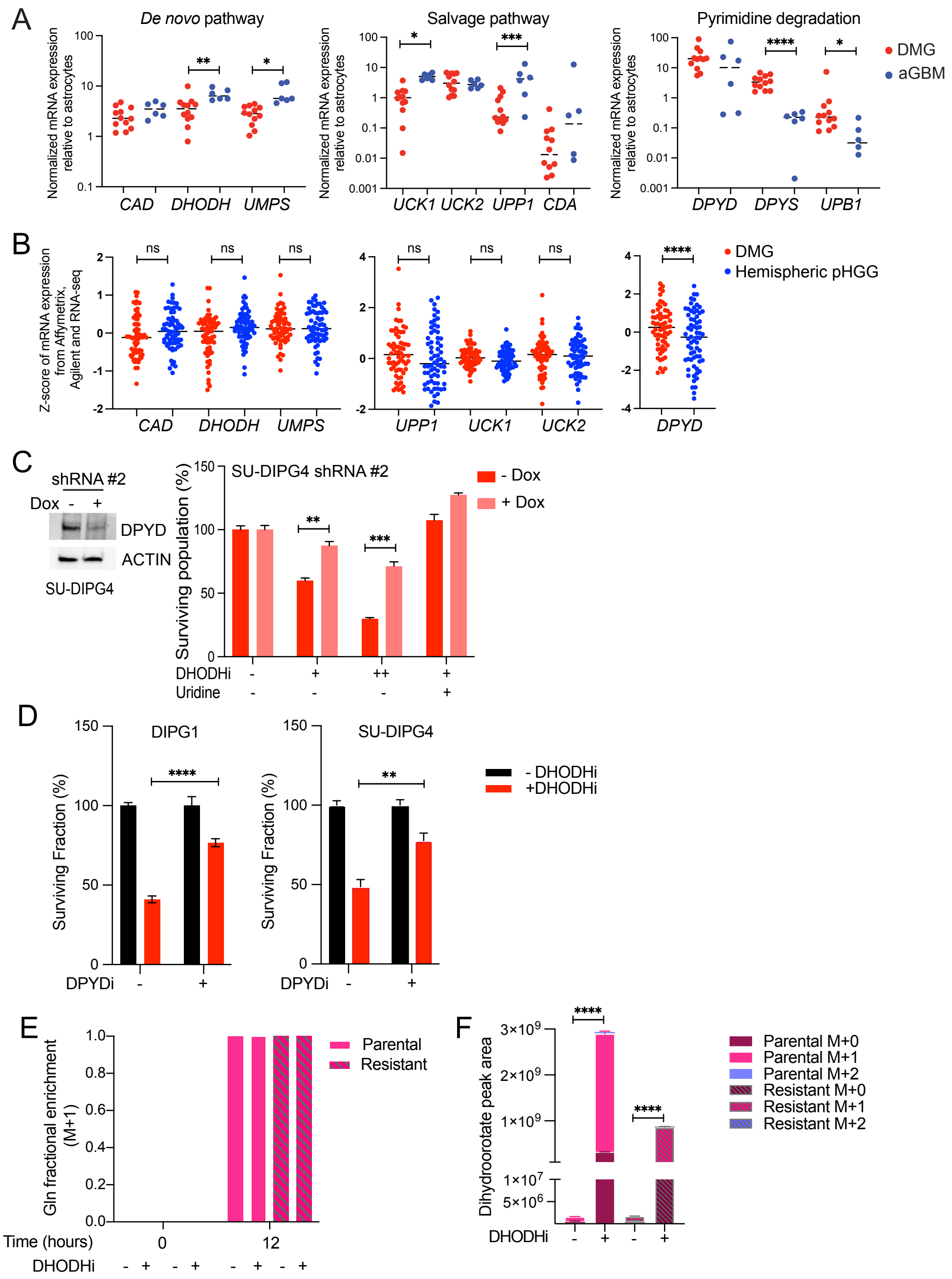
A
De novo pathway
Salvage pathway
Pyrimidine degradation
Normalized mRNA expression relative to astrocytes
CAD DHODH UMPS
UCK1 UCK2 UPP1 CDA
DPYD DPYS UPB1
DMG
aGBM
B
Z-score of mRNA expression from Affymetrix, Agilent and RNA-seq
ns
****
CAD DHODH UMPS
UPP1 UCK1 UCK2
DPYD
DMG
Hemispheric pHGG
C
shRNA #2
Dox - +
DPYD
ACTIN
SU-DIPG4
SU-DIPG4 shRNA #2
Surviving population (%)
- Dox
+ Dox
DHODHi
Uridine
D
DIPG1
SU-DIPG4
Surviving Fraction (%)
- DHODHi
+DHODHi
DPYDi
E
Gln fractional enrichment (M+1)
Parental
Resistant
Time (hours)
DHODHi
F
Dihydroorotate peak area
Parental M+0
Parental M+1
Parental M+2
Resistant M+0
Resistant M+1
Resistant M+2
DHODHi

**Figure S4: Pyrimidine degradation enhances BAY2402234 sensitivity. Related to Figure 4.**

(A) mRNA expression of genes involved in pyrimidine homeostasis in DMG (red) and aGBM (blue) cell lines, quantified by RT-qPCR in triplicates and presented relative to normal astrocytes. Genes in *de novo* synthesis (left), salvage (middle), and pyrimidine degradation (right) are shown. 18S was used for normalization. Unpaired t-test, * p<0.05, ** p<0.01, *** p<0.001, **** p<0.0001.

(B) RNA expression of genes belonging to pyrimidine biosynthesis in the *de novo* (left) and salvage (middle) pathways as well as *DPYD*, a pyrimidine degradation gene, in patient-derived primary newly diagnosed (pre-treatment) tumors from DMG or pediatric hemispheric high-grade glioma patients. Data as z-scores were extracted from PedcBioPortal dataset (ICR London, 2017) (Mackay et al., 2017). Unpaired t-test, **** p< 0.0001, ns-not significant.

(C) Western blot analyses of extracts from SU-DIPG4 cells expressing doxycycline-inducible shRNA #2 against *DPYD* with or without doxycycline (Dox; 500 ng/ml) treatment for 72 hours (left panel). DMG cells as in left panel were treated with BAY2402234 (+0.5 nM, ++1 nM), with or without uridine supplementation (100 $\mu$M), for 72 hours and surviving populations quantified. Unpaired t-test, ** p<0.01, *** p<0.001.

(D) Inhibition of DPYD, an enzyme that catalyzes the first step of uracil degradation, by gimeracil (DPYDi; 1.5 $\mu$M) blunts BAY2402234 sensitivity of DMG cells (DIPG1: 1nM, 72 hours; SU-DIPG4: 0.5 nM, 48 hours). Graphs shows mean ± SEM (n=3). Unpaired t-test, ** p<0.01, **** p<0.0001.

(E) Fractional enrichment of $^{15}$N-glutamine (M+1) in parental and resistant DIPG1 following 0 or 12 hours of exposure to media containing $^{15}$N-glutamine with or without BAY2402234 (DHODHi; 1.25 nM, 26 hours). Length of BAY2402234 treatment is the same for all samples.

(F) Peak area of dihydroorotate unlabeled (M+0) and labeled ($^{15}$N-dihydroorotate, M+1) in parental and resistant DIPG1 following 12 hours of exposure to media containing $^{15}$N-glutamine with or without BAY2402234 (DHODHi; 1.25nM). For DHODHi samples, cells were pretreated with BAY2402234 for 14 hours prior to switching to media containing $^{15}$N-glutamine. Length of BAY2402234 treatment is the same for all samples. * p<0.05, **** p<0.0001 for comparisons of indicated M+1 peak areas, ANOVA with Tukey's multiple comparison test.

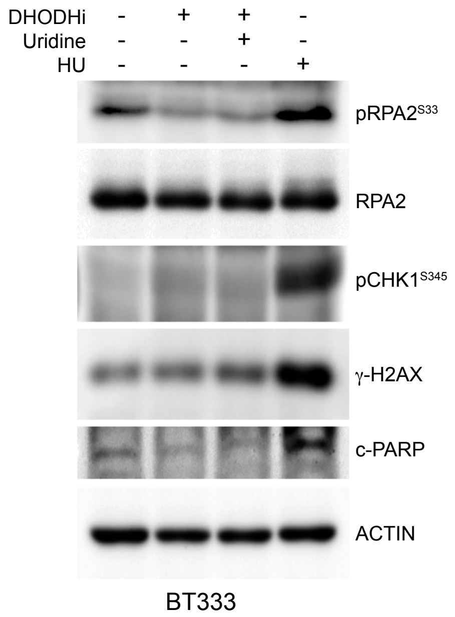


**Figure S5. BAY2402234 fails to induce replication stress in aGBM. Related to Figure 5.** Western blot analyses of extracts from BT333 cells treated with or without BAY2402234 (DHODHi; 1.25 nM, 24 hours) with or without uridine supplementation (100 μM) and evaluated for indicated proteins. pRPA2$^{S33}$: phosphorylated RPA2 at serine 33. pCHK1$^{S345}$: phosphorylated CHK1 at serine 345. c-PARP: cleaved PARP. Hydroxyurea (HU), a known inducer of replication stress, is included as a positive control.

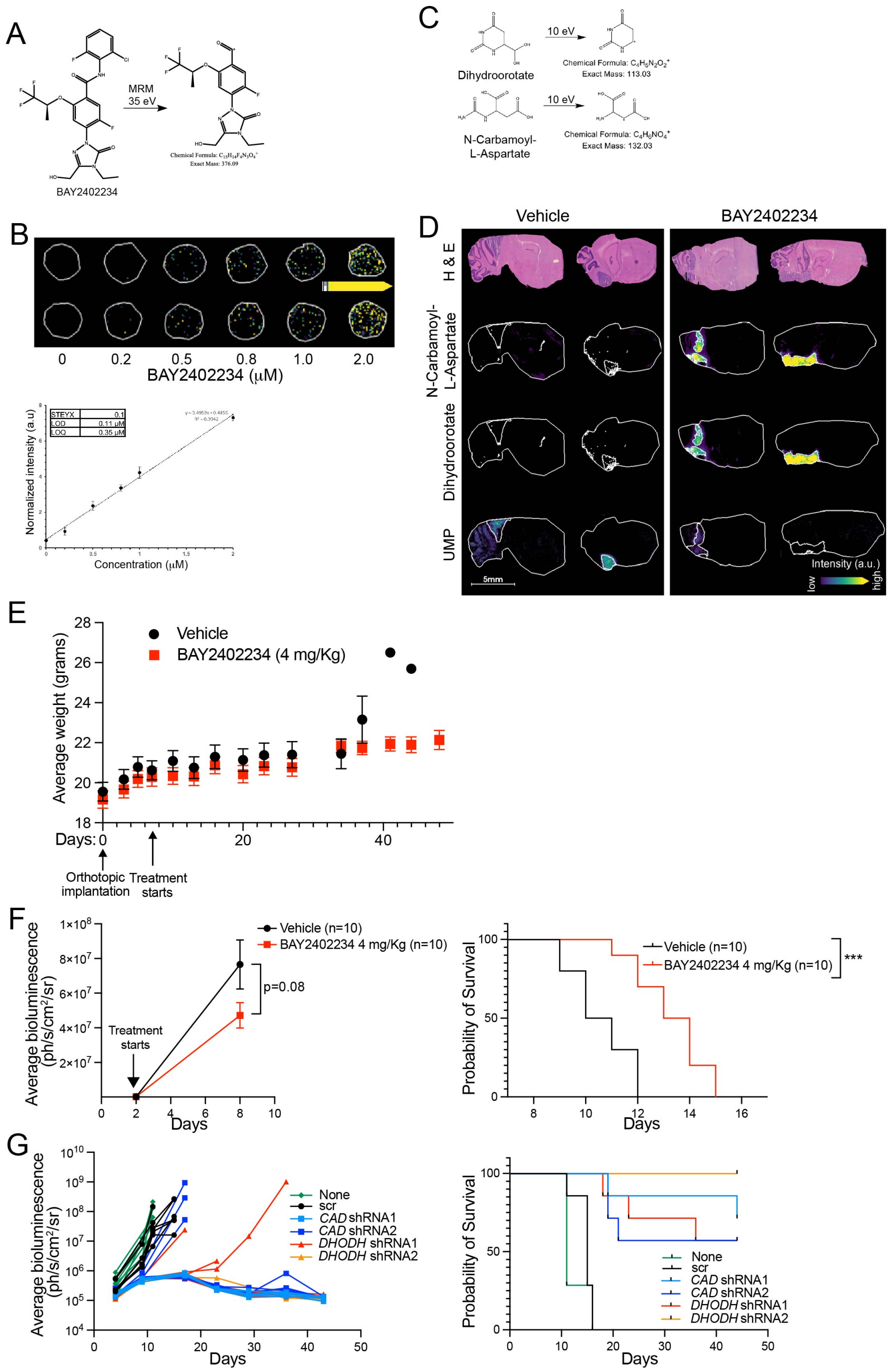
A
MRM
35 eV
BAY2402234
C
10 eV
Dihydroorotate
N-Carbamoyl-
L-Aspartate
Exact Mass: 113.03
Exact Mass: 132.03
B
0
0.2
0.5
0.8
1.0
2.0
BAY2402234 (μM)
Normalized intensity (a.u)
Concentration (μM)
STEYX 0.1
LOD 0.11 μM
LOQ 0.35 μM
D
Vehicle
BAY2402234
H & E
N-Carbamoyl-L-Aspartate
Dihydroorotate
UMP
5mm
Intensity (a.u.)
low
high
E
Vehicle
BAY2402234 (4 mg/Kg)
Average weight (grams)
Days:
Orthotopic implantation
Treatment starts
F
Vehicle (n=10)
BAY2402234 4 mg/Kg (n=10)
p=0.08
Treatment starts
Average bioluminescence (ph/s/cm²/sr)
Days
Probability of Survival
***
G
None
scr
CAD shRNA1
CAD shRNA2
DHODH shRNA1
DHODH shRNA2
Average bioluminescence (ph/s/cm²/sr)
Days
Probability of Survival

**Figure S6. Inhibition of *de novo* pyrimidine biosynthesis is well tolerated, hinders tumor growth, and improves overall survival. Related to Figure 6.**

(A) Chemical structure of precursor and product ions of BAY2402234 used for drug quantification in mouse tissues.

(B) Tissue mimetic analysis of BAY2402234 ranging from 0.2-2 μM used to construct calibration curve for BAY2402234 (right panel) MALDI MSI quantification with an $R^2$ = 0.9942.

(C) Chemical structures of dihydroorotate and N-carbamoyl-L-aspartate showing precursor and product ions that were monitored by MALDI MSI.

(D) Biological replicates (n=2) showing molecular ion images of N-carbamoyl-L-aspartate, dihydroorotate, and UMP with corresponding H&E optical microscopy images.

(E) Weight of mice bearing DIPG13-P* orthotopic tumors, treated with either vehicle or BAY2402234 (4 mg/Kg, daily by oral gavage).

(F) Mice bearing luciferized DIPG1 orthotopic tumors, treated with either vehicle or BAY2402234 (4 mg/Kg, daily by oral gavage, until euthanasia). Tumor growth monitored by bioluminescence imaging (BLI, left); data represent average BLI signal values. Kaplan-Meier curve delineating survival of mice (right).

(G) Luciferized DIPG1 transduced with either control shRNA (scr) or shRNA against *CAD* or *DHODH* (two shRNAs for each gene) were implanted in mice brainstems 48 hours after transduction (n=7 for each group). Non-transduced cells serve as controls. Tumor growth monitored by bioluminescence imaging (BLI, left) and data for each mouse are plotted. Kaplan-Meier curve delineating survival [study end day 44 (right)].

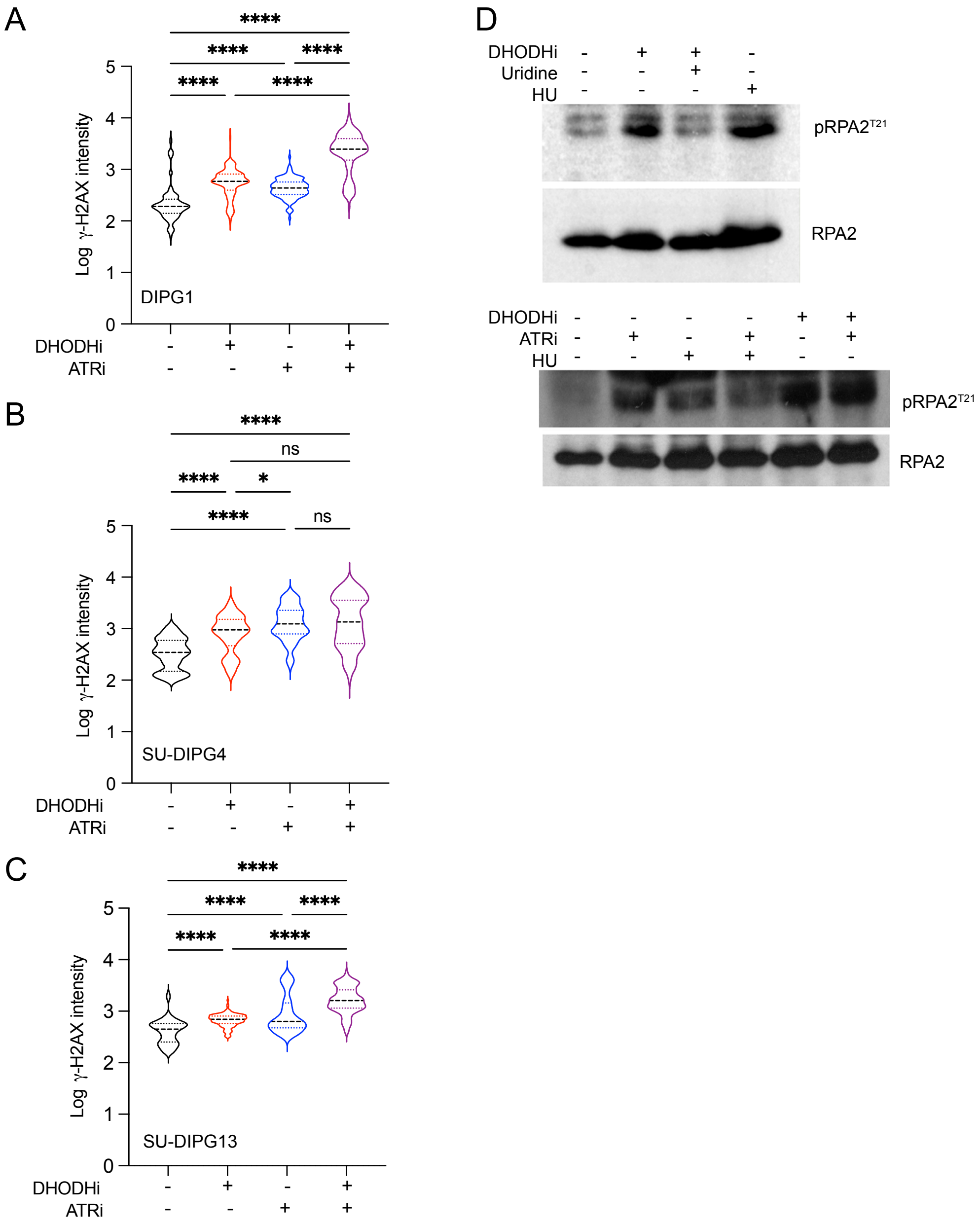
A
****
****
****
****
****
5
4
3
2
1
0
Log γ-H2AX intensity
DIPG1
DHODHi - + - +
ATRi - - + +
B
****
ns
****
*
****
ns
Log γ-H2AX intensity
SU-DIPG4
DHODHi - + - +
ATRi - - + +
C
****
****
****
****
****
Log γ-H2AX intensity
SU-DIPG13
DHODHi - + - +
ATRi - - + +
D
DHODHi - + + -
Uridine - - + -
HU - - - +
pRPA2T21
RPA2
DHODHi - - - - + +
ATRi - + - + - +
HU - - + + - -
pRPA2T21
RPA2

**Figure S7. Combination of ATR- and DHODH-inhibition augments DNA damage but does not affect pRPA2$^{T21}$ phosphorylation that is induced by replication stress in an ATR-independent fashion. Related to Figure 7.**

(A-C) γ-H2AX measured by immunostaining in DMG cells after treatment with BAY2402234 (DHODHi; 1.25 nM, 24 hours), elimusertib (ATRi; 200 nM, 24 hours), or their combination; (A) DIPG1, (B) SU-DIPG4, (C) SU-DIPG13. Intensity of γ-H2AX staining per cell determined by Cell Profiler software. ANOVA with Tukey's multiple comparison test * $p<0.05$, **** $p<0.0001$ and ns-not significant.

(D) Western blot analysis of whole cell extracts of SU-DIPG4 cells after treatment with BAY2402234 (DHODHi; 1.25 nM, 24 hours), elimusertib (ATRi; 200 nM, 24 hours), hydroxyurea (HU; 500 mM; 24 hours), or their combinations. pRPA2$^{T21}$: phosphorylated RPA2 at threonine 21.

**Table S1. Representation of pan-cancer essential genes among the top negatively selected genes required for cell viability. Related to Figure 1.**

| DMG cell line | Negatively selected genes ($p<0.01$) | Pan-cancer essential gene number | Pan-cancer essential genes (%) |
|---|---|---|---|
| SF8628 | 550 | 277 | 50.36% |
| DIPG1 | 2903 | 1340 | 46.16% |
| SU-DIPG13 | 1380 | 933 | 67.61% |

**Table S3. Common genetic alterations in DMG and aGBM lines used in this study. Related to Figure 2.**

| Cell line | Histone H3 | IDH1/2 | TP53 | PI3K pathway | ACVR1 | Other genetic alterations |
|---|---|---|---|---|---|---|
| DIPG1 (CCHMC-DIPG1) | WT | IDH1 MT*<br>IDH2 MT** | TP53 MT | MT PIK3R1, PIK3R3, PIK3CG, PIK3C2G, PIK3R6, PIK3R2, PIK3IP1 | WT | NTRK2 MT |
| CCHMC-DIPG2 | H3.3K27M | NA | NA | NA | WT | NA |
| BT869 | H3.3K27M | NA | TP53 WT | PIK3CA MT | NA | KMT2C MT PPMID MT |
| HSJD-DIPG007 | H3.3K27M | NA | TP53 WT | NA | NA | PPMID MT |
| SU-DIPG13 | H3.3K27M | NA | TP53 MT | NA | NA | NA |
| SU-DIPG17 | H3.3K27M | NA | TP53 MT | NA | NA | NA |
| SF8628 | H3.3K27M | NA | TP53 MT | NA | WT | NA |
| SU-DIPG4 | H3.1K27M | NA | TP53 MT; MDM4 amp | NA | MT | NA |
| SU-DIPG21 | H3.1K27M | NA | NA | NA | MT | NF1 MT, PDGFRA amp |
| SU-DIPG33 | H3.1K27M | NA | NA | PIK3CA MT | NA | |
| SU-DIPG36 | H3.1K27M | NA | NA | | MT | BCOR frameshift |
| BT145 | WT | WT | NA | PTEN MT | NA | CDKN2A/B deletion |
| BT189 | WT | WT | NA | PTEN biallelic inactivation | NA | CDKN2A/B deletion |
| BT333 | WT | WT | TP53 MT | NA | NA | EGFR amp CDK6 amp |
| BT424 | WT | WT | NA | | NA | NA |
| BT924 | WT | WT | NA | PIK3R1 MT | NA | NA |
| BT954 | WT | WT | NA | PIK3CA MT | NA | CDKN2A/B deletion |

WT-wild type; MT-mutated; amp-amplified; NA-information not available
* IDH1: rs34218846:V178I; **IDH2: C113R

**Table S5. Sequence of primers used for quantitative polymerase chain reaction (PCR) assays. Related to STAR methods.**

| Gene | Forward Primer | Reverse Primer |
|---|---|---|
| *18S* | 5'- GAGACTCTGGCATGCTAACTA G -3' | 5'- GGACATCTAAGGGCATCACAG -3' |
| *CAD* | 5'-TT AGCAAGTTCCTGCGAGTC-3' | 5'-CACACAGTTCTCATCCACCAT-3' |
| *DHODH* | 5'-AAGGAAACCCTAGACCCAGA-3' | 5'-GT AACCTGTGTTCCACCACT-3' |
| *UMPS* | 5'-CCAGAATTTCTTCACTTGACTCC-3' | 5'-TGCCACGACCT ACAATGATG-3' |
| *UCK1* | 5'-GTCAGGCTGTCTCGAAGAG-3' | 5'-TCAGAATGTCCTGGATGTGC-3' |
| *UCK2* | 5'-GTGGATACAGATGCGGACA-3' | 5'-GCAAGCAGAATTCCTCAAAGG-3' |
| *UPP1* | 5'-GCAGAGAAAGCTGAAAGTCAC-3' | 5'-ACACAAACTTCACATCTCCAAAC-3' |
| *CDA* | 5'-CTGGGCATCTGTGCTGAA-3' | 5'-GTGCCAAACTCTCTCATGACT-3' |
| *DPYD* | 5'-GCACACGACTCTTGGTGAG-3' | 5'-GCAGCTCCATAATAGTTCTTGTTTG-3' |
| *DPYS* | 5'-GCTGTGAACTGTCCTCTCTAC-3' | 5'-TCCAGTAGTGAGTGCCATCT-3' |
| *UPB1* | 5'-TCATAGACGCATAAAGGCTATCG-3' | 5'-CCCATCCTCTGCTGACTCA-3' |
