## Supplementary table S2 for "A druggable addiction to *de novo* pyrimidine biosynthesis in diffuse midline glioma"

Supplemental Table 1 (related to Figure 1):Gene list defined as common essential genes bas

| GeneSymbol |
|---|
| AAMP |
| AARS |
| AARS2 |
| AATF |
| ABCB7 |
| ABCE1 |
| ABCF1 |
| ABHD11 |
| ABT1 |
| ACIN1 |
| ACLY |
| ACO2 |
| ACTB |
| ACTG1 |
| ACTL6A |
| ACTR10 |
| ACTR1A |
| ACTR2 |
| ACTR3 |
| ACTR6 |
| ACTR8 |
| ADAM11 |
| ADAT3 |
| ADSL |
| AFG3L2 |
| AGAP2 |
| AGAP5 |
| AGAP6 |
| AHCTF1 |
| AHCY |
| AIFM1 |
| AK6 |
| AKIRIN1 |
| AKIRIN2 |
| ALDOA |
| ALG1 |
| ALG11 |
| ALG13 |
| ALG14 |
| ALG1L |
| ALG2 |
| ALX3 |
| ALYREF |

ANAPC1
ANAPC10
ANAPC11
ANAPC13
ANAPC15
ANAPC2
ANAPC4
ANAPC5
ANKLE2
ANLN
AP2M1
AP2S1
AQP7
AQR
ARCN1
ARF4
ARFRP1
ARGLU1
ARHGAP11B
ARIH1
ARL2
ARL4D
ARMC7
ARPC2
ARPC3
ARPC4
ASNA1
ASS1
ATF4
ATL2
ATP1A1
ATP23
ATP2A2
ATP5F1A
ATP5F1B
ATP5F1C
ATP5F1D
ATP5MC1
ATP5ME
ATP5MF
ATP5MG
ATP5MGL
ATP5PB
ATP5PD
ATP5PF

ATP5PO
ATP6AP1
ATP6AP2
ATP6V0B
ATP6V0C
ATP6V0D1
ATP6V1A
ATP6V1B2
ATP6V1C1
ATP6V1D
ATP6V1E1
ATP6V1F
ATP6V1G1
ATP6V1H
ATR
ATRIP
ATXN10
AURKA
AURKAIP1
AURKB
BAK1
BANF1
BANP
BAP1
BARD1
BCAS2
BCCIP
BCL2L1
BCLAF1
BCS1L
BDP1
BIRC5
BMS1
BNIP1
BOP1
BORA
BPTF
BRCA1
BRCA2
BRD4
BRD8
BRF1
BRF2
BRIX1
BTAF1

BTF3
BUB1
BUB1B
BUB3
BUD13
BUD23
BUD31
BYSL
C12orf10
C12orf45
C12orf66
C16orf86
C17orf49
C17orf58
C18orf21
C19orf53
C1orf109
C1orf131
C1QBP
C7orf26
C9orf78
CA5A
CACNB3
CACTIN
CADM4
CAMLG
CAP1
CAPZB
CARS
CASC3
CASTOR3
CBLL1
CBWD2
CBWD5
CCAR1
CCDC107
CCDC115
CCDC12
CCDC130
CCDC137
CCDC174
CCDC43
CCDC59
CCDC84
CCDC86

CCNA2
CCNB1
CCND1
CCNH
CCNK
CCNL1
CCT2
CCT3
CCT4
CCT5
CCT6A
CCT7
CCT8
CD2BP2
CD3EAP
CD8B
CDAN1
CDC123
CDC16
CDC20
CDC23
CDC25A
CDC26
CDC27
CDC37
CDC40
CDC42
CDC42SE1
CDC45
CDC5L
CDC6
CDC7
CDC73
CDCA5
CDCA8
CDIPT
CDK1
CDK12
CDK17
CDK2
CDK4
CDK7
CDK9
CDT1
CEBPZ

CELSR2
CENPA
CENPC
CENPE
CENPI
CENPK
CENPL
CENPM
CENPN
CENPT
CENPW
CEP131
CEP192
CEP97
CFAP20
CFAP298
CFC1
CFDP1
CFL1
CHAF1A
CHAF1B
CHCHD1
CHCHD2
CHCHD4
CHD4
CHEK1
CHERP
CHMP2A
CHMP3
CHMP4B
CHMP5
CHMP6
CHMP7
CHORDC1
CHTF8
CHTOP
CIAO1
CIAO2A
CIAO2B
CIAO3
CIAPIN1
CINP
CIR1
CIT
CITED4

CKAP5
CLASRP
CLCC1
CLEC18C
CLK2
CLNS1A
CLP1
CLSPN
CLTC
CMPK1
CMTR1
CNIH4
CNN2
CNOT1
CNOT2
CNOT3
CNOT9
COA3
COA5
COA7
COASY
COG1
COG2
COG3
COG4
COG6
COG8
COIL
COPA
COPB1
COPB2
COPE
COPG1
COPS2
COPS3
COPS4
COPS5
COPS6
COPS8
COPZ1
COQ2
COQ4
COQ5
CORO6
COX11

COX17
COX20
COX5A
COX5B
COX6A1
COX6B1
COX6C
COX7B
COX7C
CPSF1
CPSF2
CPSF3
CPSF4
CPSF6
CRCP
CRCT1
CRKL
CRLS1
CRNKL1
CS
CSE1L
CSNK1A1
CSNK2B
CSTF1
CSTF3
CTAGE9
CTCF
CTDNEP1
CTDP1
CTNNBL1
CTPS1
CTR9
CTU1
CTU2
CUL1
CUL2
CUL3
CWC22
CWC25
CWF19L2
CXCL2
CXXC1
CYC1
CYCS
CYFIP1

CYP4F2
DAD1
DAP3
DARS
DBF4
DBR1
DCAF6
DCLRE1B
DCTN1
DCTN2
DCTN3
DCTN4
DCTN5
DCTN6
DDA1
DDB1
DDN
DDOST
DDX1
DDX10
DDX11
DDX18
DDX19A
DDX20
DDX21
DDX23
DDX24
DDX27
DDX28
DDX39B
DDX3X
DDX41
DDX42
DDX46
DDX47
DDX49
DDX5
DDX51
DDX52
DDX54
DDX55
DDX56
DDX59
DDX6
DENR

DGCR8
DHDDS
DHFR
DHFR2
DHPS
DHX15
DHX16
DHX33
DHX36
DHX37
DHX38
DHX8
DHX9
DICER1
DIMT1
DIS3
DKC1
DMAP1
DNAJA1
DNAJA3
DNAJC11
DNAJC17
DNAJC8
DNAJC9
DNLZ
DNM1L
DNM2
DNMT1
DNTTIP2
DOHH
DOLK
DONSON
DPAGT1
DPF2
DPH1
DPH3
DPM2
DPY19L2
DR1
DRAP1
DROSHA
DSN1
DTL
DTYMK
DUT

DUXA
DYNC1H1
DYNC1I2
DYNLL1
DYNLRB1
E4F1
EAPP
EARS2
EBNA1BP2
ECD
ECT2
EDC4
EEF1A1
EEF1B2
EEF1G
EEF2
EEF2KMT
EFL1
EFTUD2
EHMT2
EIF1
EIF1AD
EIF1AX
EIF2B1
EIF2B2
EIF2B3
EIF2B4
EIF2B5
EIF2S1
EIF2S2
EIF2S3
EIF3A
EIF3B
EIF3C
EIF3D
EIF3E
EIF3F
EIF3G
EIF3H
EIF3I
EIF3J
EIF3L
EIF3M
EIF4A1
EIF4A3

EIF4B
EIF4E
EIF4G1
EIF4G2
EIF5
EIF5A
EIF5B
EIF6
ELAC2
ELL
ELOB
ELOC
ELP1
ELP2
ELP3
ELP4
ELP5
ELP6
EMC1
EMC3
EMC4
EMC7
ENY2
EP400
EPOP
EPRS
ERAL1
ERCC2
ERCC3
ERH
ERVW-1
ESF1
ESPL1
ESS2
ETF1
EWSR1
EXOC1
EXOC2
EXOC3
EXOC4
EXOC5
EXOC7
EXOC8
EXOSC1
EXOSC10

EXOSC2
EXOSC3
EXOSC4
EXOSC5
EXOSC6
EXOSC7
EXOSC8
EXOSC9
FAAP100
FAAP20
FAAP24
FAM133B
FAM136A
FAM166A
FAM229A
FAM32A
FAM69A
FAM86C1
FARS2
FARSA
FARSB
FAU
FBL
FBLIM1
FBXO5
FCF1
FDPS
FDX2
FDXR
FECH
FEN1
FIP1L1
FNBP4
FNTA
FNTB
FOXD4
FPGS
FTSJ3
FXN
GABPA
GABPB1
GAK
GAPDH
GAR1
GARS

GATAD2A
GBF1
GCN1
GEMIN2
GEMIN4
GEMIN5
GEMIN6
GEMIN7
GEMIN8
GFER
GFM1
GFPT1
GGPS1
GGT1
GGTLC1
GGTLC2
GINS1
GINS2
GINS3
GINS4
GJA3
GLE1
GLRX5
GLTP
GMPPB
GMPS
GNB1L
GNL2
GNL3
GNL3L
GOLGA6D
GON4L
GOSR2
GPKOW
GPN1
GPN2
GPN3
GPR61
GPS1
GRB2
GRPEL1
GRWD1
GSDMA
GSPT1
GTF2A1

GTF2A2
GTF2B
GTF2E1
GTF2E2
GTF2F1
GTF2F2
GTF2H1
GTF2H2
GTF2H2C
GTF2H3
GTF2H4
GTF3A
GTF3C1
GTF3C2
GTF3C3
GTF3C4
GTF3C5
GTPBP4
GUK1
H2AFJ
H2AFX
H2AFZ
H3F3A
H3F3B
HAMP
HAP1
HAPLN2
HARS
HARS2
HAUS1
HAUS2
HAUS3
HAUS4
HAUS5
HAUS6
HAUS7
HAUS8
HCFC1
HCFC1R1
HDAC3
HEATR1
HGS
HIGD2A
HINFP
HIRA

HIST1H1C
HIST1H2AB
HIST1H2AC
HIST1H2AD
HIST1H2AE
HIST1H2AG
HIST1H2AH
HIST1H2AK
HIST1H2AL
HIST1H2AM
HIST1H2BB
HIST1H2BC
HIST1H2BD
HIST1H2BE
HIST1H2BF
HIST1H2BG
HIST1H2BH
HIST1H2BJ
HIST1H2BK
HIST1H2BL
HIST1H2BM
HIST1H2BN
HIST1H2BO
HIST1H3A
HIST1H3C
HIST1H3G
HIST1H3I
HIST2H2AC
HIST2H2BE
HIST2H2BF
HIST2H3D
HIST3H3
HJURP
HJV
HMGA1
HMGB1
HMGCR
HMGCS1
HMGN2
HNRNPC
HNRNPH1
HNRNPK
HNRNPL
HNRNPM
HNRNPR

HNRNPU
HOXC10
HRK
HS6ST1
HSCB
HSD17B10
HSF1
HSP90B1
HSPA5
HSPA8
HSPA9
HSPD1
HSPE1
HTATSF1
HUWE1
HYOU1
HYPK
IARS
IARS2
ICE1
IFITM2
IFITM3
IFNA21
IFNL2
IGBP1
IK
ILF2
ILF3
IMMT
IMP3
IMP4
IMPDH2
INCENP
ING3
INHBE
INO80
INO80B
INS
INTS1
INTS10
INTS11
INTS12
INTS14
INTS2
INTS3

INTS4
INTS5
INTS6
INTS7
INTS8
INTS9
IPO11
IPO13
IPO7
IPO9
ISCA2
ISCU
ISG20L2
ISY1
ISYNA1
JAZF1
JMJD6
KANSL1
KANSL2
KANSL3
KARS
KAT5
KAT8
KATNB1
KCMF1
KCNA10
KCNJ12
KDM2A
KDM8
KIAA0100
KIAA0391
KIAA1143
KIF11
KIF14
KIF18A
KIF18B
KIF1BP
KIF20A
KIF22
KIF23
KIF4A
KIN
KLC2
KLF16
KLHL11

KMT5A
KNL1
KPNB1
KRR1
KRT8
KRT86
KRTAP10-6
KRTAP10-9
KRTAP1-3
KRTAP17-1
KRTAP19-3
KRTAP21-1
KRTAP4-11
KRTAP4-2
KRTAP4-6
KRTAP5-1
KRTAP5-10
KRTAP5-11
KRTAP5-3
KRTAP5-7
KRTAP9-6
KTI12
LAGE3
LAMTOR2
LAMTOR3
LAMTOR4
LARS
LARS2
LAS1L
LCE1A
LCE1E
LCE2A
LCE2B
LCE3A
LCE5A
LENG1
LENG8
LETM1
LGALS9B
LIAS
LILRA6
LIMD2
LIN52
LIN54
LONP1

LRPPRC
LRR1
LRRC37A3
LRRC37B
LSG1
LSM10
LSM11
LSM14A
LSM2
LSM3
LSM4
LSM5
LSM6
LSM7
LSM8
LST1
LTF
LTO1
LTV1
LUC7L3
LYRM4
LYSMD1
MAD2L1
MAD2L1BP
MAD2L2
MAGEA6
MAGOH
MAK16
MANF
MAP1LC3B
MARS
MARS2
MASTL
MAT2A
MAU2
MAX
MBTD1
MBTPS1
MBTPS2
MCL1
MCM10
MCM2
MCM3
MCM3AP
MCM4

MCM5
MCM6
MCM7
MCMBP
MCRS1
MDN1
MEAF6
MED1
MED10
MED11
MED12
MED14
MED17
MED18
MED20
MED21
MED22
MED26
MED27
MED28
MED29
MED30
MED31
MED4
MED6
MED7
MED8
MED9
MEIOC
MEPCE
METAP1
METAP2
METTL1
METTL14
METTL16
METTL17
METTL3
MFAP1
MFN2
MINOS1
MIS12
MIS18A
MIS18BP1
MKRN1
MLST8

MLX
MMS19
MMS22L
MNAT1
MOB4
MOCS3
MPHOSPH10
MRE11
MRGBP
MRPL10
MRPL12
MRPL14
MRPL15
MRPL16
MRPL17
MRPL18
MRPL19
MRPL2
MRPL20
MRPL21
MRPL23
MRPL24
MRPL27
MRPL28
MRPL3
MRPL32
MRPL33
MRPL34
MRPL35
MRPL36
MRPL37
MRPL38
MRPL39
MRPL4
MRPL41
MRPL43
MRPL44
MRPL45
MRPL46
MRPL47
MRPL48
MRPL49
MRPL50
MRPL51
MRPL53

MRPL54
MRPL55
MRPL57
MRPL58
MRPL9
MRPS10
MRPS11
MRPS12
MRPS14
MRPS16
MRPS18A
MRPS18B
MRPS2
MRPS21
MRPS22
MRPS23
MRPS24
MRPS25
MRPS26
MRPS27
MRPS28
MRPS31
MRPS34
MRPS35
MRPS5
MRPS6
MRPS7
MRPS9
MRTO4
MSANTD3
MSL1
MST1
MSTO1
MT1A
MT1B
MT1E
MT1H
MTBP
MTG2
MTIF2
MTOR
MTPAP
MTREX
MVD
MVK

MYBBP1A
MYC
MYO1H
MZT1
NAA10
NAA15
NAA20
NAA25
NAA30
NAA35
NAA50
NACA
NAE1
NAF1
NAPA
NAPG
NARS
NASP
NAT10
NBAS
NBPF3
NCAPD2
NCAPD3
NCAPG
NCAPG2
NCAPH
NCAPH2
NCBP1
NCBP2
NCKAP1
NCL
NCOA4
NDC1
NDC80
NDE1
NDOR1
NDUFA11
NDUFA13
NDUFA2
NDUFA4
NDUFA6
NDUFA8
NDUFAB1
NDUFAF1
NDUFAF3

NDUFAF4
NDUFAF8
NDUFB10
NDUFB3
NDUFB4
NDUFB6
NDUFB7
NDUFB8
NDUFB9
NDUFC2
NDUFS2
NDUFS5
NDUFS8
NDUFV1
NEDD1
NEDD8
NELFA
NELFB
NELFCD
NELFE
NEPRO
NFATC2IP
NFE2L3
NFKBIE
NFRKB
NFS1
NFYA
NFYB
NFYC
NGDN
NHLRC2
NHP2
NHP2L1
NIFK
NIP7
NISCH
NKAP
NKAPD1
NLE1
NMD3
NME2
NMT1
NOB1
NOC2L
NOC3L

NOC4L
NOL10
NOL11
NOL12
NOL6
NOL7
NOL8
NOL9
NOLC1
NOM1
NOP10
NOP14
NOP16
NOP2
NOP53
NOP56
NOP58
NOP9
NPAT
NPB
NPIPA5
NPLOC4
NPM1
NRBP1
NRDE2
NRF1
NSA2
NSF
NSL1
NSMCE1
NSMCE2
NSMCE3
NSUN4
NUBP1
NUBP2
NUDC
NUDCD3
NUDT21
NUF2
NUFIP1
NUMA1
NUP107
NUP133
NUP153
NUP155

NUP160
NUP205
NUP214
NUP35
NUP43
NUP50
NUP54
NUP58
NUP62
NUP85
NUP88
NUP93
NUP98
NUS1
NUTF2
NVL
NXF1
OGDH
OGT
OIP5
OLFML3
OPA1
ORC1
ORC3
ORC4
ORC5
ORC6
OSBP
OSGEP
OSTC
OTOP1
OXA1L
PA2G4
PABPC1
PABPN1
PAF1
PAFAH1B1
PAICS
PAK1IP1
PAM16
PAPOLA
PARS2
PAXBP1
PC
PCBP1

PCBP2
PCF11
PCID2
PCNA
PCYT1A
PDAP1
PDCD11
PDCD2
PDCD5
PDCD6
PDCD7
PDE4DIP
PDPK1
PDRG1
PEF1
PELO
PELP1
PES1
PFDN1
PFDN2
PFDN5
PFDN6
PFN1
PGAM1
PGAM4
PGD
PGGT1B
PGK1
PGS1
PHAX
PHB
PHB2
PHF12
PHF5A
PI4KA
PIFO
PIK3C3
PIK3R4
PIP5K1A
PISD
PKM
PKMYT1
PLA2G10
PLK1
PLK4

PLRG1
PMF1
PMPCA
PMPCB
PNISR
PNKP
PNN
PNO1
PNPT1
POGZ
POLA1
POLA2
POLD1
POLD2
POLD3
POLE
POLE2
POLG2
POLQ
POLR1A
POLR1B
POLR1C
POLR1E
POLR2B
POLR2C
POLR2D
POLR2E
POLR2F
POLR2G
POLR2H
POLR2I
POLR2K
POLR2L
POLR3A
POLR3B
POLR3C
POLR3D
POLR3E
POLR3F
POLR3H
POLR3K
POLRMT
POM121C
POMP
POP1

POP4
POP5
POP7
POTEG
POTEH
POTEI
POTEM
POU5F1B
PPA1
PPAN
PPARGC1B
PPAT
PPIA
PPIAL4G
PPIE
PPIL1
PPIL2
PPIL4
PPP1CA
PPP1CB
PPP1R10
PPP1R11
PPP1R12A
PPP1R14B
PPP1R15B
PPP1R2
PPP1R7
PPP1R8
PPP2CA
PPP2R1A
PPP2R3C
PPP4C
PPP4R2
PPP6C
PPRC1
PPWD1
PPY
PRAC1
PRAMEF18
PRC1
PRCC
PRDM10
PREB
PRELID1
PRELID3B

PRIM1
PRKAB2
PRKRA
PRKRIP1
PRMT1
PRMT5
PRPF18
PRPF19
PRPF3
PRPF31
PRPF38A
PRPF38B
PRPF39
PRPF4
PRPF40A
PRPF4B
PRPF6
PRPF8
PRR13
PRRC2A
PRUNE
PRUNE1
PSMA1
PSMA2
PSMA3
PSMA4
PSMA5
PSMA6
PSMA7
PSMB1
PSMB2
PSMB3
PSMB4
PSMB5
PSMB6
PSMB7
PSMC1
PSMC2
PSMC3
PSMC4
PSMC5
PSMC6
PSMD1
PSMD11
PSMD12

PSMD13
PSMD14
PSMD2
PSMD3
PSMD4
PSMD6
PSMD7
PSMD8
PSME1
PSMG2
PSMG3
PSMG4
PTBP1
PTCD1
PTCD3
PTK2
PTMA
PTPA
PTPMT1
PTPN11
PTPN23
PTTG1
PUF60
PWP1
PWP2
PYM1
PYROXD1
PYURF
QARS
QRSL1
RAB1B
RAB6A
RABGGTA
RABGGTB
RABL2A
RAC1
RACGAP1
RACK1
RAD1
RAD17
RAD21
RAD51
RAD51C
RAD51D
RAD9A

RAE1
RAMAC
RAN
RANBP2
RANGAP1
RARS
RARS2
RASA4
RBBP4
RBBP5
RBBP6
RBBP8
RBM10
RBM14
RBM17
RBM19
RBM22
RBM25
RBM28
RBM39
RBM42
RBM48
RBM8A
RBMX
RBMX2
RBMXL1
RBX1
RCC1
RCC1L
RCL1
RCOR1
RELA
REV3L
RFC1
RFC2
RFC3
RFC4
RFC5
RFT1
RGP1
RHBDF1
RHEB
RHOQ
RILPL2
RINT1

RIOK1
RIOK2
RMI1
RMI2
RNF113A
RNF123
RNF168
RNF20
RNF4
RNF40
RNF8
RNGTT
RNMT
RNPC3
RNPS1
ROMO1
RPA1
RPA2
RPA3
RPAIN
RPAP1
RPAP2
RPAP3
RPE
RPF1
RPF2
RPL10
RPL10A
RPL11
RPL12
RPL13
RPL13A
RPL14
RPL15
RPL17
RPL18
RPL18A
RPL19
RPL21
RPL22
RPL23
RPL23A
RPL24
RPL26
RPL27

RPL27A
RPL28
RPL3
RPL30
RPL31
RPL32
RPL34
RPL35
RPL35A
RPL36
RPL36A
RPL36AL
RPL37
RPL37A
RPL38
RPL4
RPL5
RPL6
RPL7
RPL7A
RPL7L1
RPL8
RPL9
RPLP0
RPLP1
RPLP2
RPN1
RPN2
RPP14
RPP21
RPP30
RPP38
RPP40
RPS10
RPS11
RPS12
RPS13
RPS14
RPS15
RPS15A
RPS16
RPS17
RPS18
RPS19
RPS19BP1

RPS2
RPS20
RPS21
RPS23
RPS24
RPS25
RPS26
RPS27A
RPS28
RPS29
RPS3
RPS3A
RPS4X
RPS5
RPS6
RPS7
RPS8
RPS9
RPSA
RPTOR
RPUSD4
RRAGA
RRM1
RRM2
RRN3
RRP1
RRP12
RRP15
RRP7A
RRP9
RRS1
RSAD1
RSL1D1
RSL24D1
RSPH9
RSRC2
RTCB
RTEL1
RTF1
RTF2
RTRAF
RTTN
RUVBL1
RUVBL2
SACM1L

SAE1
SAMM50
SAP130
SAP18
SAP30BP
SARNP
SARS
SARS2
SART1
SART3
SASS6
SBDS
SBNO1
SCAP
SCD
SCFD1
SCG5
SCNM1
SCYL1
SDAD1
SDE2
SDHAF2
SDHC
SDHD
SEC13
SEC16A
SEC61A1
SEC61B
SEC61G
SEC62
SEC63
SEH1L
SEM1
SENP6
SERBP1
SERINC4
SETD1A
SETD2
SETDB1
SF1
SF3A1
SF3A2
SF3A3
SF3B1
SF3B2

SF3B3
SF3B4
SF3B5
SF3B6
SFPQ
SFSWAP
SGF29
SGO1
SHFM1
SHQ1
SIN3A
SINHCAF
SKA1
SKA2
SKA3
SKP1
SKP2
SLBP
SLC25A10
SLC25A26
SLC25A3
SLC25A38
SLC25A5
SLC35B1
SLC39A10
SLC39A7
SLC3A2
SLC4A5
SLC7A6OS
SLC9B1
SLU7
SMAGP
SMARCA5
SMARCB1
SMARCE1
SMC1A
SMC2
SMC3
SMC4
SMC5
SMC6
SMG1
SMG5
SMG6
SMG7

SMNDC1
SMR3B
SMU1
SNAP23
SNAPC1
SNAPC2
SNAPC3
SNAPC4
SNAPC5
SNAPIN
SNF8
SNIP1
SNRNP200
SNRNP25
SNRNP27
SNRNP35
SNRNP40
SNRNP48
SNRNP70
SNRPA1
SNRPB
SNRPC
SNRPD1
SNRPD2
SNRPD3
SNRPE
SNRPF
SNRPG
SNU13
SNUPN
SNW1
SOD1
SOD2
SON
SOWAHC
SPAG5
SPATA31A6
SPATA5
SPATA5L1
SPC24
SPC25
SPCS2
SPCS3
SPDL1
SPDYE1

SPOUT1
SPRR2G
SPRTN
SPRYD4
SPTLC1
SRBD1
SRCAP
SRF
SRFBP1
SRP14
SRP19
SRP54
SRP68
SRP72
SRP9
SRPRA
SRPRB
SRRM1
SRRM2
SRRT
SRSF1
SRSF10
SRSF11
SRSF2
SRSF3
SRSF6
SRSF7
SS18L2
SSB
SSBP1
SSBP3
SSRP1
SSSCA1
SSU72
STARD7
STAT5A
STIL
STRAP
STRIP1
STX18
STX5
SUDS3
SUGP1
SUGT1
SUMO2

SUPT16H
SUPT4H1
SUPT5H
SUPT6H
SUPV3L1
SURF6
SYF2
SYMPK
SYS1
SYVN1
TACC3
TADA2A
TAF1
TAF10
TAF12
TAF13
TAF1A
TAF1B
TAF1C
TAF1D
TAF2
TAF5
TAF6
TAF7
TAF8
TAMM41
TANGO6
TARDBP
TARS
TARS2
TBC1D28
TBCA
TBCB
TBCC
TBCD
TBCE
TBL3
TBP
TCERG1
TCF7
TCOF1
TCP1
TDGF1
TEFM
TELO2

TEN1
TERF1
TEX10
TFAM
TFB2M
TFDP1
TFIP11
TFRC
TGS1
THAP1
THAP11
THG1L
THOC1
THOC2
THOC3
THOC5
THOC6
THOC7
THOP1
TICRR
TIGD1
TIMELESS
TIMM10
TIMM13
TIMM22
TIMM23
TIMM29
TIMM44
TIMM8A
TIMM9
TINF2
TIPIN
TKT
TLCD1
TLK2
TLN1
TMA16
TMA7
TMED10
TMED2
TMEM183A
TMEM199
TMEM258
TNFRSF8
TNNT2

TNPO3
TOE1
TOMM20
TOMM22
TOMM40
TONSL
TOP1
TOP2A
TOP3A
TOPBP1
TOR2A
TOX4
TP53RK
TPI1
TPR
TPRKB
TPT1
TPX2
TRA2B
TRAIP
TRAPPC1
TRAPPC11
TRAPPC3
TRAPPC4
TRAPPC5
TRAPPC8
TRIAP1
TRIM37
TRIM48
TRMT10C
TRMT112
TRMT5
TRMT6
TRNT1
TRPM7
TRRAP
TSEN2
TSEN34
TSEN54
TSFM
TSG101
TSPYL5
TSR1
TSR2
TSTA3

TTC1
TTC27
TTC4
TTF1
TTF2
TTI1
TTI2
TTK
TUBA1B
TUBA1C
TUBA3D
TUBB
TUBG1
TUBG2
TUBGCP2
TUBGCP3
TUBGCP4
TUBGCP5
TUBGCP6
TUFM
TUT1
TWISTNB
TWNK
TXN
TXNL4A
TXNL4B
U2AF1
U2AF2
U2SURP
UBA1
UBA2
UBA3
UBA5
UBA52
UBAP1
UBB
UBC
UBE2C
UBE2D3
UBE2H
UBE2I
UBE2L3
UBE2M
UBE2N
UBE2Z

UBL5
UBQLN4
UBR4
UBR5
UBTF
UFD1
UFM1
UGT1A6
UNC50
UPF1
UPF2
UQCC2
UQCC3
UQCR10
UQCR11
UQCRB
UQCRC1
UQCRC2
UQCRFS1
UQCRH
UQCRQ
URB1
URB2
URI1
URM1
UROD
USE1
USP10
USP17L5
USP18
USP36
USP37
USP39
USP5
USP7
USP8
USP9X
USPL1
UTP11
UTP14A
UTP15
UTP18
UTP20
UTP23
UTP25

UTP3
UTP4
UTP6
UXT
VARS
VARS2
VBP1
VCL
VCP
VDAC1
VEZT
VHL
VIRMA
VMP1
VPS13D
VPS16
VPS18
VPS25
VPS28
VPS33A
VPS35
VPS37A
VPS41
VPS51
VPS52
VPS54
VPS72
VTI1B
WAC
WARS
WARS2
WASHC2A
WBP1
WBP11
WDHD1
WDR1
WDR12
WDR18
WDR24
WDR25
WDR26
WDR3
WDR33
WDR36
WDR43

WDR46
WDR5
WDR55
WDR60
WDR61
WDR7
WDR70
WDR74
WDR75
WDR77
WDR82
WDR83
WDR92
WDTC1
WEE1
WNK1
WTAP
XAB2
XAGE3
XPO1
XPO5
XPOT
XRCC2
XRCC3
XRCC5
XRCC6
XRN1
XRN2
YAE1
YARS
YARS2
YBX1
YEATS2
YEATS4
YJEFN3
YJU2
YKT6
YPEL1
YPEL5
YRDC
YTHDC1
YWHAZ
YY1
ZBTB11
ZBTB17

ZBTB48
ZBTB8OS
ZC3H13
ZC3H18
ZC3H3
ZC3H8
ZCCHC9
ZCRB1
ZFC3H1
ZMAT2
ZMAT5
ZNF131
ZNF141
ZNF207
ZNF236
ZNF335
ZNF407
ZNF468
ZNF492
ZNF559
ZNF574
ZNF593
ZNF676
ZNF687
ZNF720
ZNF730
ZNF763
ZNF780B
ZNF830
ZNF85
ZNHIT1
ZNHIT2
ZNHIT3
ZNHIT6
ZNRD1
ZPR1
ZRANB2
ZRSR2
ZWINT
ZZZ3

sed on DepMap.
