## Supplementary table S4 for "A druggable addiction to *de novo* pyrimidine biosynthesis in diffuse midline glioma"

### Supplemental Table 3 (Related to Figure 1): DMG dependency genes ide

| | DMG cell line beta score | | | |
|---|---|---|---|---|
| Gene symbol | SF8628 beta score | DIPG1 beta score | DIPG13 beta score | Average beta score |
| UMPS | -1.9476 | -2.5365 | -1.6441 | -2.04273333 |
| CAD | -2.457 | -1.7409 | -1.8094 | -2.00243333 |
| PPIAL4F | -2.2128 | -1.9257 | -1.2401 | -1.79286667 |
| TCEB1 | -1.7287 | -1.7979 | -0.89827 | -1.47495667 |
| DUX4 | -1.8415 | -1.8364 | -0.64309 | -1.44033 |
| DIEXF | -1.4819 | -2.1914 | -0.51606 | -1.39645333 |
| PRIM2 | -2.0207 | -1.3635 | -0.69287 | -1.35902333 |
| ADAT2 | -1.4354 | -1.6151 | -0.50848 | -1.18632667 |
| TIMMDC1 | -1.3461 | -1.279 | -0.8151 | -1.14673333 |
| G6PD | -1.4908 | -0.88694 | -1.0176 | -1.13178 |
| USP17L24 | -1.7213 | -1.13 | -0.50133 | -1.11754333 |
| FANCC | -1.6804 | -0.90175 | -0.55627 | -1.04614 |
| OARD1 | -1.4143 | -0.89603 | -0.67676 | -0.99569667 |
| LSS | NA | -2.6953 | -2.2127 | -2.454 |
| KIAA2012 | -2.6991 | -1.8782 | NA | -2.28865 |
| CEP55 | -1.9976 | -2.3955 | NA | -2.19655 |
| VPRBP | -2.1979 | -2.1632 | NA | -2.18055 |
| ICE2 | -1.4054 | -2.4522 | NA | -1.9288 |
| UFD1L | -2.0068 | -1.6746 | NA | -1.8407 |
| CTC1 | -2.0812 | -1.4917 | NA | -1.78645 |
| ANKRD49 | -1.9812 | -1.5194 | NA | -1.7503 |
| CCDC101 | -2.3759 | -1.0799 | NA | -1.7279 |
| KIF3B | -1.6966 | -1.7507 | NA | -1.72365 |
| TOB1 | -1.9022 | -1.5419 | NA | -1.72205 |
| MBD6 | -1.91 | -1.5126 | NA | -1.7113 |
| NPEPPS | -2.3969 | -1.0241 | NA | -1.7105 |
| EGLN1 | -2.245 | -1.1708 | NA | -1.7079 |
| FAM60A | -1.8839 | -1.4216 | NA | -1.65275 |
| TIMM23B | -1.9279 | -1.2742 | NA | -1.60105 |
| ELOF1 | -2.0775 | -1.1222 | NA | -1.59985 |
| FASN | NA | -1.1834 | -2.014 | -1.5987 |
| CCDC94 | -1.9962 | -1.1337 | NA | -1.56495 |
| MSMO1 | NA | -2.0156 | -1.1079 | -1.56175 |
| GFM2 | -1.4055 | -1.709 | NA | -1.55725 |
| FBXW7 | -2.2496 | -0.80559 | NA | -1.527595 |
| PPCS | NA | -2.4174 | -0.6372 | -1.5273 |
| NSDHL | NA | -1.9786 | -1.0754 | -1.527 |
| MAEA | -1.3398 | -1.6962 | NA | -1.518 |
| FANCG | -1.5807 | -1.4517 | NA | -1.5162 |
| COL4A6 | -1.912 | -1.111 | NA | -1.5115 |

| | | | | |
|---|---|---|---|---|
| TMPRSS9 | -1.6412 | -1.3585 | NA | -1.49985 |
| MRPL13 | -1.4748 | -1.5209 | NA | -1.49785 |
| RPL39 | NA | -2.2462 | -0.7327 | -1.48945 |
| FGFR1 | NA | -1.0506 | -1.9125 | -1.48155 |
| TCP11 | -1.4288 | -1.5331 | NA | -1.48095 |
| FAM210A | -1.5953 | -1.3633 | NA | -1.4793 |
| RPS27 | NA | -1.7502 | -1.1629 | -1.45655 |
| HLF | -1.7289 | -1.1838 | NA | -1.45635 |
| PTGES3L | -1.9472 | -0.92405 | NA | -1.435625 |
| POTEB2 | -1.6241 | -1.2074 | NA | -1.41575 |
| CHD2 | NA | -2.2397 | -0.56208 | -1.40089 |
| FAM180B | -1.7527 | -1.0404 | NA | -1.39655 |
| ORAOV1 | NA | -0.9657 | -1.8236 | -1.39465 |
| SP2 | -1.8212 | -0.96614 | NA | -1.39367 |
| ZNF567 | -1.839 | -0.93327 | NA | -1.386135 |
| DSCC1 | -1.8418 | -0.9013 | NA | -1.37155 |
| MRPS15 | -1.8966 | -0.84083 | NA | -1.368715 |
| GIGYF2 | -2.0464 | NA | -0.68227 | -1.364335 |
| RPL39L | -1.497 | -1.1624 | NA | -1.3297 |
| POLR2A | NA | -1.1291 | -1.5263 | -1.3277 |
| TRAPPC13 | NA | -1.5887 | -1.0666 | -1.32765 |
| TTC14 | -1.6116 | -1.0429 | NA | -1.32725 |
| RHOA | -1.3278 | -1.3203 | NA | -1.32405 |
| KNTC1 | -1.3576 | -1.2755 | NA | -1.31655 |
| COX15 | NA | -1.9622 | -0.65803 | -1.310115 |
| PRKDC | -1.5497 | -1.048 | NA | -1.29885 |
| ANKRD20A4 | -1.4036 | -1.1879 | NA | -1.29575 |
| FOSL1 | -1.682 | -0.90923 | NA | -1.295615 |
| SPATA2L | -1.6716 | -0.91722 | NA | -1.29441 |
| CCNI | -1.6796 | -0.88631 | NA | -1.282955 |
| PPIAL4E | NA | -1.5355 | -1.0276 | -1.28155 |
| RICTOR | -1.6048 | NA | -0.9524 | -1.2786 |
| DNA2 | NA | -1.4109 | -1.1309 | -1.2709 |
| GULP1 | -1.4618 | -1.0765 | NA | -1.26915 |
| PRDM9 | -1.6347 | -0.89891 | NA | -1.266805 |
| ILK | -1.4506 | -1.0813 | NA | -1.26595 |
| MBNL1 | -1.5986 | NA | -0.89358 | -1.24609 |
| POLR2J | NA | -1.5839 | -0.89384 | -1.23887 |
| CALHM3 | -1.572 | -0.89826 | NA | -1.23513 |
| SSX2IP | -1.5814 | -0.87625 | NA | -1.228825 |
| MEPE | -1.6235 | -0.81519 | NA | -1.219345 |
| WDR4 | -1.6882 | -0.7456 | NA | -1.2169 |
| NDUFB5 | NA | -1.7829 | -0.65023 | -1.216565 |
| GADD45GIP1 | -1.5681 | -0.8511 | NA | -1.2096 |
| GNB2L1 | NA | -1.2461 | -1.1513 | -1.1987 |

| | | | | |
|---|---|---|---|---|
| SQLE | NA | -1.4785 | -0.9113 | -1.1949 |
| TYMS | NA | -0.93679 | -1.4097 | -1.173245 |
| MS4A2 | -1.4252 | -0.90839 | NA | -1.166795 |
| BRIP1 | NA | -0.84195 | -1.4852 | -1.163575 |
| TERF2 | NA | -1.2655 | -1.0568 | -1.16115 |
| FH | NA | -1.4786 | -0.84308 | -1.16084 |
| A2M | -1.3542 | -0.94543 | NA | -1.149815 |
| MED15 | -1.5419 | -0.74859 | NA | -1.145245 |
| FGD6 | -1.3409 | -0.94617 | NA | -1.143535 |
| ISCA1 | NA | -1.6393 | -0.644 | -1.14165 |
| TBC1D3C | NA | -1.3411 | -0.93704 | -1.13907 |
| PPA2 | NA | -1.1101 | -1.1607 | -1.1354 |
| SLC31A1 | -1.3535 | -0.9135 | NA | -1.1335 |
| SEPSECS | NA | -1.2612 | -0.99933 | -1.130265 |
| PTAR1 | -1.2988 | -0.95537 | NA | -1.127085 |
| ARHGEF11 | -1.4153 | -0.83437 | NA | -1.124835 |
| POT1 | NA | -1.7181 | -0.5179 | -1.118 |
| SEC22B | NA | -1.327 | -0.90309 | -1.115045 |
| UCHL5 | -1.3534 | -0.8546 | NA | -1.104 |
| UGT1A5 | -1.4566 | -0.74637 | NA | -1.101485 |
| BBIP1 | -1.3393 | -0.85992 | NA | -1.09961 |
| USP17L30 | NA | -1.456 | -0.72924 | -1.09262 |
| DGCR14 | NA | -1.4075 | -0.71943 | -1.063465 |
| CCDC160 | -1.3675 | -0.7523 | NA | -1.0599 |
| PGLS | NA | -1.1372 | -0.98091 | -1.059055 |
| SALL4 | -1.3631 | -0.74925 | NA | -1.056175 |
| MGEA5 | NA | -0.88378 | -1.2237 | -1.05374 |
| UBA6 | NA | -1.0631 | -1.025 | -1.04405 |
| FANCI | NA | -1.4112 | -0.67453 | -1.042865 |
| PPP2R4 | NA | -0.80513 | -1.2664 | -1.035765 |
| ATP5O | NA | -1.3688 | -0.69394 | -1.03137 |
| UHRF1 | NA | -0.87888 | -1.1743 | -1.02659 |
| USO1 | NA | -1.1181 | -0.93396 | -1.02603 |
| PAMR1 | -1.2971 | -0.74531 | NA | -1.021205 |
| NBPF7 | -1.3972 | NA | -0.638 | -1.0176 |
| TBC1D3K | NA | -0.98013 | -1.0395 | -1.009815 |
| DLST | NA | -1.1187 | -0.89687 | -1.007785 |
| UBE2T | NA | -0.77947 | -1.2287 | -1.004085 |
| SKIV2L2 | NA | -0.75859 | -1.2423 | -1.000445 |
| LIX1L | NA | -1.3556 | -0.6448 | -1.0002 |
| VPS36 | NA | -0.84555 | -1.1497 | -0.997625 |
| MT1G | NA | -1.1612 | -0.82761 | -0.994405 |
| PPIAL4C | NA | -0.98757 | -0.99955 | -0.99356 |
| AMD1 | NA | -0.96055 | -1.0171 | -0.988825 |
| RNF25 | NA | -1.2109 | -0.75726 | -0.98408 |

| | | | | |
|---|---|---|---|---|
| ALG1L2 | NA | -1.3242 | -0.64371 | -0.983955 |
| AASDHPPT | NA | -0.96324 | -0.99923 | -0.981235 |
| SUZ12 | NA | -0.90936 | -1.0441 | -0.97673 |
| ATP5B | NA | -1.3887 | -0.55997 | -0.974335 |
| FAM96B | NA | -1.1839 | -0.76434 | -0.97412 |
| MUS81 | NA | -1.0118 | -0.93182 | -0.97181 |
| ELAVL1 | NA | -1.2248 | -0.71518 | -0.96999 |
| GOLGA6L1 | NA | -1.2185 | -0.71729 | -0.967895 |
| CENPH | NA | -1.3339 | -0.59895 | -0.966425 |
| DPY30 | NA | -0.90899 | -0.997 | -0.952995 |
| MAD1L1 | NA | -1.3923 | -0.51209 | -0.952195 |
| CBWD3 | NA | -1.1683 | -0.73272 | -0.95051 |
| BRAT1 | NA | -1.3422 | -0.55511 | -0.948655 |
| FANCL | NA | -1.3315 | -0.56159 | -0.946545 |
| TP53BP2 | -1.2939 | NA | -0.59592 | -0.94491 |
| TMEM56 | -1.3367 | NA | -0.5154 | -0.92605 |
| USP17L3 | NA | -1.2937 | -0.55188 | -0.92279 |
| USP17L22 | NA | -1.0142 | -0.8283 | -0.92125 |
| SAMD4B | NA | -1.0286 | -0.80826 | -0.91843 |
| RPIA | NA | -1.0882 | -0.74849 | -0.918345 |
| DDI2 | NA | -1.1398 | -0.68917 | -0.914485 |
| PRRC2C | NA | -0.89472 | -0.93128 | -0.913 |
| MYBL2 | NA | -1.0859 | -0.73396 | -0.90993 |
| DHODH | NA | -1.2956 | -0.52266 | -0.90913 |
| UTP11L | NA | -0.98633 | -0.82882 | -0.907575 |
| VPS53 | NA | -1.0046 | -0.80702 | -0.90581 |
| SYNCRIP | NA | -1.2294 | -0.54768 | -0.88854 |
| USP17L25 | NA | -1.2487 | -0.51951 | -0.884105 |
| ZMAT1 | NA | -1.0339 | -0.72832 | -0.88111 |
| SEL1L | NA | -0.87153 | -0.88648 | -0.879005 |
| GOLGA6L6 | NA | -0.74302 | -1.0007 | -0.87186 |
| BLM | NA | -1.0936 | -0.64427 | -0.868935 |
| MRPS33 | NA | -0.84322 | -0.88756 | -0.86539 |
| OR11G2 | NA | -1.1996 | -0.53075 | -0.865175 |
| CIRH1A | NA | -1.0174 | -0.70955 | -0.863475 |
| SHOC2 | NA | -1.1243 | -0.59643 | -0.860365 |
| AMPD2 | NA | -1.074 | -0.64216 | -0.85808 |
| GABARAP | NA | -0.75859 | -0.95692 | -0.857755 |
| IWS1 | NA | -1.0793 | -0.60154 | -0.84042 |
| NARFL | NA | -1.0656 | -0.60529 | -0.835445 |
| PLD3 | NA | -0.92514 | -0.73394 | -0.82954 |
| KEAP1 | NA | -0.75355 | -0.90206 | -0.827805 |
| SDHB | NA | -1.0516 | -0.60139 | -0.826495 |
| OR10J3 | NA | -0.99494 | -0.63707 | -0.816005 |
| LCMT1 | NA | -0.74806 | -0.87365 | -0.810855 |

| | | | | |
|---|---|---|---|---|
| FTH1 | NA | -0.92992 | -0.68212 | -0.80602 |
| HNRNPA1L2 | NA | -1.0991 | -0.51239 | -0.805745 |
| TMX2 | NA | -1.0397 | -0.56966 | -0.80468 |
| OTUD5 | NA | -0.74314 | -0.86503 | -0.804085 |
| PHF14 | NA | -1.0005 | -0.59775 | -0.799125 |
| COX4I1 | NA | -0.79167 | -0.7957 | -0.793685 |
| ATAD5 | NA | -0.88696 | -0.67924 | -0.7831 |
| ZNF594 | NA | -1.0346 | -0.53135 | -0.782975 |
| CDK5RAP1 | NA | -1.005 | -0.53796 | -0.77148 |
| RIMKLA | NA | -0.80523 | -0.7265 | -0.765865 |
| PATZ1 | NA | -0.86069 | -0.66968 | -0.765185 |
| VKORC1L1 | NA | -0.89816 | -0.62599 | -0.762075 |
| PYDC2 | NA | -0.90125 | -0.62223 | -0.76174 |
| ITPA | NA | -0.88682 | -0.63326 | -0.76004 |
| DHX30 | NA | -0.76358 | -0.74846 | -0.75602 |
| NT5C1A | NA | -0.86704 | -0.63797 | -0.752505 |
| FANCA | NA | -0.8394 | -0.66116 | -0.75028 |
| OR4F17 | NA | -0.88821 | -0.60262 | -0.745415 |
| HIST2H3A | NA | -0.98778 | -0.50049 | -0.744135 |
| OR4F29 | NA | -0.96952 | -0.51867 | -0.744095 |
| TRMT1 | NA | -0.96792 | -0.51663 | -0.742275 |
| UGP2 | NA | -0.74898 | -0.72861 | -0.738795 |
| DOPEY2 | NA | -0.86567 | -0.60853 | -0.7371 |
| SP6 | NA | -0.91454 | -0.54014 | -0.72734 |
| HIST2H3C | NA | -0.78221 | -0.66827 | -0.72524 |
| FUBP1 | NA | -0.79015 | -0.65522 | -0.722685 |
| DPYS | NA | -0.90236 | -0.52434 | -0.71335 |
| ATP5H | NA | -0.77709 | -0.63519 | -0.70614 |
| FRS2 | NA | -0.90387 | -0.50341 | -0.70364 |
| RBM27 | NA | -0.87842 | -0.51587 | -0.697145 |
| ATP5F1 | NA | -0.7615 | -0.6217 | -0.6916 |
| AHCYL1 | NA | -0.85065 | -0.52912 | -0.689885 |
| SOGA1 | NA | -0.79536 | -0.57973 | -0.687545 |
| FRG2B | NA | -0.84919 | -0.52451 | -0.68685 |
| STXBP3 | NA | -0.83257 | -0.51888 | -0.675725 |
| CRELD2 | NA | -0.76056 | -0.58935 | -0.674955 |
| ZNF568 | NA | -0.79081 | -0.55562 | -0.673215 |
| CD248 | NA | -0.74277 | -0.59906 | -0.670915 |
| PBRM1 | -17.477 | NA | NA | -17.477 |
| ZNF652 | -2.6971 | NA | NA | -2.6971 |
| GLTSCR1L | -2.5527 | NA | NA | -2.5527 |
| SYNJ2 | -2.4695 | NA | NA | -2.4695 |
| KCTD6 | -2.4177 | NA | NA | -2.4177 |
| HEATR5A | -2.3889 | NA | NA | -2.3889 |
| HEY1 | NA | -2.3707 | NA | -2.3707 |

| | | | | |
|---|---|---|---|---|
| OXSM | NA | -2.3254 | NA | -2.3254 |
| NDNL2 | NA | -2.2217 | NA | -2.2217 |
| UROS | NA | -2.221 | NA | -2.221 |
| IFNGR1 | -2.2153 | NA | NA | -2.2153 |
| VSIG10L | -2.2096 | NA | NA | -2.2096 |
| SOCS3 | -2.2026 | NA | NA | -2.2026 |
| HBS1L | NA | -2.1917 | NA | -2.1917 |
| WDR73 | NA | -2.1822 | NA | -2.1822 |
| PCDHB1 | NA | -2.1797 | NA | -2.1797 |
| MRPL11 | NA | -2.1754 | NA | -2.1754 |
| MSTN | -2.1448 | NA | NA | -2.1448 |
| GKAP1 | -2.1326 | NA | NA | -2.1326 |
| DNAH3 | -2.1182 | NA | NA | -2.1182 |
| MED24 | -2.0989 | NA | NA | -2.0989 |
| GNA13 | NA | -2.0632 | NA | -2.0632 |
| PHOSPHO2 | -2.0611 | NA | NA | -2.0611 |
| CRISP3 | -2.0606 | NA | NA | -2.0606 |
| GTSE1 | -2.0515 | NA | NA | -2.0515 |
| CFAP36 | -2.0496 | NA | NA | -2.0496 |
| XRCC1 | -2.0481 | NA | NA | -2.0481 |
| ITGAV | -2.0431 | NA | NA | -2.0431 |
| SLC35A2 | NA | -2.0419 | NA | -2.0419 |
| FANCD2 | NA | -2.0107 | NA | -2.0107 |
| CHD1 | -2.0089 | NA | NA | -2.0089 |
| GCSH | NA | -2.0054 | NA | -2.0054 |
| ZSCAN26 | -2.0014 | NA | NA | -2.0014 |
| NMNAT1 | NA | NA | -1.9804 | -1.9804 |
| PMVK | NA | -1.9693 | NA | -1.9693 |
| OSCP1 | -1.9661 | NA | NA | -1.9661 |
| KCNK10 | -1.9599 | NA | NA | -1.9599 |
| SOX10 | NA | NA | -1.9525 | -1.9525 |
| PHKA2 | -1.9483 | NA | NA | -1.9483 |
| PANK3 | NA | -1.9443 | NA | -1.9443 |
| TOPAZ1 | NA | -1.9435 | NA | -1.9435 |
| ALG6 | -1.9392 | NA | NA | -1.9392 |
| ZFP2 | -1.9219 | NA | NA | -1.9219 |
| KDELR3 | -1.9194 | NA | NA | -1.9194 |
| KIF24 | -1.8937 | NA | NA | -1.8937 |
| SLC17A5 | -1.8937 | NA | NA | -1.8937 |
| SENP7 | NA | -1.8927 | NA | -1.8927 |
| PBX3 | -1.8898 | NA | NA | -1.8898 |
| GOLGB1 | -1.8879 | NA | NA | -1.8879 |
| BMI1 | NA | -1.8876 | NA | -1.8876 |
| OR7D2 | -1.887 | NA | NA | -1.887 |
| NDUFA9 | NA | -1.8763 | NA | -1.8763 |

| | | | | |
|---|---|---|---|---|
| CDK11B | NA | -1.8649 | NA | -1.8649 |
| ADSS | NA | -1.8644 | NA | -1.8644 |
| PLEKHG1 | -1.8567 | NA | NA | -1.8567 |
| IDE | -1.8489 | NA | NA | -1.8489 |
| SLC22A23 | -1.8454 | NA | NA | -1.8454 |
| RFPL4AL1 | -1.8422 | NA | NA | -1.8422 |
| OR4F3 | NA | -1.8383 | NA | -1.8383 |
| GPATCH8 | -1.8378 | NA | NA | -1.8378 |
| PCSK1 | -1.836 | NA | NA | -1.836 |
| RABAC1 | -1.8313 | NA | NA | -1.8313 |
| A1CF | -1.8203 | NA | NA | -1.8203 |
| ZNF235 | -1.8176 | NA | NA | -1.8176 |
| PLEKHM2 | -1.8163 | NA | NA | -1.8163 |
| ACACA | NA | NA | -1.8082 | -1.8082 |
| S100A9 | -1.8068 | NA | NA | -1.8068 |
| ANKRD20A2 | NA | -1.8003 | NA | -1.8003 |
| RRAGC | -1.7992 | NA | NA | -1.7992 |
| CRYBG3 | -1.7991 | NA | NA | -1.7991 |
| SPATA24 | -1.7981 | NA | NA | -1.7981 |
| OR52E2 | -1.7891 | NA | NA | -1.7891 |
| USP47 | -1.7881 | NA | NA | -1.7881 |
| IPPK | NA | -1.787 | NA | -1.787 |
| SLC35A5 | -1.7702 | NA | NA | -1.7702 |
| FAM104A | -1.7689 | NA | NA | -1.7689 |
| RNF6 | -1.7644 | NA | NA | -1.7644 |
| ADRM1 | NA | -1.7588 | NA | -1.7588 |
| B3GALTL | -1.7479 | NA | NA | -1.7479 |
| HEATR3 | NA | -1.7474 | NA | -1.7474 |
| ZNF513 | -1.7458 | NA | NA | -1.7458 |
| PHF20L1 | -1.7447 | NA | NA | -1.7447 |
| SMN1 | NA | -1.7426 | NA | -1.7426 |
| OR52E6 | -1.7419 | NA | NA | -1.7419 |
| NFE2L1 | -1.7412 | NA | NA | -1.7412 |
| CEP295 | -1.7245 | NA | NA | -1.7245 |
| SERINC1 | -1.7217 | NA | NA | -1.7217 |
| TEX15 | NA | -1.7205 | NA | -1.7205 |
| FKTN | NA | -1.7188 | NA | -1.7188 |
| HNRNPCL3 | NA | -1.7103 | NA | -1.7103 |
| POLR2J2 | NA | -1.7097 | NA | -1.7097 |
| GLTSCR2 | NA | -1.7058 | NA | -1.7058 |
| MRPS30 | NA | -1.6987 | NA | -1.6987 |
| HUS1 | NA | -1.6974 | NA | -1.6974 |
| TNC | -1.6968 | NA | NA | -1.6968 |
| OR51B6 | -1.6967 | NA | NA | -1.6967 |
| PGA5 | -1.6943 | NA | NA | -1.6943 |

| | | | | |
|---|---|---|---|---|
| CCS | NA | -1.6897 | NA | -1.6897 |
| ATP5D | NA | -1.6771 | NA | -1.6771 |
| HNRNPUL2 | -1.6757 | NA | NA | -1.6757 |
| MPL | NA | -1.6741 | NA | -1.6741 |
| CAAP1 | -1.6695 | NA | NA | -1.6695 |
| EPHA2 | -1.665 | NA | NA | -1.665 |
| MTERF3 | NA | -1.6619 | NA | -1.6619 |
| TRAF3IP2 | -1.6594 | NA | NA | -1.6594 |
| DYX1C1 | -1.6584 | NA | NA | -1.6584 |
| PROS1 | -1.658 | NA | NA | -1.658 |
| PAXIP1 | -1.6556 | NA | NA | -1.6556 |
| SPATA13 | NA | -1.654 | NA | -1.654 |
| ADAR | NA | -1.6526 | NA | -1.6526 |
| FOXA3 | NA | -1.6504 | NA | -1.6504 |
| TMC2 | -1.6392 | NA | NA | -1.6392 |
| PDE12 | NA | -1.6371 | NA | -1.6371 |
| MTO1 | NA | -1.634 | NA | -1.634 |
| PDSS1 | NA | -1.6202 | NA | -1.6202 |
| SDHA | NA | -1.6196 | NA | -1.6196 |
| SCO2 | NA | -1.6128 | NA | -1.6128 |
| DHX58 | -1.6123 | NA | NA | -1.6123 |
| FAM186A | -1.6098 | NA | NA | -1.6098 |
| ALDH1L2 | -1.6084 | NA | NA | -1.6084 |
| ALG8 | NA | -1.6039 | NA | -1.6039 |
| SH3GLB2 | -1.6038 | NA | NA | -1.6038 |
| AQP10 | -1.6008 | NA | NA | -1.6008 |
| HIST1H4K | NA | -1.6008 | NA | -1.6008 |
| ALDH2 | -1.6 | NA | NA | -1.6 |
| ZNF554 | -1.5983 | NA | NA | -1.5983 |
| MPV17L | -1.5978 | NA | NA | -1.5978 |
| MYOM3 | NA | -1.5932 | NA | -1.5932 |
| APC | -1.5904 | NA | NA | -1.5904 |
| BIRC6 | NA | -1.5899 | NA | -1.5899 |
| CREBBP | -1.5853 | NA | NA | -1.5853 |
| FADD | -1.5794 | NA | NA | -1.5794 |
| TMEM2 | -1.5792 | NA | NA | -1.5792 |
| TRUB2 | NA | -1.5743 | NA | -1.5743 |
| NAA16 | -1.573 | NA | NA | -1.573 |
| SWI5 | -1.5679 | NA | NA | -1.5679 |
| PDP2 | -1.5671 | NA | NA | -1.5671 |
| BHMT | -1.5611 | NA | NA | -1.5611 |
| ODF1 | NA | -1.5602 | NA | -1.5602 |
| ZSCAN2 | -1.5578 | NA | NA | -1.5578 |
| IFNA8 | NA | -1.5471 | NA | -1.5471 |
| STOML1 | -1.5468 | NA | NA | -1.5468 |

| | | | | |
|---|---|---|---|---|
| CENPF | -1.5456 | NA | NA | -1.5456 |
| PDSS2 | NA | -1.5442 | NA | -1.5442 |
| HTT | -1.5426 | NA | NA | -1.5426 |
| ASUN | NA | -1.5424 | NA | -1.5424 |
| WASH1 | NA | NA | -1.5383 | -1.5383 |
| FZR1 | NA | -1.5376 | NA | -1.5376 |
| SECISBP2 | -1.5363 | NA | NA | -1.5363 |
| ANKAR | -1.5347 | NA | NA | -1.5347 |
| TNFRSF11B | NA | -1.5336 | NA | -1.5336 |
| MIPEP | NA | -1.5322 | NA | -1.5322 |
| PPP3R1 | -1.532 | NA | NA | -1.532 |
| PRDX1 | NA | NA | -1.5282 | -1.5282 |
| CKAP2L | NA | -1.5275 | NA | -1.5275 |
| MRPL40 | NA | -1.5263 | NA | -1.5263 |
| NGEF | -1.5193 | NA | NA | -1.5193 |
| ANKRD18A | NA | -1.5186 | NA | -1.5186 |
| C1QTNF2 | -1.5181 | NA | NA | -1.5181 |
| RPUSD3 | NA | -1.5177 | NA | -1.5177 |
| DARS2 | NA | -1.5151 | NA | -1.5151 |
| MRAS | -1.5144 | NA | NA | -1.5144 |
| MROH9 | -1.5121 | NA | NA | -1.5121 |
| TRAF6 | -1.5115 | NA | NA | -1.5115 |
| ZNF143 | NA | -1.5109 | NA | -1.5109 |
| DRG2 | -1.5092 | NA | NA | -1.5092 |
| KRTAP4-9 | -1.5071 | NA | NA | -1.5071 |
| PIWIL1 | -1.5053 | NA | NA | -1.5053 |
| CCDC30 | -1.5045 | NA | NA | -1.5045 |
| OR2M4 | -1.5043 | NA | NA | -1.5043 |
| SLC38A10 | -1.5037 | NA | NA | -1.5037 |
| .OC10050746: | NA | NA | -1.5027 | -1.5027 |
| MPP2 | -1.5026 | NA | NA | -1.5026 |
| CUL9 | NA | -1.5022 | NA | -1.5022 |
| NDUFS7 | NA | -1.5012 | NA | -1.5012 |
| LTN1 | -1.4978 | NA | NA | -1.4978 |
| RAPGEF6 | NA | -1.4959 | NA | -1.4959 |
| PDCL | NA | -1.4958 | NA | -1.4958 |
| PAK2 | NA | -1.4939 | NA | -1.4939 |
| EIF3CL | NA | -1.4926 | NA | -1.4926 |
| MRRF | NA | -1.4918 | NA | -1.4918 |
| RBM33 | -1.4915 | NA | NA | -1.4915 |
| ATP8B3 | NA | -1.4849 | NA | -1.4849 |
| MFSD11 | -1.484 | NA | NA | -1.484 |
| GUCY1A2 | NA | -1.4837 | NA | -1.4837 |
| LHB | -1.4826 | NA | NA | -1.4826 |
| ALDH18A1 | -1.4813 | NA | NA | -1.4813 |

| | | | | |
|---|---|---|---|---|
| ATP2B1 | -1.48 | NA | NA | -1.48 |
| HTR1F | NA | -1.48 | NA | -1.48 |
| VPS4A | -1.4798 | NA | NA | -1.4798 |
| COQ6 | NA | -1.4771 | NA | -1.4771 |
| ZNF783 | -1.4718 | NA | NA | -1.4718 |
| METTL4 | -1.471 | NA | NA | -1.471 |
| SPATA20 | -1.4708 | NA | NA | -1.4708 |
| CCDC171 | NA | -1.4702 | NA | -1.4702 |
| MTG1 | NA | -1.465 | NA | -1.465 |
| CCDC151 | NA | -1.4646 | NA | -1.4646 |
| RTN4IP1 | NA | -1.4636 | NA | -1.4636 |
| FCHO1 | NA | -1.4634 | NA | -1.4634 |
| COX8A | NA | -1.4627 | NA | -1.4627 |
| MSH6 | -1.4583 | NA | NA | -1.4583 |
| PLEKHH2 | -1.458 | NA | NA | -1.458 |
| SAR1B | -1.4571 | NA | NA | -1.4571 |
| CCNB3 | -1.4569 | NA | NA | -1.4569 |
| CYP2C18 | -1.4555 | NA | NA | -1.4555 |
| FAM154A | -1.4543 | NA | NA | -1.4543 |
| MAP4 | NA | -1.4537 | NA | -1.4537 |
| HAL | -1.4531 | NA | NA | -1.4531 |
| USP16 | NA | -1.4518 | NA | -1.4518 |
| TTC23 | -1.4507 | NA | NA | -1.4507 |
| CCSER1 | NA | -1.4507 | NA | -1.4507 |
| PCMT1 | -1.4504 | NA | NA | -1.4504 |
| RSF1 | NA | -1.4477 | NA | -1.4477 |
| RAB1A | NA | NA | -1.4475 | -1.4475 |
| RPGRIP1 | -1.4474 | NA | NA | -1.4474 |
| TUBD1 | NA | -1.4472 | NA | -1.4472 |
| BMP3 | NA | -1.4471 | NA | -1.4471 |
| N4BP1 | -1.4469 | NA | NA | -1.4469 |
| TP53BP1 | -1.4449 | NA | NA | -1.4449 |
| JMY | -1.4441 | NA | NA | -1.4441 |
| IER2 | -1.4421 | NA | NA | -1.4421 |
| SCLY | -1.4396 | NA | NA | -1.4396 |
| CPSF3L | NA | -1.4379 | NA | -1.4379 |
| HMGXB4 | NA | -1.4363 | NA | -1.4363 |
| PACRG | NA | -1.4355 | NA | -1.4355 |
| TUBB4B | NA | -1.4355 | NA | -1.4355 |
| CLIC3 | -1.4349 | NA | NA | -1.4349 |
| REV1 | NA | -1.4343 | NA | -1.4343 |
| KRTAP10-8 | -1.4291 | NA | NA | -1.4291 |
| SERF1A | -1.4283 | NA | NA | -1.4283 |
| CD96 | -1.4269 | NA | NA | -1.4269 |
| NDUFA1 | NA | -1.4265 | NA | -1.4265 |

| | | | | |
|---|---|---|---|---|
| PARP11 | -1.424 | NA | NA | -1.424 |
| GNPTAB | -1.4231 | NA | NA | -1.4231 |
| BATF2 | -1.4228 | NA | NA | -1.4228 |
| USP17L27 | NA | -1.4199 | NA | -1.4199 |
| CAMSAP1 | NA | -1.4197 | NA | -1.4197 |
| IDH3A | NA | -1.4178 | NA | -1.4178 |
| MTUS2 | NA | -1.4164 | NA | -1.4164 |
| CEP290 | -1.4163 | NA | NA | -1.4163 |
| CNPY3 | NA | -1.4162 | NA | -1.4162 |
| NF103-CHMP | NA | NA | -1.4151 | -1.4151 |
| BOD1L1 | NA | -1.413 | NA | -1.413 |
| IFI44 | NA | -1.4107 | NA | -1.4107 |
| NLGN3 | -1.4094 | NA | NA | -1.4094 |
| HELB | -1.409 | NA | NA | -1.409 |
| KRTAP4-5 | -1.4089 | NA | NA | -1.4089 |
| CARM1 | -1.4047 | NA | NA | -1.4047 |
| ACER1 | -1.4043 | NA | NA | -1.4043 |
| C9orf114 | NA | NA | -1.4041 | -1.4041 |
| PPP4R1 | -1.4035 | NA | NA | -1.4035 |
| PTPN1 | -1.4034 | NA | NA | -1.4034 |
| MATR3 | NA | -1.4021 | NA | -1.4021 |
| TRMT11 | NA | -1.4019 | NA | -1.4019 |
| NOA1 | NA | -1.4012 | NA | -1.4012 |
| TSNAXIP1 | -1.4009 | NA | NA | -1.4009 |
| ERGIC3 | -1.4 | NA | NA | -1.4 |
| NDUFA10 | NA | -1.3999 | NA | -1.3999 |
| C1QTNF9 | NA | -1.3992 | NA | -1.3992 |
| NPIPA3 | NA | -1.3973 | NA | -1.3973 |
| MON2 | NA | -1.3969 | NA | -1.3969 |
| HIST1H3F | NA | -1.3938 | NA | -1.3938 |
| CYP4F22 | -1.3936 | NA | NA | -1.3936 |
| NCR2 | NA | -1.3936 | NA | -1.3936 |
| CCDC169 | -1.3926 | NA | NA | -1.3926 |
| STRN3 | NA | -1.3922 | NA | -1.3922 |
| DET1 | -1.3915 | NA | NA | -1.3915 |
| NDUFAF2 | NA | -1.3895 | NA | -1.3895 |
| FBXO39 | -1.3891 | NA | NA | -1.3891 |
| PLP1 | -1.3875 | NA | NA | -1.3875 |
| RFFL | -1.3861 | NA | NA | -1.3861 |
| KRTAP5-4 | NA | -1.3858 | NA | -1.3858 |
| FAM171A2 | -1.3831 | NA | NA | -1.3831 |
| KIAA2018 | -1.3815 | NA | NA | -1.3815 |
| AMY2A | NA | -1.3814 | NA | -1.3814 |
| DEPDC1 | NA | -1.3799 | NA | -1.3799 |
| OR51G1 | -1.3798 | NA | NA | -1.3798 |

| | | | | |
|---|---|---|---|---|
| RXFP2 | -1.3748 | NA | NA | -1.3748 |
| NDUFAF7 | NA | -1.373 | NA | -1.373 |
| CSK | -1.3697 | NA | NA | -1.3697 |
| FAM65C | -1.3688 | NA | NA | -1.3688 |
| SERPINE2 | -1.3674 | NA | NA | -1.3674 |
| KDSR | NA | -1.367 | NA | -1.367 |
| ARF3 | NA | -1.3658 | NA | -1.3658 |
| KRT25 | -1.3645 | NA | NA | -1.3645 |
| ARID4B | NA | -1.3641 | NA | -1.3641 |
| SRXN1 | NA | NA | -1.3639 | -1.3639 |
| USP17L4 | NA | -1.3625 | NA | -1.3625 |
| AP3M2 | NA | -1.3621 | NA | -1.3621 |
| REXO2 | NA | -1.3594 | NA | -1.3594 |
| FANCM | NA | -1.3578 | NA | -1.3578 |
| LRFN5 | -1.3563 | NA | NA | -1.3563 |
| GPIHBP1 | -1.3542 | NA | NA | -1.3542 |
| COMMD8 | -1.3518 | NA | NA | -1.3518 |
| MROH7 | -1.3501 | NA | NA | -1.3501 |
| DRAM2 | -1.3473 | NA | NA | -1.3473 |
| PPP1R15A | NA | -1.3469 | NA | -1.3469 |
| USP17L17 | NA | -1.3464 | NA | -1.3464 |
| ING2 | -1.3462 | NA | NA | -1.3462 |
| ABCA5 | -1.346 | NA | NA | -1.346 |
| TRIM64 | -1.3458 | NA | NA | -1.3458 |
| CEACAM21 | NA | -1.3422 | NA | -1.3422 |
| NUP188 | NA | -1.3421 | NA | -1.3421 |
| GOLGA2 | -1.3411 | NA | NA | -1.3411 |
| TSSK3 | -1.3409 | NA | NA | -1.3409 |
| CNIH2 | NA | -1.3391 | NA | -1.3391 |
| MRPL22 | NA | -1.3382 | NA | -1.3382 |
| HMBS | NA | -1.3369 | NA | -1.3369 |
| TMEM194B | NA | -1.3361 | NA | -1.3361 |
| NSRP1 | NA | -1.3353 | NA | -1.3353 |
| MAP3K2 | -1.3345 | NA | NA | -1.3345 |
| UIMC1 | -1.3345 | NA | NA | -1.3345 |
| FAM50A | NA | NA | -1.3342 | -1.3342 |
| NSMCE4A | NA | -1.3335 | NA | -1.3335 |
| TINAG | NA | -1.3321 | NA | -1.3321 |
| SPANXB1 | -1.3301 | NA | NA | -1.3301 |
| SPATA31A7 | NA | -1.3301 | NA | -1.3301 |
| GBP5 | -1.3299 | NA | NA | -1.3299 |
| OR4K17 | -1.3288 | NA | NA | -1.3288 |
| DTNA | NA | -1.3285 | NA | -1.3285 |
| SLC25A40 | -1.3243 | NA | NA | -1.3243 |
| CCDC176 | NA | -1.3243 | NA | -1.3243 |

| | | | | |
|---|---|---|---|---|
| PLSCR2 | -1.324 | NA | NA | -1.324 |
| TMEM244 | -1.3204 | NA | NA | -1.3204 |
| KIAA1377 | NA | -1.3198 | NA | -1.3198 |
| USP17L20 | NA | -1.3189 | NA | -1.3189 |
| FAM58A | NA | -1.3187 | NA | -1.3187 |
| YBEY | NA | -1.3174 | NA | -1.3174 |
| MAL | -1.3173 | NA | NA | -1.3173 |
| RB1CC1 | NA | -1.3166 | NA | -1.3166 |
| HID1 | -1.3139 | NA | NA | -1.3139 |
| SH3BGR | NA | -1.3135 | NA | -1.3135 |
| GRSF1 | NA | -1.3133 | NA | -1.3133 |
| TXNRD1 | NA | NA | -1.313 | -1.313 |
| GLI1 | -1.3106 | NA | NA | -1.3106 |
| HEG1 | NA | -1.3089 | NA | -1.3089 |
| ZFP36L1 | -1.3085 | NA | NA | -1.3085 |
| SEPW1 | -1.308 | NA | NA | -1.308 |
| IDI1 | NA | -1.3076 | NA | -1.3076 |
| OAS2 | NA | -1.3064 | NA | -1.3064 |
| L36A-HNRNP | NA | NA | -1.3062 | -1.3062 |
| MRPL1 | NA | -1.3048 | NA | -1.3048 |
| SPAG4 | -1.3044 | NA | NA | -1.3044 |
| BEND3 | NA | -1.3036 | NA | -1.3036 |
| TNFRSF19 | NA | -1.3035 | NA | -1.3035 |
| OR4F5 | NA | -1.3033 | NA | -1.3033 |
| ERCC5 | -1.3023 | NA | NA | -1.3023 |
| LGALS9 | NA | -1.3011 | NA | -1.3011 |
| MSL3 | -1.301 | NA | NA | -1.301 |
| IDNK | NA | -1.3006 | NA | -1.3006 |
| FBXL12 | NA | -1.3005 | NA | -1.3005 |
| TXN2 | NA | -1.2997 | NA | -1.2997 |
| MZT2B | -1.2979 | NA | NA | -1.2979 |
| POTEE | -1.2978 | NA | NA | -1.2978 |
| HEYL | -1.2948 | NA | NA | -1.2948 |
| DMRTC1 | NA | -1.2948 | NA | -1.2948 |
| ZBTB26 | -1.2941 | NA | NA | -1.2941 |
| FBXW11 | NA | NA | -1.2938 | -1.2938 |
| CRHR1 | NA | -1.2931 | NA | -1.2931 |
| TBC1D3H | NA | -1.2922 | NA | -1.2922 |
| FDFT1 | NA | NA | -1.2913 | -1.2913 |
| SLC22A24 | NA | -1.2902 | NA | -1.2902 |
| ZNF852 | NA | -1.2877 | NA | -1.2877 |
| NHEJ1 | NA | -1.2876 | NA | -1.2876 |
| LEMD2 | NA | -1.2834 | NA | -1.2834 |
| JUN | NA | -1.2833 | NA | -1.2833 |
| MMP3 | NA | -1.2825 | NA | -1.2825 |

| | | | | |
|---|---|---|---|---|
| NPHS2 | NA | -1.28 | NA | -1.28 |
| HIST1H1A | NA | -1.2774 | NA | -1.2774 |
| N6AMT1 | NA | -1.2766 | NA | -1.2766 |
| CHUK | NA | -1.2662 | NA | -1.2662 |
| MEF2B | NA | -1.2658 | NA | -1.2658 |
| CYP51A1 | NA | -1.2654 | NA | -1.2654 |
| FAM110D | NA | -1.2653 | NA | -1.2653 |
| MED19 | NA | -1.2647 | NA | -1.2647 |
| FRG2 | NA | -1.2607 | NA | -1.2607 |
| ZW10 | NA | -1.2602 | NA | -1.2602 |
| RAD50 | NA | -1.2579 | NA | -1.2579 |
| EFTUD1 | NA | -1.2573 | NA | -1.2573 |
| ARMC5 | NA | -1.2569 | NA | -1.2569 |
| PIKFYVE | NA | -1.2556 | NA | -1.2556 |
| FKBPL | NA | -1.2545 | NA | -1.2545 |
| USP17L28 | NA | -1.254 | NA | -1.254 |
| NMRAL1 | NA | -1.2528 | NA | -1.2528 |
| TNPO1 | NA | -1.2497 | NA | -1.2497 |
| SCAF1 | NA | -1.2473 | NA | -1.2473 |
| CHGA | NA | -1.2454 | NA | -1.2454 |
| FAM98B | NA | -1.2443 | NA | -1.2443 |
| SAFB2 | NA | -1.2443 | NA | -1.2443 |
| TMC1 | NA | -1.2432 | NA | -1.2432 |
| TRMT61A | NA | -1.2425 | NA | -1.2425 |
| TRMT12 | NA | -1.2421 | NA | -1.2421 |
| ERCC1 | NA | -1.2388 | NA | -1.2388 |
| CHKA | NA | NA | -1.238 | -1.238 |
| DPH3P1 | NA | -1.2377 | NA | -1.2377 |
| PAQR8 | NA | -1.2373 | NA | -1.2373 |
| FOXH1 | NA | -1.2363 | NA | -1.2363 |
| RAD51B | NA | -1.2337 | NA | -1.2337 |
| PTPLB | NA | -1.231 | NA | -1.231 |
| BTBD1 | NA | -1.2257 | NA | -1.2257 |
| MCAT | NA | -1.2241 | NA | -1.2241 |
| UNC13D | NA | -1.2232 | NA | -1.2232 |
| NBPF12 | NA | -1.2222 | NA | -1.2222 |
| CL2L2-PABPN | NA | NA | -1.2219 | -1.2219 |
| SCN2A | NA | -1.2203 | NA | -1.2203 |
| FANCE | NA | -1.2196 | NA | -1.2196 |
| KRT24 | NA | -1.2192 | NA | -1.2192 |
| OR10H4 | NA | -1.2181 | NA | -1.2181 |
| NPIPA1 | NA | -1.218 | NA | -1.218 |
| TMEM165 | NA | -1.2176 | NA | -1.2176 |
| BCL2L2 | NA | NA | -1.2169 | -1.2169 |
| CNOT10 | NA | -1.2143 | NA | -1.2143 |

| | | | | |
|---|---|---|---|---|
| CRYBB1 | NA | -1.2141 | NA | -1.2141 |
| CLPB | NA | -1.2139 | NA | -1.2139 |
| MRPL52 | NA | -1.2128 | NA | -1.2128 |
| CCDC172 | NA | -1.2124 | NA | -1.2124 |
| PSTK | NA | -1.2107 | NA | -1.2107 |
| ISY1-RAB43 | NA | NA | -1.2106 | -1.2106 |
| SPTLC2 | NA | -1.21 | NA | -1.21 |
| MTFMT | NA | -1.2078 | NA | -1.2078 |
| PSMF1 | NA | -1.207 | NA | -1.207 |
| MAN2C1 | NA | -1.2067 | NA | -1.2067 |
| PIANP | NA | -1.2053 | NA | -1.2053 |
| KLHL40 | NA | -1.2022 | NA | -1.2022 |
| DRG1 | NA | -1.2015 | NA | -1.2015 |
| CNOT4 | NA | -1.2012 | NA | -1.2012 |
| WDR59 | NA | -1.2006 | NA | -1.2006 |
| OR9G4 | NA | -1.1998 | NA | -1.1998 |
| HSD17B12 | NA | -1.1981 | NA | -1.1981 |
| FLT3 | NA | -1.1968 | NA | -1.1968 |
| SLC43A1 | NA | -1.1961 | NA | -1.1961 |
| DPH5 | NA | -1.1958 | NA | -1.1958 |
| NUBPL | NA | -1.1956 | NA | -1.1956 |
| COX10 | NA | -1.1902 | NA | -1.1902 |
| RAI14 | NA | -1.1873 | NA | -1.1873 |
| DNAJC2 | NA | -1.1867 | NA | -1.1867 |
| SLC7A1 | NA | -1.1854 | NA | -1.1854 |
| MRPL42 | NA | -1.1835 | NA | -1.1835 |
| SCO1 | NA | -1.1824 | NA | -1.1824 |
| RFK | NA | -1.1808 | NA | -1.1808 |
| CMTM3 | NA | -1.18 | NA | -1.18 |
| KIAA1328 | NA | -1.1794 | NA | -1.1794 |
| SMARCA2 | NA | -1.1789 | NA | -1.1789 |
| ZRANB1 | NA | -1.1786 | NA | -1.1786 |
| FKBP2 | NA | -1.1781 | NA | -1.1781 |
| 7-Sep | NA | -1.1779 | NA | -1.1779 |
| GOLT1B | NA | -1.1769 | NA | -1.1769 |
| GNB2 | NA | -1.1758 | NA | -1.1758 |
| LIG3 | NA | -1.1758 | NA | -1.1758 |
| ZWILCH | NA | -1.1755 | NA | -1.1755 |
| RNASEH2A | NA | -1.1754 | NA | -1.1754 |
| CFLAR | NA | -1.1719 | NA | -1.1719 |
| VASH1 | NA | -1.1711 | NA | -1.1711 |
| DOLPP1 | NA | -1.171 | NA | -1.171 |
| RGPD8 | NA | -1.1706 | NA | -1.1706 |
| STK11 | NA | -1.1698 | NA | -1.1698 |
| HSD17B2 | NA | -1.1669 | NA | -1.1669 |

| | | | | |
|---|---|---|---|---|
| TMEM62 | NA | -1.1663 | NA | -1.1663 |
| RNASEH2B | NA | -1.1632 | NA | -1.1632 |
| ABL2 | NA | -1.1629 | NA | -1.1629 |
| PTPRCAP | NA | -1.1603 | NA | -1.1603 |
| ZC3H15 | NA | -1.1563 | NA | -1.1563 |
| MTHFD1 | NA | -1.155 | NA | -1.155 |
| TMEM132B | NA | -1.1548 | NA | -1.1548 |
| MGAT2 | NA | -1.1543 | NA | -1.1543 |
| FAM83C | NA | -1.1524 | NA | -1.1524 |
| GOLGA4 | NA | -1.1513 | NA | -1.1513 |
| TSPYL2 | NA | -1.1493 | NA | -1.1493 |
| SULT6B1 | NA | -1.1473 | NA | -1.1473 |
| ZNF558 | NA | -1.1469 | NA | -1.1469 |
| HELLS | NA | -1.1464 | NA | -1.1464 |
| UBE4B | NA | -1.1462 | NA | -1.1462 |
| XCL1 | NA | -1.1453 | NA | -1.1453 |
| TTC16 | NA | -1.144 | NA | -1.144 |
| GATB | NA | -1.1439 | NA | -1.1439 |
| ACHE | NA | -1.1438 | NA | -1.1438 |
| PUS7 | NA | -1.1437 | NA | -1.1437 |
| KIF27 | NA | -1.1433 | NA | -1.1433 |
| CBX4 | NA | NA | -1.1432 | -1.1432 |
| SHD | NA | -1.1427 | NA | -1.1427 |
| ADAM10 | NA | -1.1421 | NA | -1.1421 |
| EMC6 | NA | -1.1408 | NA | -1.1408 |
| PHF1 | NA | -1.1408 | NA | -1.1408 |
| ASH2L | NA | -1.1397 | NA | -1.1397 |
| ACSM2A | NA | -1.1393 | NA | -1.1393 |
| TMEM33 | NA | -1.1363 | NA | -1.1363 |
| MTRF1L | NA | -1.1342 | NA | -1.1342 |
| PPL | NA | -1.1337 | NA | -1.1337 |
| PRSS48 | NA | -1.1307 | NA | -1.1307 |
| TCF23 | NA | -1.1307 | NA | -1.1307 |
| HSPE1-MOB4 | NA | NA | -1.1293 | -1.1293 |
| ATXN1 | NA | -1.1289 | NA | -1.1289 |
| TRAPPC9 | NA | -1.1284 | NA | -1.1284 |
| YAP1 | NA | -1.1282 | NA | -1.1282 |
| RASSF8 | NA | -1.1281 | NA | -1.1281 |
| OR8U8 | NA | -1.1267 | NA | -1.1267 |
| CLN6 | NA | -1.1258 | NA | -1.1258 |
| CHP1 | NA | NA | -1.1247 | -1.1247 |
| NBN | NA | -1.1246 | NA | -1.1246 |
| SRRD | NA | -1.124 | NA | -1.124 |
| FANCF | NA | -1.1234 | NA | -1.1234 |
| WBSCR16 | NA | -1.1232 | NA | -1.1232 |

| | | | | |
|---|---|---|---|---|
| SMR3A | NA | -1.1231 | NA | -1.1231 |
| PHF11 | NA | -1.1229 | NA | -1.1229 |
| NLRC5 | NA | -1.121 | NA | -1.121 |
| KATNAL1 | NA | -1.1209 | NA | -1.1209 |
| TSTD1 | NA | -1.1195 | NA | -1.1195 |
| KLF10 | NA | -1.1194 | NA | -1.1194 |
| VIPR1 | NA | -1.1187 | NA | -1.1187 |
| WBSCR22 | NA | -1.1184 | NA | -1.1184 |
| ZNF689 | NA | -1.1181 | NA | -1.1181 |
| GART | NA | -1.116 | NA | -1.116 |
| PGBD3 | NA | -1.1153 | NA | -1.1153 |
| SLC35G6 | NA | -1.1151 | NA | -1.1151 |
| TRHDE | NA | -1.1133 | NA | -1.1133 |
| GANAB | NA | -1.1114 | NA | -1.1114 |
| MAGEB4 | NA | -1.1109 | NA | -1.1109 |
| CEP85 | NA | -1.1108 | NA | -1.1108 |
| OR13C8 | NA | -1.1051 | NA | -1.1051 |
| NIN | NA | -1.1049 | NA | -1.1049 |
| PET117 | NA | -1.104 | NA | -1.104 |
| PCDHB15 | NA | -1.1024 | NA | -1.1024 |
| ENPP3 | NA | -1.102 | NA | -1.102 |
| BEND6 | NA | -1.101 | NA | -1.101 |
| SNTG1 | NA | -1.099 | NA | -1.099 |
| SULT1E1 | NA | -1.0987 | NA | -1.0987 |
| DGKZ | NA | -1.0982 | NA | -1.0982 |
| NXPH4 | NA | -1.0976 | NA | -1.0976 |
| LIPT2 | NA | -1.097 | NA | -1.097 |
| GSTZ1 | NA | -1.0967 | NA | -1.0967 |
| TMEM261 | NA | -1.0964 | NA | -1.0964 |
| TFB1M | NA | -1.0923 | NA | -1.0923 |
| HOOK2 | NA | -1.0917 | NA | -1.0917 |
| HK2 | NA | -1.0916 | NA | -1.0916 |
| STK17B | NA | -1.0906 | NA | -1.0906 |
| DNHD1 | NA | -1.0889 | NA | -1.0889 |
| SUCLG1 | NA | -1.0876 | NA | -1.0876 |
| SPIDR | NA | -1.0875 | NA | -1.0875 |
| PPP2R2B | NA | -1.087 | NA | -1.087 |
| CEP152 | NA | -1.0868 | NA | -1.0868 |
| FAM157B | NA | -1.0864 | NA | -1.0864 |
| PRKRIR | NA | -1.0845 | NA | -1.0845 |
| PLEKHG6 | NA | -1.0832 | NA | -1.0832 |
| ORM2 | NA | -1.0827 | NA | -1.0827 |
| LRRC20 | NA | -1.0825 | NA | -1.0825 |
| ZFX | NA | -1.0818 | NA | -1.0818 |
| FCN2 | NA | -1.081 | NA | -1.081 |

| | | | | |
|---|---|---|---|---|
| C1QTNF9B | NA | -1.0794 | NA | -1.0794 |
| COL23A1 | NA | -1.0785 | NA | -1.0785 |
| ELL3 | NA | -1.078 | NA | -1.078 |
| TEAD3 | NA | -1.0778 | NA | -1.0778 |
| CCNJL | NA | -1.0777 | NA | -1.0777 |
| UBAP2L | NA | -1.0775 | NA | -1.0775 |
| NFU1 | NA | -1.0774 | NA | -1.0774 |
| EPHB6 | NA | -1.0767 | NA | -1.0767 |
| PDLIM4 | NA | -1.076 | NA | -1.076 |
| CREB1 | NA | -1.0746 | NA | -1.0746 |
| PHF23 | NA | -1.0744 | NA | -1.0744 |
| SLC30A9 | NA | -1.0737 | NA | -1.0737 |
| CARS2 | NA | -1.0732 | NA | -1.0732 |
| ZFP28 | NA | -1.073 | NA | -1.073 |
| ABI3BP | NA | -1.0718 | NA | -1.0718 |
| OR2T1 | NA | -1.0718 | NA | -1.0718 |
| CMTM1 | NA | -1.0712 | NA | -1.0712 |
| SLC6A20 | NA | -1.0704 | NA | -1.0704 |
| HIST1H3H | NA | -1.0703 | NA | -1.0703 |
| PPIAL4D | NA | -1.0699 | NA | -1.0699 |
| TAZ | NA | -1.0698 | NA | -1.0698 |
| MALSU1 | NA | -1.0693 | NA | -1.0693 |
| FAM133A | NA | -1.0691 | NA | -1.0691 |
| LZTS3 | NA | -1.068 | NA | -1.068 |
| DPH6 | NA | -1.0676 | NA | -1.0676 |
| KDM4E | NA | -1.067 | NA | -1.067 |
| GOLGA6C | NA | NA | -1.067 | -1.067 |
| RRP36 | NA | -1.0664 | NA | -1.0664 |
| CAGE1 | NA | -1.0659 | NA | -1.0659 |
| KLF1 | NA | -1.0655 | NA | -1.0655 |
| TRIM49 | NA | -1.0655 | NA | -1.0655 |
| PKN2 | NA | -1.0647 | NA | -1.0647 |
| SLC22A6 | NA | -1.0643 | NA | -1.0643 |
| HIST3H2BB | NA | -1.063 | NA | -1.063 |
| FASTKD5 | NA | -1.0628 | NA | -1.0628 |
| RNF17 | NA | -1.0627 | NA | -1.0627 |
| RPL29 | NA | -1.0619 | NA | -1.0619 |
| HIST1H4B | NA | -1.0618 | NA | -1.0618 |
| KIF2C | NA | NA | -1.0611 | -1.0611 |
| ANKZF1 | NA | -1.0598 | NA | -1.0598 |
| BRPF3 | NA | -1.0589 | NA | -1.0589 |
| SEMG2 | NA | -1.0582 | NA | -1.0582 |
| GSTA3 | NA | -1.058 | NA | -1.058 |
| FAM111A | NA | -1.0576 | NA | -1.0576 |
| CHCHD6 | NA | -1.0574 | NA | -1.0574 |

| | | | | |
|---|---|---|---|---|
| PSRC1 | NA | -1.0571 | NA | -1.0571 |
| NXF5 | NA | -1.0565 | NA | -1.0565 |
| PARP4 | NA | -1.0565 | NA | -1.0565 |
| ERICH5 | NA | -1.0554 | NA | -1.0554 |
| IST1 | NA | -1.0552 | NA | -1.0552 |
| SV2A | NA | -1.0545 | NA | -1.0545 |
| MARK2 | NA | -1.0538 | NA | -1.0538 |
| ZNF534 | NA | -1.0534 | NA | -1.0534 |
| CNGB1 | NA | -1.0533 | NA | -1.0533 |
| RABIF | NA | -1.0508 | NA | -1.0508 |
| MCPH1 | NA | -1.0507 | NA | -1.0507 |
| WDR78 | NA | -1.0506 | NA | -1.0506 |
| MECR | NA | -1.0468 | NA | -1.0468 |
| NUTM2G | NA | NA | -1.0463 | -1.0463 |
| DNAJC12 | NA | -1.0449 | NA | -1.0449 |
| OR5W2 | NA | -1.0448 | NA | -1.0448 |
| RTN3 | NA | -1.0439 | NA | -1.0439 |
| MRPS36 | NA | -1.0431 | NA | -1.0431 |
| GZF1 | NA | -1.0425 | NA | -1.0425 |
| KRT84 | NA | -1.0424 | NA | -1.0424 |
| TATDN1 | NA | -1.0424 | NA | -1.0424 |
| OR51A2 | NA | -1.042 | NA | -1.042 |
| RHOT2 | NA | -1.0418 | NA | -1.0418 |
| BRD2 | NA | -1.0407 | NA | -1.0407 |
| CCBE1 | NA | -1.0398 | NA | -1.0398 |
| CDC25B | NA | -1.0395 | NA | -1.0395 |
| BLOC1S5 | NA | -1.0385 | NA | -1.0385 |
| AIP | NA | -1.038 | NA | -1.038 |
| RGL3 | NA | -1.038 | NA | -1.038 |
| TECR | NA | -1.0379 | NA | -1.0379 |
| ABHD2 | NA | NA | -1.0368 | -1.0368 |
| UGT1A3 | NA | -1.0357 | NA | -1.0357 |
| ZNF577 | NA | -1.0347 | NA | -1.0347 |
| NADK | NA | NA | -1.0347 | -1.0347 |
| RBPJ | NA | -1.0335 | NA | -1.0335 |
| OR1J4 | NA | -1.0332 | NA | -1.0332 |
| TARSL2 | NA | -1.0328 | NA | -1.0328 |
| SH3BGRL2 | NA | -1.0326 | NA | -1.0326 |
| ORO7-PAM1[illegible] | NA | NA | -1.0319 | -1.0319 |
| OR2A1 | NA | -1.0315 | NA | -1.0315 |
| CEBPG | NA | -1.0304 | NA | -1.0304 |
| CASQ1 | NA | -1.0299 | NA | -1.0299 |
| DDRGK1 | NA | -1.0299 | NA | -1.0299 |
| GPATCH11 | NA | -1.0299 | NA | -1.0299 |
| LIG4 | NA | -1.0292 | NA | -1.0292 |

| | | | | |
|---|---|---|---|---|
| TMEM243 | NA | -1.0292 | NA | -1.0292 |
| HRH4 | NA | -1.029 | NA | -1.029 |
| SLC35A1 | NA | -1.0287 | NA | -1.0287 |
| SMC1B | NA | -1.0286 | NA | -1.0286 |
| HIST2H2AA4 | NA | -1.0283 | NA | -1.0283 |
| DSCR3 | NA | -1.0281 | NA | -1.0281 |
| PRR11 | NA | -1.028 | NA | -1.028 |
| AMY1A | NA | -1.0271 | NA | -1.0271 |
| DCPS | NA | -1.0262 | NA | -1.0262 |
| KLRB1 | NA | -1.0259 | NA | -1.0259 |
| PSMB11 | NA | -1.0259 | NA | -1.0259 |
| VMA21 | NA | -1.0259 | NA | -1.0259 |
| OR1A1 | NA | -1.0258 | NA | -1.0258 |
| RHAG | NA | -1.0256 | NA | -1.0256 |
| PDYN | NA | -1.0255 | NA | -1.0255 |
| CNKSR2 | NA | -1.0249 | NA | -1.0249 |
| VPS29 | NA | -1.0246 | NA | -1.0246 |
| NME6 | NA | -1.0244 | NA | -1.0244 |
| RIC8A | NA | -1.024 | NA | -1.024 |
| TNNT1 | NA | -1.0231 | NA | -1.0231 |
| CHI3L2 | NA | -1.0222 | NA | -1.0222 |
| RR5-ARHGAP | NA | NA | -1.0217 | -1.0217 |
| SFI1 | NA | -1.0215 | NA | -1.0215 |
| DDX3Y | NA | -1.0213 | NA | -1.0213 |
| ARHGAP12 | NA | -1.0207 | NA | -1.0207 |
| POGK | NA | -1.0201 | NA | -1.0201 |
| MYBPHL | NA | -1.0199 | NA | -1.0199 |
| ATP8A2 | NA | -1.0196 | NA | -1.0196 |
| TUSC2 | NA | -1.0181 | NA | -1.0181 |
| SEC23IP | NA | NA | -1.0179 | -1.0179 |
| ENO1 | NA | -1.0176 | NA | -1.0176 |
| DPCR1 | NA | -1.0156 | NA | -1.0156 |
| HCCS | NA | -1.0156 | NA | -1.0156 |
| VIM | NA | -1.0156 | NA | -1.0156 |
| FGF4 | NA | -1.0155 | NA | -1.0155 |
| IQSEC1 | NA | -1.0154 | NA | -1.0154 |
| CCL1 | NA | -1.0145 | NA | -1.0145 |
| COLQ | NA | -1.0135 | NA | -1.0135 |
| PHF6 | NA | -1.0134 | NA | -1.0134 |
| TFAP2C | NA | -1.0129 | NA | -1.0129 |
| SLC44A2 | NA | -1.0116 | NA | -1.0116 |
| NAPB | NA | -1.0112 | NA | -1.0112 |
| CRLF3 | NA | -1.0109 | NA | -1.0109 |
| USP1 | NA | -1.0104 | NA | -1.0104 |
| WSB2 | NA | NA | -1.0096 | -1.0096 |

| | | | | |
|---|---|---|---|---|
| SURF4 | NA | NA | -1.0078 | -1.0078 |
| ZNF408 | NA | -1.0076 | NA | -1.0076 |
| MRPS18C | NA | -1.0074 | NA | -1.0074 |
| OSGIN2 | NA | -1.0074 | NA | -1.0074 |
| CES5A | NA | -1.0066 | NA | -1.0066 |
| PCDH12 | NA | -1.0061 | NA | -1.0061 |
| BCAR1 | NA | -1.0057 | NA | -1.0057 |
| GOLGA6L22 | NA | -1.0052 | NA | -1.0052 |
| CCM2 | NA | -1.0051 | NA | -1.0051 |
| GATC | NA | -1.0037 | NA | -1.0037 |
| DEFB4A | NA | -1.0035 | NA | -1.0035 |
| DCP2 | NA | -1.0033 | NA | -1.0033 |
| RALYL | NA | -1.0031 | NA | -1.0031 |
| RBM11 | NA | -1.0026 | NA | -1.0026 |
| PALB2 | NA | -1.0006 | NA | -1.0006 |
| TJP1 | NA | -1.0002 | NA | -1.0002 |
| BLOC1S6 | NA | -1.0001 | NA | -1.0001 |
| SRGAP2B | NA | -0.99985 | NA | -0.99985 |
| FAM184A | NA | -0.9993 | NA | -0.9993 |
| AP3D1 | NA | -0.9992 | NA | -0.9992 |
| UGT2A1 | NA | -0.99907 | NA | -0.99907 |
| PSMA8 | NA | -0.99601 | NA | -0.99601 |
| SLC44A3 | NA | -0.99561 | NA | -0.99561 |
| PCDHGB1 | NA | -0.99542 | NA | -0.99542 |
| FBXO42 | NA | -0.99432 | NA | -0.99432 |
| ANO5 | NA | -0.99429 | NA | -0.99429 |
| KIAA1524 | NA | -0.99422 | NA | -0.99422 |
| AOC1 | NA | -0.99229 | NA | -0.99229 |
| MED23 | NA | -0.99207 | NA | -0.99207 |
| SYCP1 | NA | -0.99187 | NA | -0.99187 |
| SLAMF9 | NA | -0.99124 | NA | -0.99124 |
| KRTAP5-5 | NA | -0.99101 | NA | -0.99101 |
| GSPT2 | NA | -0.9907 | NA | -0.9907 |
| MICAL2 | NA | -0.9902 | NA | -0.9902 |
| PARP9 | NA | -0.9897 | NA | -0.9897 |
| NDUFC1 | NA | -0.98898 | NA | -0.98898 |
| ZSCAN30 | NA | -0.98881 | NA | -0.98881 |
| SPN | NA | -0.98818 | NA | -0.98818 |
| XPO7 | NA | -0.98817 | NA | -0.98817 |
| POLG | NA | -0.98729 | NA | -0.98729 |
| UPK3B | NA | -0.98718 | NA | -0.98718 |
| DNAJC24 | NA | -0.98659 | NA | -0.98659 |
| PIK3CA | NA | -0.9863 | NA | -0.9863 |
| QTRTD1 | NA | NA | -0.98549 | -0.98549 |
| SLC25A1 | NA | -0.98486 | NA | -0.98486 |

| | | | | |
|---|---|---|---|---|
| CDC42EP1 | NA | -0.98427 | NA | -0.98427 |
| ASL | NA | -0.98424 | NA | -0.98424 |
| TYW1 | NA | -0.98403 | NA | -0.98403 |
| GUCA2A | NA | -0.98394 | NA | -0.98394 |
| BTN3A3 | NA | -0.98387 | NA | -0.98387 |
| CEP89 | NA | -0.98367 | NA | -0.98367 |
| PODN | NA | -0.9836 | NA | -0.9836 |
| MAPK9 | NA | -0.98275 | NA | -0.98275 |
| ZC3H4 | NA | -0.98267 | NA | -0.98267 |
| PHLDA1 | NA | -0.9815 | NA | -0.9815 |
| LARP1 | NA | -0.98078 | NA | -0.98078 |
| SLC39A5 | NA | -0.97999 | NA | -0.97999 |
| HIST1H2BI | NA | -0.97989 | NA | -0.97989 |
| TARBP2 | NA | -0.97906 | NA | -0.97906 |
| PRRT4 | NA | -0.9787 | NA | -0.9787 |
| PDIA5 | NA | -0.97868 | NA | -0.97868 |
| A4GNT | NA | -0.97867 | NA | -0.97867 |
| ALKBH5 | NA | -0.97847 | NA | -0.97847 |
| OR1F1 | NA | -0.97826 | NA | -0.97826 |
| ACTR3C | NA | -0.97753 | NA | -0.97753 |
| DISP1 | NA | -0.97749 | NA | -0.97749 |
| ANKRD54 | NA | -0.97654 | NA | -0.97654 |
| REXO1 | NA | -0.97447 | NA | -0.97447 |
| PARG | NA | -0.97394 | NA | -0.97394 |
| OR4F4 | NA | -0.97345 | NA | -0.97345 |
| ACPT | NA | -0.97319 | NA | -0.97319 |
| GPRASP1 | NA | -0.97297 | NA | -0.97297 |
| GHITM | NA | -0.97292 | NA | -0.97292 |
| DUS4L | NA | -0.97173 | NA | -0.97173 |
| SERPINB13 | NA | -0.97139 | NA | -0.97139 |
| TCEB2 | NA | NA | -0.97105 | -0.97105 |
| NPPA | NA | -0.97073 | NA | -0.97073 |
| C1D | NA | -0.96798 | NA | -0.96798 |
| SLC10A4 | NA | -0.9679 | NA | -0.9679 |
| SEPHS1 | NA | -0.96722 | NA | -0.96722 |
| HNRNPA1 | NA | NA | -0.9669 | -0.9669 |
| HIST1H4I | NA | -0.96689 | NA | -0.96689 |
| ARHGAP30 | NA | -0.96688 | NA | -0.96688 |
| CDKN2AIP | NA | -0.96663 | NA | -0.96663 |
| MYO1G | NA | -0.96629 | NA | -0.96629 |
| SPOP | NA | -0.96528 | NA | -0.96528 |
| CHEK2 | NA | -0.96477 | NA | -0.96477 |
| USP17L29 | NA | -0.96462 | NA | -0.96462 |
| MTX1 | NA | -0.96433 | NA | -0.96433 |
| HBG1 | NA | -0.96428 | NA | -0.96428 |

| | | | | |
|---|---|---|---|---|
| MLKL | NA | -0.96401 | NA | -0.96401 |
| MTIF3 | NA | -0.96372 | NA | -0.96372 |
| DCAF8L2 | NA | -0.96357 | NA | -0.96357 |
| TOMM34 | NA | -0.96307 | NA | -0.96307 |
| RNMTL1 | NA | -0.96296 | NA | -0.96296 |
| XPO6 | NA | -0.96277 | NA | -0.96277 |
| SETD8 | NA | -0.96263 | NA | -0.96263 |
| GOT1L1 | NA | -0.96249 | NA | -0.96249 |
| USP15 | NA | -0.96186 | NA | -0.96186 |
| HPR | NA | -0.9617 | NA | -0.9617 |
| ALOX12B | NA | -0.96128 | NA | -0.96128 |
| FABP6 | NA | -0.96115 | NA | -0.96115 |
| NDUFA3 | NA | -0.96092 | NA | -0.96092 |
| KIF4B | NA | -0.96055 | NA | -0.96055 |
| ZNF2 | NA | -0.95994 | NA | -0.95994 |
| LEO1 | NA | -0.95993 | NA | -0.95993 |
| NUDCD2 | NA | -0.95928 | NA | -0.95928 |
| PUM1 | NA | -0.95894 | NA | -0.95894 |
| COA4 | NA | -0.95802 | NA | -0.95802 |
| TAS2R43 | NA | -0.95764 | NA | -0.95764 |
| SNX27 | NA | -0.95722 | NA | -0.95722 |
| E2F1 | NA | -0.95647 | NA | -0.95647 |
| IFT20 | NA | -0.95643 | NA | -0.95643 |
| C8B | NA | -0.95639 | NA | -0.95639 |
| FDX1L | NA | NA | -0.95558 | -0.95558 |
| GTPBP6 | NA | -0.95547 | NA | -0.95547 |
| LRRC58 | NA | -0.95513 | NA | -0.95513 |
| OR2D3 | NA | -0.95491 | NA | -0.95491 |
| SNAP25 | NA | -0.95441 | NA | -0.95441 |
| TIPRL | NA | -0.95409 | NA | -0.95409 |
| NREP | NA | -0.95367 | NA | -0.95367 |
| LUZP4 | NA | -0.95292 | NA | -0.95292 |
| OR6K6 | NA | -0.9527 | NA | -0.9527 |
| RMND1 | NA | -0.95253 | NA | -0.95253 |
| LSP1 | NA | -0.95238 | NA | -0.95238 |
| ZNF37A | NA | -0.95227 | NA | -0.95227 |
| ARMS2 | NA | -0.95101 | NA | -0.95101 |
| SPINK5 | NA | -0.95076 | NA | -0.95076 |
| ZNF33B | NA | -0.95006 | NA | -0.95006 |
| SLC35B4 | NA | -0.94942 | NA | -0.94942 |
| SYT10 | NA | -0.94935 | NA | -0.94935 |
| HIC1 | NA | -0.94911 | NA | -0.94911 |
| MZF1 | NA | -0.94875 | NA | -0.94875 |
| MORC1 | NA | -0.94852 | NA | -0.94852 |
| FERMT3 | NA | -0.94818 | NA | -0.94818 |

| | | | | |
|---|---|---|---|---|
| MAP3K9 | NA | -0.94794 | NA | -0.94794 |
| CEP350 | NA | NA | -0.94783 | -0.94783 |
| RTFDC1 | NA | -0.94735 | NA | -0.94735 |
| GPR116 | NA | -0.94718 | NA | -0.94718 |
| OR5D18 | NA | -0.94688 | NA | -0.94688 |
| EXO5 | NA | -0.94616 | NA | -0.94616 |
| CCDC89 | NA | -0.94604 | NA | -0.94604 |
| GTF3C6 | NA | -0.94583 | NA | -0.94583 |
| RGPD3 | NA | -0.94582 | NA | -0.94582 |
| GPR89B | NA | NA | -0.94528 | -0.94528 |
| RASEF | NA | -0.94506 | NA | -0.94506 |
| ASCC3 | NA | -0.94415 | NA | -0.94415 |
| OR8G5 | NA | -0.94381 | NA | -0.94381 |
| MPDU1 | NA | -0.94331 | NA | -0.94331 |
| SPTAN1 | NA | -0.94257 | NA | -0.94257 |
| USP17L19 | NA | -0.94239 | NA | -0.94239 |
| FOCAD | NA | -0.94238 | NA | -0.94238 |
| BAIAP2 | NA | -0.94183 | NA | -0.94183 |
| HOXD11 | NA | -0.94176 | NA | -0.94176 |
| DPP10 | NA | -0.9416 | NA | -0.9416 |
| DPM1 | NA | -0.94159 | NA | -0.94159 |
| CDH8 | NA | -0.94122 | NA | -0.94122 |
| YWHAH | NA | -0.93934 | NA | -0.93934 |
| IRAK1 | NA | -0.93865 | NA | -0.93865 |
| MIOS | NA | -0.93832 | NA | -0.93832 |
| NPIPA2 | NA | NA | -0.93812 | -0.93812 |
| LUZP2 | NA | -0.93802 | NA | -0.93802 |
| SSPO | NA | -0.93738 | NA | -0.93738 |
| CENPJ | NA | -0.9373 | NA | -0.9373 |
| ITIH4 | NA | -0.93697 | NA | -0.93697 |
| NIPBL | NA | -0.93691 | NA | -0.93691 |
| ATAD3A | NA | -0.93651 | NA | -0.93651 |
| POTEB3 | NA | -0.93637 | NA | -0.93637 |
| PGC | NA | -0.93563 | NA | -0.93563 |
| FAM71D | NA | -0.93557 | NA | -0.93557 |
| APOBR | NA | -0.93549 | NA | -0.93549 |
| POTEF | NA | -0.93432 | NA | -0.93432 |
| HERC2 | NA | -0.93392 | NA | -0.93392 |
| PLEC | NA | -0.93371 | NA | -0.93371 |
| SI | NA | -0.93348 | NA | -0.93348 |
| TMEM168 | NA | -0.93347 | NA | -0.93347 |
| ZNF76 | NA | -0.93345 | NA | -0.93345 |
| OR2A5 | NA | -0.93341 | NA | -0.93341 |
| ABCC9 | NA | -0.93313 | NA | -0.93313 |
| ALDOC | NA | -0.93214 | NA | -0.93214 |

| | | | | |
|---|---|---|---|---|
| ATIC | NA | -0.93152 | NA | -0.93152 |
| ADTRP | NA | -0.93136 | NA | -0.93136 |
| PLCB3 | NA | -0.9307 | NA | -0.9307 |
| VN1R5 | NA | -0.9307 | NA | -0.9307 |
| ZNF121 | NA | -0.93067 | NA | -0.93067 |
| FRMPD1 | NA | -0.93059 | NA | -0.93059 |
| CHURC1-FNTE | NA | NA | -0.93038 | -0.93038 |
| PRB1 | NA | NA | -0.93016 | -0.93016 |
| NAP1L5 | NA | -0.92975 | NA | -0.92975 |
| ABHD16A | NA | -0.92965 | NA | -0.92965 |
| ZNF654 | NA | -0.92908 | NA | -0.92908 |
| HSD3B7 | NA | -0.92873 | NA | -0.92873 |
| SPDYE3 | NA | -0.92859 | NA | -0.92859 |
| TAF4 | NA | -0.92845 | NA | -0.92845 |
| FAM157A | NA | NA | -0.92669 | -0.92669 |
| BCAP29 | NA | -0.92496 | NA | -0.92496 |
| MREG | NA | -0.92496 | NA | -0.92496 |
| SH2D1B | NA | -0.92453 | NA | -0.92453 |
| ERCC8 | NA | -0.92451 | NA | -0.92451 |
| OR5A2 | NA | -0.92409 | NA | -0.92409 |
| AGTR2 | NA | -0.92403 | NA | -0.92403 |
| RBFOX2 | NA | NA | -0.92352 | -0.92352 |
| MAPK8IP1 | NA | -0.92331 | NA | -0.92331 |
| CTNND1 | NA | -0.9231 | NA | -0.9231 |
| SORT1 | NA | -0.923 | NA | -0.923 |
| FOXE1 | NA | -0.92293 | NA | -0.92293 |
| SAMD8 | NA | -0.92286 | NA | -0.92286 |
| SPRR2B | NA | NA | -0.92216 | -0.92216 |
| MESDC2 | NA | -0.92195 | NA | -0.92195 |
| P3H2 | NA | -0.92167 | NA | -0.92167 |
| ADD3 | NA | -0.92157 | NA | -0.92157 |
| PPFIA3 | NA | -0.9211 | NA | -0.9211 |
| VANGL1 | NA | -0.92037 | NA | -0.92037 |
| LDHAL6B | NA | -0.92035 | NA | -0.92035 |
| STT3B | NA | -0.91967 | NA | -0.91967 |
| MPRIP | NA | -0.91947 | NA | -0.91947 |
| TPRG1L | NA | -0.91931 | NA | -0.91931 |
| MAGEA8 | NA | -0.91908 | NA | -0.91908 |
| NUPL2 | NA | -0.91834 | NA | -0.91834 |
| SIAH1 | NA | -0.91794 | NA | -0.91794 |
| TYMSOS | NA | -0.91743 | NA | -0.91743 |
| HIST1H2AJ | NA | -0.91689 | NA | -0.91689 |
| LRP6 | NA | -0.91589 | NA | -0.91589 |
| PDE2A | NA | -0.91578 | NA | -0.91578 |
| SLC45A2 | NA | -0.91577 | NA | -0.91577 |

| | | | | |
|---|---|---|---|---|
| CNOT7 | NA | -0.91573 | NA | -0.91573 |
| TRIM40 | NA | -0.91544 | NA | -0.91544 |
| CLOCK | NA | -0.91518 | NA | -0.91518 |
| SLC6A13 | NA | -0.91507 | NA | -0.91507 |
| EDDM3A | NA | -0.91473 | NA | -0.91473 |
| KIF1A | NA | -0.91463 | NA | -0.91463 |
| UQCC1 | NA | -0.91354 | NA | -0.91354 |
| TRABD | NA | NA | -0.91342 | -0.91342 |
| ZBED2 | NA | -0.91337 | NA | -0.91337 |
| CDK2AP2 | NA | -0.91287 | NA | -0.91287 |
| TPPP3 | NA | -0.91274 | NA | -0.91274 |
| PRIMA1 | NA | -0.91216 | NA | -0.91216 |
| CCDC92 | NA | -0.91203 | NA | -0.91203 |
| PIH1D1 | NA | -0.9119 | NA | -0.9119 |
| TLCD2 | NA | -0.91184 | NA | -0.91184 |
| CUTA | NA | -0.91156 | NA | -0.91156 |
| CNTN1 | NA | -0.91076 | NA | -0.91076 |
| CCL13 | NA | -0.91067 | NA | -0.91067 |
| CDK11A | NA | -0.91064 | NA | -0.91064 |
| MRPL30 | NA | -0.91048 | NA | -0.91048 |
| DEFB126 | NA | -0.91038 | NA | -0.91038 |
| WIF1 | NA | -0.91019 | NA | -0.91019 |
| CCDC144A | NA | -0.90932 | NA | -0.90932 |
| MLLT6 | NA | -0.90884 | NA | -0.90884 |
| NACA2 | NA | -0.90839 | NA | -0.90839 |
| MCFD2 | NA | NA | -0.9079 | -0.9079 |
| GUF1 | NA | -0.90771 | NA | -0.90771 |
| SLC4A1 | NA | -0.90742 | NA | -0.90742 |
| USB1 | NA | -0.90716 | NA | -0.90716 |
| SPTBN4 | NA | -0.90695 | NA | -0.90695 |
| IGF2BP2 | NA | -0.90646 | NA | -0.90646 |
| GRAP | NA | -0.90571 | NA | -0.90571 |
| FAM86B1 | NA | -0.9057 | NA | -0.9057 |
| ART1 | NA | -0.90546 | NA | -0.90546 |
| PDE11A | NA | -0.90513 | NA | -0.90513 |
| HS3ST1 | NA | -0.90408 | NA | -0.90408 |
| HRSP12 | NA | -0.9027 | NA | -0.9027 |
| CD22 | NA | -0.90227 | NA | -0.90227 |
| MBP | NA | -0.90202 | NA | -0.90202 |
| WIZ | NA | -0.90181 | NA | -0.90181 |
| MTERF4 | NA | -0.90077 | NA | -0.90077 |
| MUC13 | NA | -0.90046 | NA | -0.90046 |
| RAB17 | NA | -0.90044 | NA | -0.90044 |
| HTR1D | NA | -0.90039 | NA | -0.90039 |
| KATNBL1 | NA | -0.90001 | NA | -0.90001 |

| | | | | |
|---|---|---|---|---|
| SLC27A1 | NA | -0.89976 | NA | -0.89976 |
| CIZ1 | NA | -0.89961 | NA | -0.89961 |
| RARRES1 | NA | -0.8996 | NA | -0.8996 |
| SEC24C | NA | -0.89953 | NA | -0.89953 |
| GATA3 | NA | -0.89936 | NA | -0.89936 |
| PFAS | NA | -0.89925 | NA | -0.89925 |
| GTPBP10 | NA | -0.89919 | NA | -0.89919 |
| CPO | NA | -0.89854 | NA | -0.89854 |
| SUPT20H | NA | -0.89834 | NA | -0.89834 |
| HLA-C | NA | -0.89799 | NA | -0.89799 |
| MCOLN3 | NA | -0.89799 | NA | -0.89799 |
| EGLN2 | NA | -0.89787 | NA | -0.89787 |
| TBC1D3L | NA | NA | -0.897 | -0.897 |
| NPIPB6 | NA | -0.89687 | NA | -0.89687 |
| ARID1A | NA | NA | -0.89676 | -0.89676 |
| COX16 | NA | NA | -0.89676 | -0.89676 |
| FBP1 | NA | -0.8963 | NA | -0.8963 |
| FLII | NA | -0.89514 | NA | -0.89514 |
| GBP6 | NA | -0.8951 | NA | -0.8951 |
| VAPA | NA | NA | -0.89461 | -0.89461 |
| ADAM28 | NA | -0.89436 | NA | -0.89436 |
| HGD | NA | -0.89391 | NA | -0.89391 |
| ZNF705E | NA | -0.89359 | NA | -0.89359 |
| CCDC168 | NA | -0.89353 | NA | -0.89353 |
| ADAM23 | NA | -0.89349 | NA | -0.89349 |
| YAE1D1 | NA | NA | -0.89271 | -0.89271 |
| CPT1A | NA | -0.89252 | NA | -0.89252 |
| RET | NA | -0.89238 | NA | -0.89238 |
| GAPVD1 | NA | -0.8922 | NA | -0.8922 |
| PRR18 | NA | -0.89214 | NA | -0.89214 |
| LMF2 | NA | -0.89173 | NA | -0.89173 |
| DND1 | NA | -0.89151 | NA | -0.89151 |
| KDM4A | NA | -0.89087 | NA | -0.89087 |
| CD81 | NA | -0.89046 | NA | -0.89046 |
| LCNL1 | NA | -0.89031 | NA | -0.89031 |
| GID8 | NA | -0.89027 | NA | -0.89027 |
| ANG | NA | -0.89024 | NA | -0.89024 |
| KLHL3 | NA | -0.89019 | NA | -0.89019 |
| ZNF491 | NA | -0.89017 | NA | -0.89017 |
| NBPF14 | NA | -0.89003 | NA | -0.89003 |
| SAP25 | NA | -0.89003 | NA | -0.89003 |
| MMP11 | NA | -0.88989 | NA | -0.88989 |
| LAMTOR5 | NA | -0.88981 | NA | -0.88981 |
| DUSP6 | NA | -0.88978 | NA | -0.88978 |
| CCDC51 | NA | -0.88927 | NA | -0.88927 |

| | | | | |
|---|---|---|---|---|
| OR5D16 | NA | -0.88906 | NA | -0.88906 |
| KCNJ4 | NA | -0.88871 | NA | -0.88871 |
| ST13 | NA | -0.88832 | NA | -0.88832 |
| CBWD1 | NA | -0.88798 | NA | -0.88798 |
| ZFAND5 | NA | -0.88716 | NA | -0.88716 |
| CRYGN | NA | -0.88703 | NA | -0.88703 |
| LBHD1 | NA | -0.88702 | NA | -0.88702 |
| SNRPA | NA | -0.88665 | NA | -0.88665 |
| SRPR | NA | NA | -0.88638 | -0.88638 |
| GSTA1 | NA | -0.88615 | NA | -0.88615 |
| MEMO1 | NA | -0.88602 | NA | -0.88602 |
| TRIM43B | NA | -0.88541 | NA | -0.88541 |
| GMEB2 | NA | NA | -0.88541 | -0.88541 |
| FXYD3 | NA | -0.88505 | NA | -0.88505 |
| DNAH7 | NA | -0.88495 | NA | -0.88495 |
| FOXM1 | NA | NA | -0.88482 | -0.88482 |
| IL31 | NA | -0.88439 | NA | -0.88439 |
| PLCZ1 | NA | -0.88421 | NA | -0.88421 |
| TROAP | NA | -0.88394 | NA | -0.88394 |
| LCE1C | NA | -0.88355 | NA | -0.88355 |
| ELTD1 | NA | -0.88285 | NA | -0.88285 |
| CEP19 | NA | -0.88274 | NA | -0.88274 |
| WFDC3 | NA | -0.8826 | NA | -0.8826 |
| EDN3 | NA | -0.88212 | NA | -0.88212 |
| FOXD4L6 | NA | -0.88199 | NA | -0.88199 |
| MCF2L2 | NA | -0.88199 | NA | -0.88199 |
| BTN2A2 | NA | -0.88076 | NA | -0.88076 |
| DNAJC18 | NA | -0.87974 | NA | -0.87974 |
| S100A10 | NA | -0.87951 | NA | -0.87951 |
| TRIM56 | NA | -0.87925 | NA | -0.87925 |
| METTL24 | NA | -0.87919 | NA | -0.87919 |
| AKAP8L | NA | -0.87911 | NA | -0.87911 |
| CELF2 | NA | -0.87883 | NA | -0.87883 |
| ZNF582 | NA | -0.87854 | NA | -0.87854 |
| ATP5C1 | NA | -0.87839 | NA | -0.87839 |
| SEZ6L2 | NA | -0.87826 | NA | -0.87826 |
| PIK3C2A | NA | -0.87752 | NA | -0.87752 |
| GDF6 | NA | -0.8774 | NA | -0.8774 |
| TMC4 | NA | -0.87702 | NA | -0.87702 |
| TRIM74 | NA | -0.87677 | NA | -0.87677 |
| FIS1 | NA | NA | -0.87649 | -0.87649 |
| ELAC1 | NA | -0.87644 | NA | -0.87644 |
| AZIN1 | NA | -0.87643 | NA | -0.87643 |
| CCDC185 | NA | -0.87616 | NA | -0.87616 |
| PCDHA2 | NA | -0.87553 | NA | -0.87553 |

| | | | | |
|---|---|---|---|---|
| HSD3B2 | NA | -0.87433 | NA | -0.87433 |
| CYP26A1 | NA | -0.87426 | NA | -0.87426 |
| KREMEN1 | NA | -0.87417 | NA | -0.87417 |
| CD4 | NA | NA | -0.87388 | -0.87388 |
| ASTN2 | NA | -0.87382 | NA | -0.87382 |
| NOBOX | NA | -0.87381 | NA | -0.87381 |
| PFN3 | NA | -0.87336 | NA | -0.87336 |
| RGAG1 | NA | -0.87327 | NA | -0.87327 |
| UBR2 | NA | -0.87302 | NA | -0.87302 |
| C2orf27B | NA | NA | -0.87302 | -0.87302 |
| BRAP | NA | -0.87248 | NA | -0.87248 |
| ANKRD52 | NA | -0.87229 | NA | -0.87229 |
| KRT9 | NA | NA | -0.87202 | -0.87202 |
| FBXL7 | NA | -0.87181 | NA | -0.87181 |
| TIPARP | NA | NA | -0.87108 | -0.87108 |
| DEDD | NA | -0.87084 | NA | -0.87084 |
| SSBP4 | NA | -0.87003 | NA | -0.87003 |
| MRPS17 | NA | -0.86943 | NA | -0.86943 |
| CXCL17 | NA | -0.8693 | NA | -0.8693 |
| TBCK | NA | -0.86914 | NA | -0.86914 |
| IL18BP | NA | -0.86873 | NA | -0.86873 |
| HCRTR1 | NA | -0.86816 | NA | -0.86816 |
| MTHFD1L | NA | -0.86793 | NA | -0.86793 |
| KIAA1549L | NA | -0.86782 | NA | -0.86782 |
| RNF32 | NA | -0.86766 | NA | -0.86766 |
| GLTSCR1 | NA | NA | -0.86736 | -0.86736 |
| WDR72 | NA | -0.86731 | NA | -0.86731 |
| DNAAF5 | NA | -0.86728 | NA | -0.86728 |
| TMEM74 | NA | -0.86709 | NA | -0.86709 |
| ZNF267 | NA | -0.86691 | NA | -0.86691 |
| ARFGEF2 | NA | -0.86658 | NA | -0.86658 |
| IFNA13 | NA | -0.86642 | NA | -0.86642 |
| MAP3K12 | NA | -0.86618 | NA | -0.86618 |
| SLC35G3 | NA | -0.86572 | NA | -0.86572 |
| TLE4 | NA | -0.86559 | NA | -0.86559 |
| SNRPN | NA | -0.86555 | NA | -0.86555 |
| NKTR | NA | -0.86517 | NA | -0.86517 |
| ACSM1 | NA | -0.86496 | NA | -0.86496 |
| SLC10A2 | NA | -0.86483 | NA | -0.86483 |
| CT45A10 | NA | -0.86455 | NA | -0.86455 |
| CYB561A3 | NA | -0.86448 | NA | -0.86448 |
| BTF3L4 | NA | NA | -0.86441 | -0.86441 |
| SHB | NA | -0.86423 | NA | -0.86423 |
| OR51V1 | NA | -0.86414 | NA | -0.86414 |
| IRF1 | NA | -0.86391 | NA | -0.86391 |

| | | | | |
|---|---|---|---|---|
| TOMM70A | NA | NA | -0.86371 | -0.86371 |
| MAGEA4 | NA | -0.86312 | NA | -0.86312 |
| GPRASP2 | NA | -0.86309 | NA | -0.86309 |
| NKX6-3 | NA | -0.86305 | NA | -0.86305 |
| CHADL | NA | -0.86278 | NA | -0.86278 |
| NT5DC2 | NA | -0.86268 | NA | -0.86268 |
| ZNF396 | NA | -0.86195 | NA | -0.86195 |
| MPPED1 | NA | -0.8616 | NA | -0.8616 |
| GIMAP1 | NA | -0.86151 | NA | -0.86151 |
| CDCA3 | NA | -0.86139 | NA | -0.86139 |
| CCNJ | NA | -0.86131 | NA | -0.86131 |
| CCZ1B | NA | -0.86089 | NA | -0.86089 |
| RBM15 | NA | -0.86039 | NA | -0.86039 |
| KIAA1467 | NA | -0.85993 | NA | -0.85993 |
| IVL | NA | -0.85925 | NA | -0.85925 |
| DLAT | NA | -0.85902 | NA | -0.85902 |
| DPRX | NA | -0.85895 | NA | -0.85895 |
| KIAA0895 | NA | -0.85893 | NA | -0.85893 |
| CD1D | NA | -0.85874 | NA | -0.85874 |
| TRAM2 | NA | NA | -0.85869 | -0.85869 |
| EMD | NA | -0.85857 | NA | -0.85857 |
| TXNDC16 | NA | -0.8579 | NA | -0.8579 |
| CHD8 | NA | NA | -0.85773 | -0.85773 |
| TMEM167B | NA | -0.85757 | NA | -0.85757 |
| UGT2B4 | NA | -0.85726 | NA | -0.85726 |
| MAGEA2B | NA | -0.85723 | NA | -0.85723 |
| SMIM17 | NA | -0.8571 | NA | -0.8571 |
| CDADC1 | NA | NA | -0.85693 | -0.85693 |
| TRIM49D2P | NA | -0.85685 | NA | -0.85685 |
| ZC3HC1 | NA | -0.8568 | NA | -0.8568 |
| CLDN19 | NA | -0.85657 | NA | -0.85657 |
| HOPX | NA | -0.8561 | NA | -0.8561 |
| SLC5A6 | NA | -0.85587 | NA | -0.85587 |
| TEKT4 | NA | -0.85576 | NA | -0.85576 |
| ROCK1 | NA | -0.85563 | NA | -0.85563 |
| MAP3K7CL | NA | -0.85559 | NA | -0.85559 |
| 3-Sep | NA | -0.85537 | NA | -0.85537 |
| STKLD1 | NA | -0.85535 | NA | -0.85535 |
| FAM175B | NA | -0.85513 | NA | -0.85513 |
| CT47A3 | NA | -0.85475 | NA | -0.85475 |
| OR2C3 | NA | -0.8543 | NA | -0.8543 |
| KHDRBS1 | NA | -0.85427 | NA | -0.85427 |
| RAB44 | NA | -0.85352 | NA | -0.85352 |
| HN1L | NA | -0.85286 | NA | -0.85286 |
| CMTM2 | NA | -0.8528 | NA | -0.8528 |

| | | | | |
|---|---|---|---|---|
| HNRNPD | NA | -0.85269 | NA | -0.85269 |
| PLEKHG7 | NA | -0.85245 | NA | -0.85245 |
| ITM2B | NA | -0.85239 | NA | -0.85239 |
| FOXB1 | NA | -0.8519 | NA | -0.8519 |
| TTC23L | NA | -0.85154 | NA | -0.85154 |
| PIK3IP1 | NA | -0.85147 | NA | -0.85147 |
| ARL1 | NA | NA | -0.85143 | -0.85143 |
| SLCO1A2 | NA | -0.85094 | NA | -0.85094 |
| KRTAP5-6 | NA | -0.85076 | NA | -0.85076 |
| CWC15 | NA | -0.8507 | NA | -0.8507 |
| CCIN | NA | -0.85069 | NA | -0.85069 |
| SLC6A5 | NA | -0.85068 | NA | -0.85068 |
| OR10AD1 | NA | -0.84949 | NA | -0.84949 |
| SH3RF2 | NA | -0.84929 | NA | -0.84929 |
| RNF11 | NA | -0.84892 | NA | -0.84892 |
| HEMK1 | NA | -0.84883 | NA | -0.84883 |
| MBOAT7 | NA | -0.84874 | NA | -0.84874 |
| PITPNB | NA | -0.84832 | NA | -0.84832 |
| LRRN4CL | NA | -0.84821 | NA | -0.84821 |
| MAST2 | NA | -0.84807 | NA | -0.84807 |
| TADA2B | NA | -0.848 | NA | -0.848 |
| SUN2 | NA | -0.84777 | NA | -0.84777 |
| ANO7 | NA | -0.84761 | NA | -0.84761 |
| KDM4C | NA | -0.84761 | NA | -0.84761 |
| IKBKAP | NA | -0.8475 | NA | -0.8475 |
| CARD14 | NA | -0.84712 | NA | -0.84712 |
| DYDC2 | NA | -0.84629 | NA | -0.84629 |
| TALDO1 | NA | NA | -0.84574 | -0.84574 |
| STT3A | NA | -0.84571 | NA | -0.84571 |
| CDS2 | NA | NA | -0.84513 | -0.84513 |
| ZCCHC7 | NA | -0.84396 | NA | -0.84396 |
| UFL1 | NA | -0.84387 | NA | -0.84387 |
| TTC24 | NA | -0.84339 | NA | -0.84339 |
| OMA1 | NA | -0.84323 | NA | -0.84323 |
| LRRC71 | NA | NA | -0.84317 | -0.84317 |
| KCTD19 | NA | -0.84265 | NA | -0.84265 |
| ALX4 | NA | -0.84258 | NA | -0.84258 |
| BCO2 | NA | -0.84254 | NA | -0.84254 |
| BRINP1 | NA | -0.84214 | NA | -0.84214 |
| CCDC64B | NA | -0.84212 | NA | -0.84212 |
| CLEC4F | NA | -0.84212 | NA | -0.84212 |
| DNAJC25 | NA | -0.84205 | NA | -0.84205 |
| CHDH | NA | -0.8414 | NA | -0.8414 |
| CCDC180 | NA | -0.84074 | NA | -0.84074 |
| E2F4 | NA | -0.84037 | NA | -0.84037 |

| | | | | |
|---|---|---|---|---|
| TUBE1 | NA | -0.84023 | NA | -0.84023 |
| BTN3A1 | NA | -0.8401 | NA | -0.8401 |
| DENND4B | NA | -0.84003 | NA | -0.84003 |
| RNF133 | NA | -0.83983 | NA | -0.83983 |
| GOLGA6L4 | NA | NA | -0.83977 | -0.83977 |
| PPIAL4B | NA | -0.83946 | NA | -0.83946 |
| FOXJ3 | NA | -0.83917 | NA | -0.83917 |
| MPP7 | NA | -0.83915 | NA | -0.83915 |
| GRIFIN | NA | -0.83889 | NA | -0.83889 |
| MIEF1 | NA | -0.83829 | NA | -0.83829 |
| MAPK12 | NA | -0.838 | NA | -0.838 |
| PCNT | NA | -0.83791 | NA | -0.83791 |
| ENOPH1 | NA | -0.83782 | NA | -0.83782 |
| ONECUT3 | NA | -0.8375 | NA | -0.8375 |
| PEX1 | NA | -0.83736 | NA | -0.83736 |
| RGS7BP | NA | -0.83688 | NA | -0.83688 |
| OLA1 | NA | -0.83617 | NA | -0.83617 |
| NRIP3 | NA | -0.83606 | NA | -0.83606 |
| SCEL | NA | -0.83591 | NA | -0.83591 |
| FAM98C | NA | -0.83588 | NA | -0.83588 |
| USP17L26 | NA | -0.83533 | NA | -0.83533 |
| CPAMD8 | NA | -0.83512 | NA | -0.83512 |
| CNTRL | NA | -0.83476 | NA | -0.83476 |
| FAS | NA | -0.83459 | NA | -0.83459 |
| BTNL8 | NA | -0.83452 | NA | -0.83452 |
| CLIC4 | NA | -0.83408 | NA | -0.83408 |
| THUMPD3 | NA | -0.83405 | NA | -0.83405 |
| SLC7A9 | NA | -0.83403 | NA | -0.83403 |
| XRCC6BP1 | NA | -0.83399 | NA | -0.83399 |
| KNOP1 | NA | -0.83381 | NA | -0.83381 |
| MCM8 | NA | -0.83346 | NA | -0.83346 |
| DEPTOR | NA | -0.83341 | NA | -0.83341 |
| FUT10 | NA | -0.83305 | NA | -0.83305 |
| ZNF423 | NA | -0.8327 | NA | -0.8327 |
| ACVR2A | NA | -0.83266 | NA | -0.83266 |
| MTMR1 | NA | -0.83166 | NA | -0.83166 |
| NPHP1 | NA | -0.83155 | NA | -0.83155 |
| NMT2 | NA | -0.83147 | NA | -0.83147 |
| TNNC2 | NA | -0.8314 | NA | -0.8314 |
| OLFM3 | NA | -0.83114 | NA | -0.83114 |
| MC5R | NA | -0.83048 | NA | -0.83048 |
| PHLDB2 | NA | -0.83017 | NA | -0.83017 |
| NDUFV3 | NA | -0.83011 | NA | -0.83011 |
| ZNF587B | NA | -0.83003 | NA | -0.83003 |
| TTC37 | NA | -0.82999 | NA | -0.82999 |

| | | | | |
|---|---|---|---|---|
| IPMK | NA | -0.82996 | NA | -0.82996 |
| KCNK7 | NA | -0.82976 | NA | -0.82976 |
| AUTS2 | NA | -0.82966 | NA | -0.82966 |
| MED16 | NA | -0.82952 | NA | -0.82952 |
| ALOXE3 | NA | -0.82946 | NA | -0.82946 |
| EXPH5 | NA | -0.82932 | NA | -0.82932 |
| SIN3B | NA | -0.8289 | NA | -0.8289 |
| ANKRD32 | NA | -0.82885 | NA | -0.82885 |
| MAB21L3 | NA | -0.82876 | NA | -0.82876 |
| RIN1 | NA | -0.82875 | NA | -0.82875 |
| PEA15 | NA | -0.82859 | NA | -0.82859 |
| ERCC6L | NA | -0.82846 | NA | -0.82846 |
| MAP2K5 | NA | -0.82822 | NA | -0.82822 |
| PML | NA | -0.82804 | NA | -0.82804 |
| PDCD1 | NA | -0.82794 | NA | -0.82794 |
| WNT4 | NA | -0.8278 | NA | -0.8278 |
| GPR45 | NA | -0.82779 | NA | -0.82779 |
| BRSK1 | NA | -0.8277 | NA | -0.8277 |
| FSBP | NA | -0.82714 | NA | -0.82714 |
| ADORA3 | NA | -0.82696 | NA | -0.82696 |
| NIPA1 | NA | -0.82677 | NA | -0.82677 |
| ZFP90 | NA | -0.82612 | NA | -0.82612 |
| ABCA9 | NA | -0.82586 | NA | -0.82586 |
| UNC119 | NA | -0.82576 | NA | -0.82576 |
| ZNF622 | NA | -0.82544 | NA | -0.82544 |
| TRIOBP | NA | -0.82533 | NA | -0.82533 |
| MMEL1 | NA | NA | -0.82522 | -0.82522 |
| CYB5R4 | NA | -0.82521 | NA | -0.82521 |
| PPOX | NA | -0.82519 | NA | -0.82519 |
| HECTD1 | NA | NA | -0.82512 | -0.82512 |
| FAM189A2 | NA | -0.82507 | NA | -0.82507 |
| KDM5B | NA | -0.82506 | NA | -0.82506 |
| NIM1K | NA | -0.82494 | NA | -0.82494 |
| SURF1 | NA | -0.82458 | NA | -0.82458 |
| LY96 | NA | -0.82437 | NA | -0.82437 |
| AGR2 | NA | -0.82435 | NA | -0.82435 |
| TXLNG | NA | -0.82383 | NA | -0.82383 |
| TBX2 | NA | -0.82351 | NA | -0.82351 |
| USP17L15 | NA | -0.82333 | NA | -0.82333 |
| ARRDC5 | NA | -0.82303 | NA | -0.82303 |
| NBPF8 | NA | -0.82299 | NA | -0.82299 |
| ATAD3B | NA | -0.82284 | NA | -0.82284 |
| CYP2E1 | NA | -0.82162 | NA | -0.82162 |
| NDUFS3 | NA | -0.82159 | NA | -0.82159 |
| YME1L1 | NA | -0.82142 | NA | -0.82142 |

| | | | | |
|---|---|---|---|---|
| LYPLA1 | NA | -0.82138 | NA | -0.82138 |
| DIO2 | NA | -0.8212 | NA | -0.8212 |
| TPRG1 | NA | -0.82103 | NA | -0.82103 |
| CYP2B6 | NA | -0.82065 | NA | -0.82065 |
| FZD8 | NA | -0.82043 | NA | -0.82043 |
| TEX33 | NA | -0.82038 | NA | -0.82038 |
| UBE2G1 | NA | -0.82014 | NA | -0.82014 |
| PINX1 | NA | -0.81998 | NA | -0.81998 |
| EHD1 | NA | -0.81988 | NA | -0.81988 |
| F8A3 | NA | NA | -0.81981 | -0.81981 |
| ZNF277 | NA | -0.81959 | NA | -0.81959 |
| GAN | NA | -0.8194 | NA | -0.8194 |
| WBP4 | NA | -0.81894 | NA | -0.81894 |
| ATP5L2 | NA | -0.8185 | NA | -0.8185 |
| TBC1D9B | NA | -0.81811 | NA | -0.81811 |
| SLC4A9 | NA | -0.81807 | NA | -0.81807 |
| ZFR2 | NA | -0.81801 | NA | -0.81801 |
| CPB1 | NA | -0.81789 | NA | -0.81789 |
| CD6 | NA | -0.81761 | NA | -0.81761 |
| GADD45G | NA | NA | -0.81744 | -0.81744 |
| FAM126A | NA | -0.81736 | NA | -0.81736 |
| CABP2 | NA | -0.81731 | NA | -0.81731 |
| CPN1 | NA | -0.81663 | NA | -0.81663 |
| DNMBP | NA | -0.81648 | NA | -0.81648 |
| MKL1 | NA | -0.81616 | NA | -0.81616 |
| CIB4 | NA | -0.81613 | NA | -0.81613 |
| COG5 | NA | NA | -0.81557 | -0.81557 |
| DCDC2 | NA | -0.81556 | NA | -0.81556 |
| IL9R | NA | -0.81552 | NA | -0.81552 |
| BRINP2 | NA | -0.81527 | NA | -0.81527 |
| ISM2 | NA | -0.81473 | NA | -0.81473 |
| PDE7A | NA | -0.81452 | NA | -0.81452 |
| TMEM242 | NA | -0.81446 | NA | -0.81446 |
| HDAC7 | NA | -0.81418 | NA | -0.81418 |
| SLX4 | NA | NA | -0.81406 | -0.81406 |
| AKR1C1 | NA | -0.81342 | NA | -0.81342 |
| ZNF789 | NA | -0.81308 | NA | -0.81308 |
| PTP4A2 | NA | NA | -0.81304 | -0.81304 |
| AIM2 | NA | -0.81275 | NA | -0.81275 |
| PGR | NA | -0.81275 | NA | -0.81275 |
| DDX39A | NA | -0.81245 | NA | -0.81245 |
| GOLGA6L2 | NA | -0.81241 | NA | -0.81241 |
| MRGPRE | NA | -0.81238 | NA | -0.81238 |
| WDR48 | NA | -0.81234 | NA | -0.81234 |
| PIWIL4 | NA | -0.81222 | NA | -0.81222 |

| | | | | |
|---|---|---|---|---|
| BAG4 | NA | -0.81168 | NA | -0.81168 |
| RIBC2 | NA | -0.81124 | NA | -0.81124 |
| VSIG1 | NA | -0.81071 | NA | -0.81071 |
| COX14 | NA | -0.81021 | NA | -0.81021 |
| GPR114 | NA | -0.81021 | NA | -0.81021 |
| PLOD2 | NA | -0.81 | NA | -0.81 |
| CPNE6 | NA | NA | -0.80986 | -0.80986 |
| SPRY4 | NA | -0.8098 | NA | -0.8098 |
| ZSWIM8 | NA | -0.80953 | NA | -0.80953 |
| RALBP1 | NA | -0.80949 | NA | -0.80949 |
| MRGPRX2 | NA | -0.80921 | NA | -0.80921 |
| LNX2 | NA | -0.80881 | NA | -0.80881 |
| ARL14EP | NA | -0.80866 | NA | -0.80866 |
| REM1 | NA | -0.80853 | NA | -0.80853 |
| HOXD12 | NA | -0.80775 | NA | -0.80775 |
| CCDC58 | NA | -0.80722 | NA | -0.80722 |
| SOWAHB | NA | -0.8067 | NA | -0.8067 |
| FUT7 | NA | -0.80666 | NA | -0.80666 |
| ZCCHC5 | NA | -0.8065 | NA | -0.8065 |
| NMRK1 | NA | -0.80628 | NA | -0.80628 |
| NGLY1 | NA | NA | -0.80612 | -0.80612 |
| AATK | NA | -0.80607 | NA | -0.80607 |
| SGSH | NA | -0.80592 | NA | -0.80592 |
| VEGFB | NA | -0.80573 | NA | -0.80573 |
| FAM156A | NA | -0.80561 | NA | -0.80561 |
| CNBD2 | NA | -0.80557 | NA | -0.80557 |
| ACOX1 | NA | -0.80555 | NA | -0.80555 |
| PREX2 | NA | -0.80507 | NA | -0.80507 |
| ZNF195 | NA | -0.80451 | NA | -0.80451 |
| ELMO3 | NA | -0.8044 | NA | -0.8044 |
| FLNA | NA | -0.80438 | NA | -0.80438 |
| RARG | NA | -0.80276 | NA | -0.80276 |
| ATP2A1 | NA | -0.80272 | NA | -0.80272 |
| FOXL2NB | NA | -0.80255 | NA | -0.80255 |
| STRA13 | NA | -0.80245 | NA | -0.80245 |
| KRT80 | NA | -0.80237 | NA | -0.80237 |
| ZHX2 | NA | -0.8019 | NA | -0.8019 |
| TMEM208 | NA | -0.80174 | NA | -0.80174 |
| IBA57 | NA | NA | -0.80172 | -0.80172 |
| EPHB1 | NA | -0.80128 | NA | -0.80128 |
| ANKRD60 | NA | -0.8006 | NA | -0.8006 |
| WFDC12 | NA | -0.80044 | NA | -0.80044 |
| CT47A6 | NA | NA | -0.8004 | -0.8004 |
| AKNAD1 | NA | -0.80032 | NA | -0.80032 |
| SVOP | NA | -0.7996 | NA | -0.7996 |

| | | | | |
|---|---|---|---|---|
| ECE1 | NA | -0.79951 | NA | -0.79951 |
| HAAO | NA | -0.79932 | NA | -0.79932 |
| GEN1 | NA | -0.79886 | NA | -0.79886 |
| TMX3 | NA | -0.79856 | NA | -0.79856 |
| ZBTB12 | NA | -0.79849 | NA | -0.79849 |
| DNAAF2 | NA | -0.79831 | NA | -0.79831 |
| PHKG2 | NA | -0.79819 | NA | -0.79819 |
| LRRC27 | NA | -0.79785 | NA | -0.79785 |
| GMPR2 | NA | -0.79779 | NA | -0.79779 |
| TUBA4A | NA | -0.79753 | NA | -0.79753 |
| IKBKB | NA | -0.79728 | NA | -0.79728 |
| DCAF4 | NA | -0.79717 | NA | -0.79717 |
| RNF216 | NA | -0.79677 | NA | -0.79677 |
| EIF5AL1 | NA | -0.79673 | NA | -0.79673 |
| RPS6KC1 | NA | -0.79673 | NA | -0.79673 |
| AMIGO1 | NA | -0.79583 | NA | -0.79583 |
| CNOT11 | NA | -0.7956 | NA | -0.7956 |
| ZNF92 | NA | -0.79553 | NA | -0.79553 |
| VWC2L | NA | -0.79503 | NA | -0.79503 |
| GCNT1 | NA | -0.79497 | NA | -0.79497 |
| KLHL12 | NA | -0.79489 | NA | -0.79489 |
| SYTL2 | NA | -0.79463 | NA | -0.79463 |
| LRRC42 | NA | -0.7945 | NA | -0.7945 |
| BHLHA15 | NA | -0.794 | NA | -0.794 |
| ZNF510 | NA | -0.79375 | NA | -0.79375 |
| PENK | NA | -0.79323 | NA | -0.79323 |
| TAX1BP1 | NA | -0.79309 | NA | -0.79309 |
| OR11H6 | NA | -0.79248 | NA | -0.79248 |
| MANSC1 | NA | -0.7921 | NA | -0.7921 |
| POTEC | NA | NA | -0.79134 | -0.79134 |
| MAPT | NA | -0.79126 | NA | -0.79126 |
| TIGD6 | NA | -0.79106 | NA | -0.79106 |
| NOV | NA | -0.79072 | NA | -0.79072 |
| SPARCL1 | NA | -0.79068 | NA | -0.79068 |
| JMJD4 | NA | -0.79032 | NA | -0.79032 |
| ERCC6 | NA | -0.79001 | NA | -0.79001 |
| FSTL1 | NA | -0.78999 | NA | -0.78999 |
| TEAD1 | NA | -0.78996 | NA | -0.78996 |
| HIPK4 | NA | -0.78974 | NA | -0.78974 |
| BSG | NA | -0.78958 | NA | -0.78958 |
| ZBTB16 | NA | -0.78914 | NA | -0.78914 |
| TMEM141 | NA | -0.78898 | NA | -0.78898 |
| SCNN1A | NA | -0.78888 | NA | -0.78888 |
| OR2B3 | NA | -0.78883 | NA | -0.78883 |
| IL2RB | NA | -0.78861 | NA | -0.78861 |

| | | | | |
|---|---|---|---|---|
| TDRD6 | NA | -0.78855 | NA | -0.78855 |
| PPP4R4 | NA | -0.78838 | NA | -0.78838 |
| USP17L12 | NA | -0.78835 | NA | -0.78835 |
| SLC7A6 | NA | -0.7882 | NA | -0.7882 |
| CBWD6 | NA | -0.78775 | NA | -0.78775 |
| CCDC142 | NA | -0.78735 | NA | -0.78735 |
| ARMC4 | NA | -0.78714 | NA | -0.78714 |
| LRIG1 | NA | -0.78707 | NA | -0.78707 |
| AMER3 | NA | -0.78683 | NA | -0.78683 |
| RFWD2 | NA | NA | -0.78652 | -0.78652 |
| GAGE4 | NA | -0.78637 | NA | -0.78637 |
| PTK6 | NA | -0.78571 | NA | -0.78571 |
| CFP | NA | -0.78557 | NA | -0.78557 |
| CMTR2 | NA | -0.78529 | NA | -0.78529 |
| NKRF | NA | -0.78475 | NA | -0.78475 |
| MFSD12 | NA | -0.78441 | NA | -0.78441 |
| PGPEP1 | NA | -0.78415 | NA | -0.78415 |
| PDS5A | NA | -0.78377 | NA | -0.78377 |
| LIN9 | NA | -0.78374 | NA | -0.78374 |
| OR5I1 | NA | -0.78365 | NA | -0.78365 |
| ZNF74 | NA | -0.78337 | NA | -0.78337 |
| KLHDC4 | NA | -0.78327 | NA | -0.78327 |
| EPX | NA | -0.78263 | NA | -0.78263 |
| TM7SF2 | NA | -0.78245 | NA | -0.78245 |
| HDAC4 | NA | -0.7823 | NA | -0.7823 |
| CPNE1 | NA | -0.78202 | NA | -0.78202 |
| CCL18 | NA | -0.78165 | NA | -0.78165 |
| CASP8AP2 | NA | -0.78152 | NA | -0.78152 |
| IDH1 | NA | -0.78123 | NA | -0.78123 |
| SPZ1 | NA | -0.78041 | NA | -0.78041 |
| SIRPB2 | NA | -0.78024 | NA | -0.78024 |
| ATP8B2 | NA | -0.78017 | NA | -0.78017 |
| CCDC106 | NA | -0.78006 | NA | -0.78006 |
| TUBA3C | NA | -0.77965 | NA | -0.77965 |
| EGF | NA | -0.77954 | NA | -0.77954 |
| UTS2R | NA | -0.77953 | NA | -0.77953 |
| SDF4 | NA | -0.7795 | NA | -0.7795 |
| CD209 | NA | -0.77944 | NA | -0.77944 |
| METTL15 | NA | -0.77938 | NA | -0.77938 |
| SH3GLB1 | NA | -0.77937 | NA | -0.77937 |
| PGLYRP3 | NA | -0.779 | NA | -0.779 |
| STUB1 | NA | -0.77885 | NA | -0.77885 |
| PRSS45 | NA | -0.77882 | NA | -0.77882 |
| KLHL38 | NA | -0.7788 | NA | -0.7788 |
| UPK3BL | NA | -0.7778 | NA | -0.7778 |

| | | | | |
|---|---|---|---|---|
| FXYD5 | NA | -0.77759 | NA | -0.77759 |
| KDM5A | NA | -0.77743 | NA | -0.77743 |
| EXOC3L4 | NA | -0.77741 | NA | -0.77741 |
| TRIM25 | NA | -0.77718 | NA | -0.77718 |
| NCCRP1 | NA | -0.77712 | NA | -0.77712 |
| OR2A7 | NA | -0.77645 | NA | -0.77645 |
| UFC1 | NA | -0.77645 | NA | -0.77645 |
| FOXD4L4 | NA | NA | -0.77633 | -0.77633 |
| CRACR2B | NA | -0.77618 | NA | -0.77618 |
| PHOSPHO1 | NA | -0.77598 | NA | -0.77598 |
| ASXL2 | NA | -0.77595 | NA | -0.77595 |
| ARSB | NA | -0.77588 | NA | -0.77588 |
| TMEM52 | NA | -0.77558 | NA | -0.77558 |
| TCF21 | NA | -0.77545 | NA | -0.77545 |
| TRIB1 | NA | -0.7754 | NA | -0.7754 |
| TRIM77 | NA | -0.77531 | NA | -0.77531 |
| DYRK1A | NA | -0.77523 | NA | -0.77523 |
| CARD11 | NA | -0.77504 | NA | -0.77504 |
| ZNF322 | NA | NA | -0.77489 | -0.77489 |
| FXR1 | NA | -0.77486 | NA | -0.77486 |
| TSPAN2 | NA | -0.77484 | NA | -0.77484 |
| ZNF772 | NA | -0.77472 | NA | -0.77472 |
| DLGAP5 | NA | -0.77462 | NA | -0.77462 |
| OR6C6 | NA | -0.77457 | NA | -0.77457 |
| PPP3CC | NA | -0.77421 | NA | -0.77421 |
| CASC5 | NA | -0.77406 | NA | -0.77406 |
| CCDC148 | NA | -0.77379 | NA | -0.77379 |
| SYT13 | NA | -0.77377 | NA | -0.77377 |
| PPP3R2 | NA | -0.77349 | NA | -0.77349 |
| SAMD13 | NA | -0.77346 | NA | -0.77346 |
| LARP7 | NA | -0.77342 | NA | -0.77342 |
| ENDOD1 | NA | -0.77319 | NA | -0.77319 |
| FAM72A | NA | NA | -0.77281 | -0.77281 |
| BCAP31 | NA | -0.77275 | NA | -0.77275 |
| PDLIM7 | NA | NA | -0.77269 | -0.77269 |
| MFSD4 | NA | -0.77236 | NA | -0.77236 |
| SOD3 | NA | -0.77187 | NA | -0.77187 |
| ZDHHC19 | NA | -0.77178 | NA | -0.77178 |
| WDR34 | NA | -0.77137 | NA | -0.77137 |
| GTF2IRD2B | NA | -0.7713 | NA | -0.7713 |
| GTPBP3 | NA | -0.77108 | NA | -0.77108 |
| EED | NA | -0.77107 | NA | -0.77107 |
| CUTC | NA | -0.7708 | NA | -0.7708 |
| TYW3 | NA | -0.77073 | NA | -0.77073 |
| SELL | NA | -0.77061 | NA | -0.77061 |

| | | | | |
|---|---|---|---|---|
| SLC25A19 | NA | -0.77 | NA | -0.77 |
| COL21A1 | NA | -0.7696 | NA | -0.7696 |
| FRMPD2 | NA | -0.76945 | NA | -0.76945 |
| PAK4 | NA | -0.76807 | NA | -0.76807 |
| ZBTB8A | NA | -0.76799 | NA | -0.76799 |
| GDF7 | NA | -0.76798 | NA | -0.76798 |
| TP53TG3D | NA | -0.76784 | NA | -0.76784 |
| LYPLA2 | NA | -0.76747 | NA | -0.76747 |
| ACAP1 | NA | -0.76742 | NA | -0.76742 |
| LTB | NA | -0.76738 | NA | -0.76738 |
| SGOL1 | NA | -0.76735 | NA | -0.76735 |
| TMCO6 | NA | -0.76698 | NA | -0.76698 |
| IFT22 | NA | -0.76697 | NA | -0.76697 |
| PRMT7 | NA | -0.76678 | NA | -0.76678 |
| ADAM18 | NA | -0.76676 | NA | -0.76676 |
| PLEKHB1 | NA | -0.76665 | NA | -0.76665 |
| HK3 | NA | -0.76663 | NA | -0.76663 |
| DHCR24 | NA | -0.76646 | NA | -0.76646 |
| DHRS4 | NA | -0.76639 | NA | -0.76639 |
| POC1A | NA | -0.76606 | NA | -0.76606 |
| TAS2R41 | NA | -0.76591 | NA | -0.76591 |
| AMOT | NA | -0.76574 | NA | -0.76574 |
| MYSM1 | NA | -0.76571 | NA | -0.76571 |
| CHPF2 | NA | -0.76556 | NA | -0.76556 |
| DUSP10 | NA | -0.76546 | NA | -0.76546 |
| TRIM67 | NA | -0.76542 | NA | -0.76542 |
| ACKR1 | NA | -0.7652 | NA | -0.7652 |
| HIST1H4D | NA | -0.76491 | NA | -0.76491 |
| ZNF846 | NA | -0.76441 | NA | -0.76441 |
| PKHD1 | NA | -0.76431 | NA | -0.76431 |
| HEATR5B | NA | -0.76423 | NA | -0.76423 |
| TBC1D2B | NA | -0.76329 | NA | -0.76329 |
| KHSRP | NA | -0.76327 | NA | -0.76327 |
| PNP | NA | -0.76307 | NA | -0.76307 |
| SREBF2 | NA | -0.76305 | NA | -0.76305 |
| HNF1A | NA | -0.76302 | NA | -0.76302 |
| OTULIN | NA | -0.76291 | NA | -0.76291 |
| WNT5A | NA | -0.76289 | NA | -0.76289 |
| DRD1 | NA | -0.76243 | NA | -0.76243 |
| COMP | NA | -0.76236 | NA | -0.76236 |
| CCER1 | NA | -0.76221 | NA | -0.76221 |
| YTHDF2 | NA | NA | -0.7622 | -0.7622 |
| MMP2 | NA | -0.76183 | NA | -0.76183 |
| SLX1B | NA | -0.76161 | NA | -0.76161 |
| APOBEC2 | NA | -0.76136 | NA | -0.76136 |

| | | | | |
|---|---|---|---|---|
| OR2K2 | NA | -0.76109 | NA | -0.76109 |
| TRIP12 | NA | -0.76087 | NA | -0.76087 |
| ZSCAN25 | NA | -0.76056 | NA | -0.76056 |
| FAM25A | NA | -0.76018 | NA | -0.76018 |
| ZNF180 | NA | -0.76016 | NA | -0.76016 |
| CPOX | NA | -0.76011 | NA | -0.76011 |
| KDM6B | NA | -0.76007 | NA | -0.76007 |
| CCHCR1 | NA | -0.75979 | NA | -0.75979 |
| ST8SIA3 | NA | -0.75884 | NA | -0.75884 |
| BDNF | NA | -0.75878 | NA | -0.75878 |
| SENP1 | NA | -0.7587 | NA | -0.7587 |
| PLEKHG5 | NA | -0.75864 | NA | -0.75864 |
| POLR1D | NA | -0.75846 | NA | -0.75846 |
| FAM89B | NA | -0.75831 | NA | -0.75831 |
| DLX4 | NA | -0.75824 | NA | -0.75824 |
| WTH3DI | NA | -0.75757 | NA | -0.75757 |
| LAMA1 | NA | -0.75738 | NA | -0.75738 |
| TRIM51 | NA | -0.75733 | NA | -0.75733 |
| PDCD10 | NA | NA | -0.75708 | -0.75708 |
| ATG14 | NA | NA | -0.75699 | -0.75699 |
| ANKRD44 | NA | -0.75691 | NA | -0.75691 |
| SLC23A3 | NA | -0.75664 | NA | -0.75664 |
| STAG2 | NA | NA | -0.75613 | -0.75613 |
| RFX3 | NA | -0.75571 | NA | -0.75571 |
| HFE2 | NA | -0.75554 | NA | -0.75554 |
| STRADA | NA | -0.75545 | NA | -0.75545 |
| CTSV | NA | -0.75538 | NA | -0.75538 |
| RBCK1 | NA | -0.75445 | NA | -0.75445 |
| HHLA1 | NA | -0.75411 | NA | -0.75411 |
| MTCH2 | NA | -0.7541 | NA | -0.7541 |
| TTBK1 | NA | -0.75357 | NA | -0.75357 |
| SLMO2 | NA | -0.75333 | NA | -0.75333 |
| USP17L11 | NA | -0.75333 | NA | -0.75333 |
| CNNM1 | NA | -0.75298 | NA | -0.75298 |
| PRAMEF6 | NA | -0.75282 | NA | -0.75282 |
| NSD1 | NA | -0.75263 | NA | -0.75263 |
| OTUD7A | NA | -0.75228 | NA | -0.75228 |
| CT45A1 | NA | -0.75218 | NA | -0.75218 |
| PPTC7 | NA | -0.75209 | NA | -0.75209 |
| MMP1 | NA | -0.75206 | NA | -0.75206 |
| 4-Sep | NA | -0.75202 | NA | -0.75202 |
| FER1L5 | NA | -0.75169 | NA | -0.75169 |
| GTSCR1 | NA | -0.75138 | NA | -0.75138 |
| INCA1 | NA | -0.75134 | NA | -0.75134 |
| TAS2R31 | NA | -0.75075 | NA | -0.75075 |

| | | | | |
|---|---|---|---|---|
| ZNF284 | NA | -0.75075 | NA | -0.75075 |
| FAM46D | NA | -0.75048 | NA | -0.75048 |
| USH1C | NA | -0.75048 | NA | -0.75048 |
| IQCF2 | NA | -0.75023 | NA | -0.75023 |
| WHAMM | NA | -0.74979 | NA | -0.74979 |
| COMMD7 | NA | -0.74965 | NA | -0.74965 |
| STK32C | NA | -0.749 | NA | -0.749 |
| COL4A3BP | NA | -0.74859 | NA | -0.74859 |
| ETNPPL | NA | -0.74783 | NA | -0.74783 |
| CD320 | NA | -0.74762 | NA | -0.74762 |
| SULT4A1 | NA | -0.74729 | NA | -0.74729 |
| ALPP | NA | -0.74678 | NA | -0.74678 |
| MT1M | NA | -0.74666 | NA | -0.74666 |
| NPIPB9 | NA | -0.74636 | NA | -0.74636 |
| RAD51AP2 | NA | -0.74564 | NA | -0.74564 |
| ZFP57 | NA | -0.74562 | NA | -0.74562 |
| NTN5 | NA | -0.74548 | NA | -0.74548 |
| TMEM47 | NA | -0.74524 | NA | -0.74524 |
| ENTPD1 | NA | -0.74508 | NA | -0.74508 |
| NUDT8 | NA | -0.74501 | NA | -0.74501 |
| IGSF6 | NA | -0.74495 | NA | -0.74495 |
| TEDDM1 | NA | -0.7449 | NA | -0.7449 |
| TIMM17B | NA | -0.74489 | NA | -0.74489 |
| ZNF540 | NA | -0.74485 | NA | -0.74485 |
| ALG9 | NA | -0.74482 | NA | -0.74482 |
| PHKB | NA | -0.74469 | NA | -0.74469 |
| C5AR2 | NA | -0.74435 | NA | -0.74435 |
| OTUD4 | NA | -0.7442 | NA | -0.7442 |
| TAF1L | NA | -0.74411 | NA | -0.74411 |
| INPP4A | NA | -0.74374 | NA | -0.74374 |
| POLR2J3 | NA | -0.74337 | NA | -0.74337 |
| PBXIP1 | NA | -0.74314 | NA | -0.74314 |
| TRMT1L | NA | -0.74308 | NA | -0.74308 |
| RAVER1 | NA | -0.74305 | NA | -0.74305 |
| OR10K1 | NA | -0.74293 | NA | -0.74293 |
| COQ3 | NA | -0.7429 | NA | -0.7429 |
| AGO1 | NA | -0.74289 | NA | -0.74289 |
| CT45A2 | NA | -0.74286 | NA | -0.74286 |
| NARR | NA | -0.74281 | NA | -0.74281 |
| CTSK | NA | -0.74269 | NA | -0.74269 |
| NFAT5 | NA | -0.74246 | NA | -0.74246 |
| IFI35 | NA | -0.74214 | NA | -0.74214 |
| PPT1 | NA | -0.74204 | NA | -0.74204 |
| TOM1L1 | NA | -0.74191 | NA | -0.74191 |
| DACH1 | NA | -0.7419 | NA | -0.7419 |

| | | | | |
|---|---|---|---|---|
| VWA2 | NA | -0.7417 | NA | -0.7417 |
| PLBD1 | NA | -0.74165 | NA | -0.74165 |
| FAM13C | NA | -0.74123 | NA | -0.74123 |
| ZNF98 | NA | -0.74109 | NA | -0.74109 |
| KIF15 | NA | -0.74102 | NA | -0.74102 |
| PPP2R5D | NA | -0.74099 | NA | -0.74099 |
| GIT1 | NA | -0.74087 | NA | -0.74087 |
| BOLA3 | NA | -0.74086 | NA | -0.74086 |
| SCAMP5 | NA | -0.74049 | NA | -0.74049 |
| SLC25A32 | NA | -0.74019 | NA | -0.74019 |
| ATXN7L2 | NA | NA | -0.73965 | -0.73965 |
| PRAMEF5 | NA | NA | -0.73231 | -0.73231 |
| KRT33A | NA | NA | -0.7309 | -0.7309 |
| SLC16A8 | NA | NA | -0.72961 | -0.72961 |
| RPS10-NUDT3 | NA | NA | -0.72724 | -0.72724 |
| CHST13 | NA | NA | -0.72017 | -0.72017 |
| SFT2D2 | NA | NA | -0.72005 | -0.72005 |
| C14orf166 | NA | NA | -0.71891 | -0.71891 |
| TRAPPC2L | NA | NA | -0.71625 | -0.71625 |
| PGM3 | NA | NA | -0.71468 | -0.71468 |
| PRRC2B | NA | NA | -0.71379 | -0.71379 |
| DUFC2-KCTD1 | NA | NA | -0.71328 | -0.71328 |
| SLC9A5 | NA | NA | -0.7132 | -0.7132 |
| RER1 | NA | NA | -0.70942 | -0.70942 |
| SREK1 | NA | NA | -0.70623 | -0.70623 |
| REPIN1 | NA | NA | -0.70613 | -0.70613 |
| ECM1 | NA | NA | -0.69756 | -0.69756 |
| NPIPB3 | NA | NA | -0.69624 | -0.69624 |
| C10orf2 | NA | NA | -0.69498 | -0.69498 |
| LDHB | NA | NA | -0.69466 | -0.69466 |
| EEFSEC | NA | NA | -0.69437 | -0.69437 |
| DCAF7 | NA | NA | -0.69387 | -0.69387 |
| CEACAM6 | NA | NA | -0.69315 | -0.69315 |
| CKS1B | NA | NA | -0.69269 | -0.69269 |
| EEF1A2 | NA | NA | -0.692 | -0.692 |
| SERPINB1 | NA | NA | -0.69139 | -0.69139 |
| OR11H4 | NA | NA | -0.68931 | -0.68931 |
| ATP5E | NA | NA | -0.68899 | -0.68899 |
| OST4 | NA | NA | -0.68512 | -0.68512 |
| MAPKAP1 | NA | NA | -0.68429 | -0.68429 |
| TMEM41B | NA | NA | -0.68277 | -0.68277 |
| MSN | NA | NA | -0.68182 | -0.68182 |
| NEK2 | NA | NA | -0.68098 | -0.68098 |
| TMED7 | NA | NA | -0.68073 | -0.68073 |
| TBC1D3 | NA | NA | -0.67886 | -0.67886 |

| | | | | |
|---|---|---|---|---|
| RBM26 | NA | NA | -0.67825 | -0.67825 |
| ABLIM2 | NA | NA | -0.67753 | -0.67753 |
| ARF1 | NA | NA | -0.677 | -0.677 |
| TPM3 | NA | NA | -0.67646 | -0.67646 |
| LRRC57 | NA | NA | -0.67529 | -0.67529 |
| FAM35A | NA | NA | -0.67296 | -0.67296 |
| GATA2 | NA | NA | -0.6719 | -0.6719 |
| DIO3 | NA | NA | -0.67044 | -0.67044 |
| GOLGA6L10 | NA | NA | -0.66598 | -0.66598 |
| FOXD4L1 | NA | NA | -0.66461 | -0.66461 |
| TM9SF3 | NA | NA | -0.66222 | -0.66222 |
| STK40 | NA | NA | -0.65507 | -0.65507 |
| TEX22 | NA | NA | -0.65228 | -0.65228 |
| DENND2A | NA | NA | -0.65058 | -0.65058 |
| SYNC | NA | NA | -0.64503 | -0.64503 |
| HIST1H2AI | NA | NA | -0.64468 | -0.64468 |
| DPYSL3 | NA | NA | -0.6393 | -0.6393 |
| COLCA2 | NA | NA | -0.63669 | -0.63669 |
| ST14 | NA | NA | -0.63346 | -0.63346 |
| FCGR1A | NA | NA | -0.62932 | -0.62932 |
| FNDC3B | NA | NA | -0.62739 | -0.62739 |
| ACTA1 | NA | NA | -0.62404 | -0.62404 |
| C8orf33 | NA | NA | -0.62347 | -0.62347 |
| SAFB | NA | NA | -0.61782 | -0.61782 |
| MPHOSPH8 | NA | NA | -0.61715 | -0.61715 |
| SEPHS2 | NA | NA | -0.61682 | -0.61682 |
| SMAD4 | NA | NA | -0.61665 | -0.61665 |
| GZMB | NA | NA | -0.61506 | -0.61506 |
| MICALL2 | NA | NA | -0.61415 | -0.61415 |
| APH1A | NA | NA | -0.61107 | -0.61107 |
| GOLGA6L9 | NA | NA | -0.61042 | -0.61042 |
| MOB3C | NA | NA | -0.60864 | -0.60864 |
| C20orf196 | NA | NA | -0.60776 | -0.60776 |
| RASL12 | NA | NA | -0.60656 | -0.60656 |
| SHISA6 | NA | NA | -0.60611 | -0.60611 |
| MAFK | NA | NA | -0.60571 | -0.60571 |
| ZNF213 | NA | NA | -0.60548 | -0.60548 |
| HAVCR1 | NA | NA | -0.60309 | -0.60309 |
| LPCAT1 | NA | NA | -0.60174 | -0.60174 |
| SPDYE2B | NA | NA | -0.59966 | -0.59966 |
| KCNQ4 | NA | NA | -0.59875 | -0.59875 |
| C16orf47 | NA | NA | -0.59854 | -0.59854 |
| CTDSPL2 | NA | NA | -0.59846 | -0.59846 |
| CXorf38 | NA | NA | -0.59834 | -0.59834 |
| SPATA2 | NA | NA | -0.59593 | -0.59593 |

| | | | | |
|---|---|---|---|---|
| PPME1 | NA | NA | -0.5957 | -0.5957 |
| ARHGAP21 | NA | NA | -0.59396 | -0.59396 |
| C3orf38 | NA | NA | -0.59306 | -0.59306 |
| GPR3 | NA | NA | -0.59246 | -0.59246 |
| TBC1D3F | NA | NA | -0.59217 | -0.59217 |
| ATP2C1 | NA | NA | -0.59062 | -0.59062 |
| HSD17B7 | NA | NA | -0.58984 | -0.58984 |
| TRIM49D2 | NA | NA | -0.5895 | -0.5895 |
| TRPM5 | NA | NA | -0.58892 | -0.58892 |
| GRHL3 | NA | NA | -0.58841 | -0.58841 |
| MAPKBP1 | NA | NA | -0.58777 | -0.58777 |
| PPIAL4A | NA | NA | -0.58773 | -0.58773 |
| HEXIM1 | NA | NA | -0.58309 | -0.58309 |
| ATP5A1 | NA | NA | -0.58246 | -0.58246 |
| CTCFL | NA | NA | -0.58089 | -0.58089 |
| PRCD | NA | NA | -0.57652 | -0.57652 |
| OR2L3 | NA | NA | -0.57639 | -0.57639 |
| CTDSP1 | NA | NA | -0.57621 | -0.57621 |
| FUS | NA | NA | -0.57549 | -0.57549 |
| MOB2 | NA | NA | -0.57504 | -0.57504 |
| MGAT5 | NA | NA | -0.57397 | -0.57397 |
| PSENEN | NA | NA | -0.57389 | -0.57389 |
| HOOK3 | NA | NA | -0.57306 | -0.57306 |
| CA3 | NA | NA | -0.57275 | -0.57275 |
| LCE1B | NA | NA | -0.57264 | -0.57264 |
| DCLRE1C | NA | NA | -0.5691 | -0.5691 |
| GJD4 | NA | NA | -0.56821 | -0.56821 |
| RGS6 | NA | NA | -0.56755 | -0.56755 |
| GPR135 | NA | NA | -0.56644 | -0.56644 |
| ZNF280C | NA | NA | -0.56602 | -0.56602 |
| SEC31A | NA | NA | -0.56581 | -0.56581 |
| FLYWCH1 | NA | NA | -0.56574 | -0.56574 |
| MPI | NA | NA | -0.5657 | -0.5657 |
| ARHGEF33 | NA | NA | -0.56438 | -0.56438 |
| ZNF697 | NA | NA | -0.56412 | -0.56412 |
| CYP11B2 | NA | NA | -0.56332 | -0.56332 |
| MYH4 | NA | NA | -0.56305 | -0.56305 |
| AGTPBP1 | NA | NA | -0.55993 | -0.55993 |
| PRR20E | NA | NA | -0.55902 | -0.55902 |
| PIK3CD | NA | NA | -0.5586 | -0.5586 |
| ABCB8 | NA | NA | -0.55858 | -0.55858 |
| KSR1 | NA | NA | -0.55768 | -0.55768 |
| ZNRF3 | NA | NA | -0.55725 | -0.55725 |
| PAN-P2RY11 | NA | NA | -0.55691 | -0.55691 |
| HP1BP3 | NA | NA | -0.55627 | -0.55627 |

| | | | | |
|---|---|---|---|---|
| LAYN | NA | NA | -0.55431 | -0.55431 |
| SPRR2A | NA | NA | -0.55418 | -0.55418 |
| OR51A7 | NA | NA | -0.55366 | -0.55366 |
| EZH2 | NA | NA | -0.55311 | -0.55311 |
| WDYHV1 | NA | NA | -0.5527 | -0.5527 |
| KIF20B | NA | NA | -0.55187 | -0.55187 |
| NSUN5 | NA | NA | -0.55086 | -0.55086 |
| CHRM2 | NA | NA | -0.55068 | -0.55068 |
| MIEN1 | NA | NA | -0.54908 | -0.54908 |
| SLC7A8 | NA | NA | -0.54783 | -0.54783 |
| SLC45A3 | NA | NA | -0.54688 | -0.54688 |
| RBM7 | NA | NA | -0.54643 | -0.54643 |
| CCDC27 | NA | NA | -0.54544 | -0.54544 |
| TESC | NA | NA | -0.54195 | -0.54195 |
| ZNF99 | NA | NA | -0.54127 | -0.54127 |
| CCL24 | NA | NA | -0.54115 | -0.54115 |
| SEC24A | NA | NA | -0.54088 | -0.54088 |
| KIF2A | NA | NA | -0.54024 | -0.54024 |
| KIR2DS4 | NA | NA | -0.54017 | -0.54017 |
| ANGPTL3 | NA | NA | -0.53781 | -0.53781 |
| NCOR1 | NA | NA | -0.53683 | -0.53683 |
| INPP5K | NA | NA | -0.53658 | -0.53658 |
| YLPM1 | NA | NA | -0.53538 | -0.53538 |
| JTB | NA | NA | -0.53527 | -0.53527 |
| BAHCC1 | NA | NA | -0.53481 | -0.53481 |
| RPRD1B | NA | NA | -0.53309 | -0.53309 |
| RPL10L | NA | NA | -0.53203 | -0.53203 |
| NAA60 | NA | NA | -0.53195 | -0.53195 |
| SCGB3A2 | NA | NA | -0.53026 | -0.53026 |
| SH3GL3 | NA | NA | -0.52946 | -0.52946 |
| AFMID | NA | NA | -0.52837 | -0.52837 |
| GOLGA8K | NA | NA | -0.52741 | -0.52741 |
| OPLAH | NA | NA | -0.52631 | -0.52631 |
| ATG12 | NA | NA | -0.52469 | -0.52469 |
| OGFOD1 | NA | NA | -0.52452 | -0.52452 |
| KRTCAP2 | NA | NA | -0.5242 | -0.5242 |
| DSE | NA | NA | -0.5233 | -0.5233 |
| ZNF672 | NA | NA | -0.52273 | -0.52273 |
| ZNF248 | NA | NA | -0.52239 | -0.52239 |
| ARHGEF7 | NA | NA | -0.52197 | -0.52197 |
| ATP6V0A1 | NA | NA | -0.52143 | -0.52143 |
| PIF1 | NA | NA | -0.52079 | -0.52079 |
| DGAT2 | NA | NA | -0.52015 | -0.52015 |
| SLC39A1 | NA | NA | -0.51997 | -0.51997 |
| ZNRF2 | NA | NA | -0.51919 | -0.51919 |

| | | | | |
|---|---|---|---|---|
| SNX1 | NA | NA | -0.51909 | -0.51909 |
| NRG4 | NA | NA | -0.51874 | -0.51874 |
| NHSL1 | NA | NA | -0.51859 | -0.51859 |
| GRM3 | NA | NA | -0.51779 | -0.51779 |
| FMN1 | NA | NA | -0.51658 | -0.51658 |
| SNX5 | NA | NA | -0.51559 | -0.51559 |
| TNNI2 | NA | NA | -0.51351 | -0.51351 |
| A3GALT2 | NA | NA | -0.51226 | -0.51226 |
| LRRC18 | NA | NA | -0.51211 | -0.51211 |
| OMMD3-BMI | NA | NA | -0.5121 | -0.5121 |
| GAB4 | NA | NA | -0.51203 | -0.51203 |
| SCG2 | NA | NA | -0.51195 | -0.51195 |
| SGSM3 | NA | NA | -0.51167 | -0.51167 |
| SUCLA2 | NA | NA | -0.51002 | -0.51002 |
| ERCC6L2 | NA | NA | -0.51 | -0.51 |
| ZSWIM5 | NA | NA | -0.50986 | -0.50986 |
| PPP6R3 | NA | NA | -0.50971 | -0.50971 |
| DERL1 | NA | NA | -0.50943 | -0.50943 |
| KDM1A | NA | NA | -0.50854 | -0.50854 |
| MEN1 | NA | NA | -0.50702 | -0.50702 |
| ZBTB18 | NA | NA | -0.50698 | -0.50698 |
| VMAC | NA | NA | -0.50528 | -0.50528 |
| ST6GALNAC5 | NA | NA | -0.50421 | -0.50421 |
| UXS1 | NA | NA | -0.50209 | -0.50209 |
| MPST | NA | NA | -0.50146 | -0.50146 |
| SOX2 | NA | NA | -0.50141 | -0.50141 |
| GPRC5C | NA | NA | -0.50025 | -0.50025 |
| ZNF625 | NA | NA | -0.49975 | -0.49975 |
| PLA2G3 | NA | NA | -0.49955 | -0.49955 |
| FBXO16 | NA | NA | -0.49941 | -0.49941 |
| EFS | NA | NA | -0.49927 | -0.49927 |
| KCNIP3 | NA | NA | -0.49925 | -0.49925 |
| SPCS1 | NA | NA | -0.49865 | -0.49865 |
| ALAS1 | NA | NA | -0.49828 | -0.49828 |
| SSTR3 | NA | NA | -0.49798 | -0.49798 |
| SRM | NA | NA | -0.49743 | -0.49743 |
| 1-Mar | NA | NA | -0.41576 | -0.41576 |

**ntified in each DMG line using CRISPR screen**
